# Mid-aged mice rapidly normalize dysglycemia but aggravate obesity-induced hypothalamic and microglial changes upon dietary obesity reversal

**DOI:** 10.1101/2025.02.26.640035

**Authors:** Alon Zemer, Yulia Haim, Alexandra Tsitrina, Vered Chalifa-Caspi, Habib Muallem, Yair Pincu, Uri Yoel, Alon Monsonego, Assaf Rudich

## Abstract

**Objective:** Obesity-induced-dysglycemia and hypothalamic-microgliosis coincide, but whether they remain linked upon obesity reversal, and what is the effect of age, remain unclear. Here we hypothesized that rapid normalization of dysglycemia upon obesity-reversal remains linked to microgliosis resolution, but differs between young and mid-aged mice.

**Methods:** Young (7w) and mid-aged (1y) mice were fed normal chow (NC) or high-fat diet (HFD,8w), then switched to NC (Rev,2w).

**Results:** Compared to young mice, NC-fed mid-aged mice were heavier and weight-stable, gained weight with HFD comparably, and lost less weight in Rev. HFD-induced dysglycemia was less severe in mid-aged compared to young mice, but similarly normalized by obesity-reversal. However, whole-hypothalamus RNA sequencing revealed 2,419 differentially expressed genes (DEGs) in mid-aged mice, ∼4-times more than in young mice, and in both age-groups ∼80% of DEGs obesity-induced changes were aggravated in Rev. Furthermore, compared with young mice, middle-aged mice showed greater obesity-induced microglial cyto-morphological changes in the arcuate nucleus (ARC), which associated with increased p-NFκB-(p-p65) nuclear staining. Only in middle-aged mice obesity-induced microglial changes were aggravated by obesity reversal, with cell volume correlating (Rho(ρ)=0.691, p=0.001) with adipose tissue crown-like-structures.

**Conclusions:** In conclusion, rapid dysglycemia normalization is uncoupled to the resolution of hypothalamic microgliosis, more-so in mid-age.

## Introduction

Investigating the pathophysiology of obesity and its complications has been greatly aided by rodent models. Yet, while in the clinic losing excess weight is the common dealing, preclinical research has focused on the triggers, mechanisms, and consequences of weight gain, leaving the sequence/dynamics of changes during weight-loss poorly characterized. This is an important gap of knowledge, since uncovering which obesity-related alterations reverse before others can reveal, or at least exclude, key causal relationships, according to the temporality principle in revealing causation (in essence, if B precedes A, A cannot be the cause for B). For example, hypothalamic inflammation, characterized by glial cells’ activation, is a rapid response to hyper-nutrition and weight-gain^1^, documented in both rodent obesity models^1–4^, and humans^5^, and preceding obesity-induced glucose intolerance. Given CNS’s documented regulation of peripheral metabolism^6^, hypothalamic inflammation is thereby considered causal to the occurrence of dysglycemia^7–9^. Yet, similar analysis of how these resolve during weight-loss (obesity reversal), vis-a-vis weight change dynamics and resolution of adipose inflammation, remains unclear. In particular, is resolution of hypothalamic inflammation during weight-loss required/causal for normalization of dysglycemia to occur?

An additional level of complexity is how age affects obesity reversal. Here, again, a gap exist: most rodent obesity models utilize young mice, equivalent to human late-teen/young-adulthood, while obesity in humans is particularly high amongst mid-aged humans^10^. Age is a major factor in weight dynamics and in the pathogenesis of obesity and its complications: Weight at 17-25 years still increases, particularly in males, at a higher rate than during young adulthood (25-45y), a phenomenon observed in all baseline BMI percentiles^11^. Furthermore, weight-loss and/or regain may be more harmful in mid-age, due to largely unknown reasons^12,13^. Thus, how age affects weight gain and loss and their impact on health^14,15^, and what is the contribution of age per-se, versus merely the longer duration (in mid-age vs. young) of the obesogenic insult, remain largely unknown.

In the present study, we hypothesized that reversal of obesity-induced hypothalamic changes is required for normalization of obesity-induced dysglycemia, and that mid-age mice would exhibit different reversal dynamics than young mice, even under same duration of obesity insult. Our results demonstrate that, contrary to our hypothesis, obesity-induced transcriptomic and microglial hypothalamic changes do not reverse before normalization of dysglycemia in young mice. Moreover, mid-aged mice exhibit aggravated transcriptional and microglial changes early in the course of obesity reversal, correlating with sustained adipose tissue inflammation.

## Research Design and Methods

### Animals and experimental protocols

The study was approved by the Ben-Gurion University Institutional Animal Care and Use Committee (Protocol IL-69-11-2021) and was conducted according to the Israeli Animal Welfare Act (National Research Council, 1996). Five-weeks-old male C57BL/6 mice were purchased from ENVIGO RMS Ltd. (Jerusalem, Israel). Mice were acclimatized for 2w, with free access to a normal chow (NC) diet (15% of calories from fat, Ssniff Spezialdiäten GmbH) and water. Animals were housed two mice per cage under controlled temperature (22 ± 1°C), in a 12-h light-dark cycles. The dietary protocol for diet-induced obesity and early weight-loss was published previously^16^. Briefly, at 7w, 2/3 of the mice were randomly transferred to a high-fat diet (HFD, 60% calories from fat, D12492, Research Diets, NJ), and weighed weekly. Eight weeks after HFF initiation, half of the HFD fed mice were switched back to NC diet for an additional 2w of obesity reversal (‘Reverse’ group), totaling 10 weeks of dietary intervention (**Fig. 1A**). Mid-aged (‘retired breeder’) mice were purchased at the age of 8 months from ENVIGO RMS Ltd. and acclimatized one mouse/cage until the age of 12m. Then, mice were allocated to the same dietary protocol as described above for 10 weeks. At the end of each experiment, mice were euthanized with isoflurane. The brain, blood, and gonadal (visceral) fat depot were weighed and collected.

**Figure 1.**
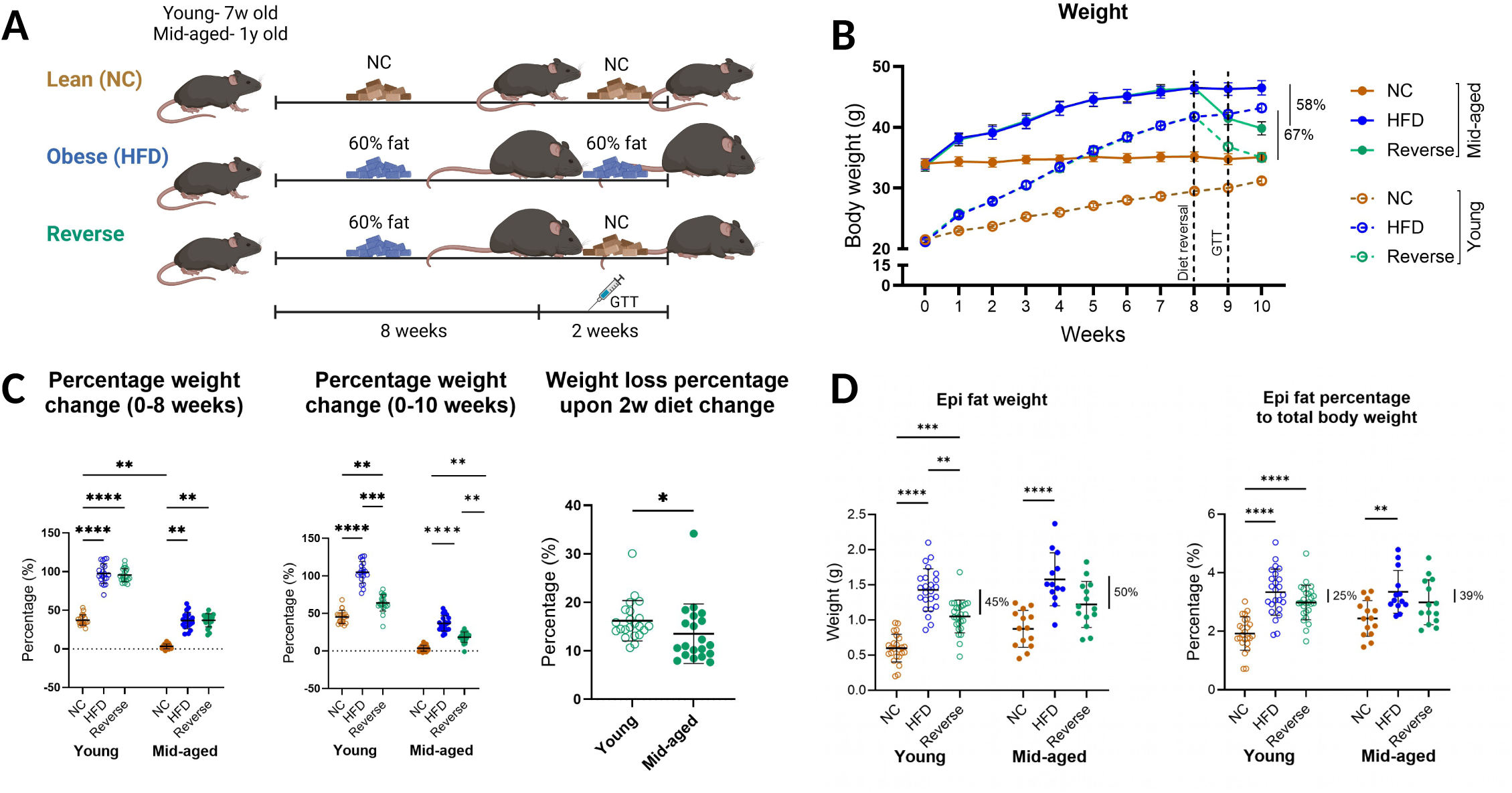
Weight dynamics reveal maintained ability for weight-loss in mid-aged mice. **A.** Mice were fed NC diet (20 and 18 mice per group of young and mid-aged, respectively) or HFD (42 and 33 mice per group of young and mid-aged, respectively) for 8 weeks, after which half of the HFD-fed mice were switched back to NC diet for 2 more weeks. GTT was performed at week 9. **B.** Weekly weight measurements in grams (mean ± SD). **C.** Percentage of weight change (mean ± SD) at the end of 8w of dietary intervention (left) and 10w of dietary intervention (middle). Percentage of weight-loss from maximal weight following 2w of dietary change in the reverse group (right). **D.** Epididymal fat pad weight (left) and percentage from total body weight (right) (mean). *-p<0.05, ** - p<0.01, *** - p<0.001, ****-p<0.0001.

### Glucose tolerance tests (GTT)

For GTT, at week 9 of the dietary intervention, mice were fasted for 6h and injected i.p. with

1.5 g/kg body weight glucose in sterile PBS solution. We used this based on considerations detailed in^17,18^. Blood samples were obtained from the tail vein immediately before injection and 15, 30, 45, 60, 90, and 120 minutes after glucose admission. Glucose levels were measured via a glucometer (FreeStyle freedom lite, Abbott).

### Brain collection and immunostaining

Animals were anesthetized with Isoflurane and perfused with 20mL of sterile PBS. Brains were excised and incubated overnight in a 4% paraformaldehyde (PFA) and then transferred to 30% sucrose solution for an additional 48h. The brains were sectioned into 45μm sections using a Leica cryostat. The sections were washed with PSB containing 0.05% Triton X-100 and subsequently incubated overnight at 4°C with anti-Iba1 and anti-p-NFκB antibodies, as detailed in **Table S1**. Next, sections were washed 3 times with 0.01% Tween-20 in PBS and then incubated with the appropriate secondary antibodies as detailed in **Table S1**. Finally, the sections were stained with DAPI for nuclear staining.

### Confocal microscopy and image analysis

Images were obtained using the Olympus FluoView 1000 confocal microscope (Olympus, Hamburg, Germany) and Zeiss LSM 880 confocal microscope at 800 × 800 and 1024 × 1024-pixel resolutions, each image contained 20-22 steps z-stacks, with 0.75µm step size. Whole-brain imaging was performed using ZEISS Cell Discoverer 7. Representative images of the whole hypothalamic area were created with FIJI software. Reconstitution of microglial cells was done using z-stack images. Imaris software (Oxford instruments, UK) was used to assess the volume of surfaces, as well as the colocalization of microglial pNFkB-positive cells, and to detect nuclei. Each analysis is based on all ARC microglia (as described in **Fig. 4A**), with a minimum of 21 analyzed microglia per mouse.

### Adipose tissue histology and PCR

Epididymal adipose tissue was subjected to several analyses upon its collection: 1. For histological studies–tissue was preserved in PFA 4%, processed and embedded in paraffin. Then, 5μm sections were stained with hematoxylin and eosin (H&E). Adipocyte size measurements were performed using QuPath software (v 0.3.2), with average measurement of (n=10 fields, including an average of 563 adipocytes/mouse). Crown-like structures were manually identified and counted per 100 adipocytes. 2. For gene expression studies-100 mg of tissue were snap frozen in liquid nitrogen. Then RNA was extracted using RNeasy Lipid tissue mini kit (Qiagen, Germany), and 2µg were reversed transcribed with High-Capacity cDNA kit (Applied Biosystems, Thermo-Fischer Scientific group, CA, USA). Taq-man probes were used to amplify specific genes (Il1rn-Mm00446186_m1, Ccl2-Mm00441242_m1), gene expression results were normalized to 2 house-keeping genes (Rplpo-Mm00725448_s1, Hprt1-Mm03024075_m1) and calculated using -ΔΔCt equation.

### RNA extraction and RNA sequencing data analysis

Total RNA was extracted from young and mid-aged mice’ hypothalamic, and epididymal adipose tissue from mid-aged mice with RNeasy Lipid Tissue Mini-Kit (QIAGEN, Germantown, MD) and quantified using Tape-Station. RNA from 45 mice (4 or more from each group) was sequenced using NextSeq 500 Illumina. Bioinformatic analysis was carried out as detailed in **Supplementary Methods**. Briefly, sequence reads were quality trimmed and filtered, aligned to the mouse genome (GRCm38) and quantified using RSEM. Statistical testing for differential expression was done using DESeq2^19^. In each tissue/age group, DEGs were identified as genes with an FDR-adjusted p-value<0.05 in at least one of the three possible between-group contrasts (HFD/NC, Reverse/NC, Reverse/HFD). DEGs were then divided into 8 predefined clusters, according to their fold change (FC) between diet groups, in an analysis we termed “patterns of change analysis” (**Supplemental Fig. 1**). FC was calculated from the z-score of normalized Log2 expression values and expressed on a linear scale. The following trends were defined: ‘up’ (FC≥1.2), ‘down’ (FC≤-1.2), and ‘unchanged’ (-1.2<FC<1.2). Clusters with equivalent but opposite patterns were grouped into broader categories (**Supplemental Fig. 1**): ‘persistent’ (up-unchanged and down-unchanged), ‘reversed’ (up-down and down-up), ‘aggravated’ (up-up and down-down), and ‘reverse only’ (unchanged-up and unchanged-down). Enrichment analysis was carried out on the original 8 clusters using Enrichr (https://maayanlab.cloud/Enrichr/).

### Statistical analysis

Statistical analysis was performed using GraphPad Prism 10.4.1. Statistical significance was assessed by a nonparametric unpaired t test or nonparametric ANOVA when two groups or more were compared.

### Data Resource Sharing and Availability

Bulk RNA-seq datasets and resources generated and analyzed during the current study are available from the corresponding author upon reasonable request.

## Results

### Reversal of dysglycemia and hypothalamus transcriptomic changes induced by obesity

Despite obesogenic memory, many metabolic and organ changes induced in mice by dietary obesity are fully reversible given sufficient time back on NC diet ^16,20^. However, the relative tissue-specific dynamics of reversal, and how mice’ age per-se, (i.e., in response to a similar duration of HFD), affects the reversal processes, have not been fully characterized. We therefore set up an ad-libitum HFD and dietary obesity reversal paradigm to assess the effects of aging on weight, glycemia, and hypothalamic changes early during obesity reversal, before excess weight-gain had been completely lost. For this purpose, young (7w) or mid-aged (1y) old mice were subjected to NC or HFD (60% fat) for 8 weeks. At this point, 50% of the HFD group was switched back to ad-libitum NC diet for up to 2 weeks (early obesity reversal group, ‘Rev’). Glucose tolerance test was performed 1 week after the dietary switch, and brains and gonadal adipose tissues were collected 2 weeks after the dietary switch (**Fig. 1A**).

Whereas young mice on NC gained weight during the entire 10-week protocol, mid-age mice weighed 60% more at baseline (21.36±1.45 vs. 34.19±3.10 g, respectively, p<0.0001), and were weight-stable (**Fig. 1B,C**). Ten weeks of HFD consumption enhanced body weight gain in the young mice, resulting in 38% weight gain compared with that of the NC control mice (p<0.0001). In mid-aged mice, HFD-induced weight gain stabilized by 7 weeks, resulting in 32% excess weight compared with that in age-matched controls (p<0.0001). Two weeks into the dietary switch back to NC resulted in a rapid, 67% and 58% loss of excess weight (defined as the difference between the mean weight of HFD-NC), in the young and mid-aged mice, respectively (p=NS). Yet, mid-aged mice lost less %total body weight when calculated relative to their individual weight at 8w of HFD (p=0.03, **Fig. 1C**). Adiposity, as assessed by epididymal fat weight, was similarly increased in young and mid-aged mice by HFD, and partially decreased during the 2 weeks dietary reversal (**Fig. 1D**). Yet, the percentage of epididymal fat pads to total body weight did not decrease significantly in Rev compared to HFD. Together, these data demonstrate similarities and differences in weight dynamics between the two ages: on NC, mid-aged mice are weight-stable while young mice exhibit continuous weight gain; in response to the dietary intervention, weight dynamics is similar, with mid-aged mice exhibiting significantly lower ability to rapidly lose previously-gained weight.

Given evidence for central sensing and regulation of peripheral glucose homeostasis, and obesity-induced hypothalamic changes, we next assessed glycemic parameters and their relation to hypothalamus transcriptomic changes. Ten weeks of HFD consumption induced fasting hyperglycemia and significant glucose intolerance in both mouse age-groups, the severity of which was greater in young than in the mid-aged mice (**Fig. 2A**). Nevertheless, 1w into the dietary switch, when only 44% and 42% of the excess body weight gained from 8w-HFD was lost in the reverse group (**Fig. 1B**), both fasting glycemia as well as glucose excursion after an intra-peritoneal glucose load were fully normalized (**Fig. 2A**).

**Figure 2.**
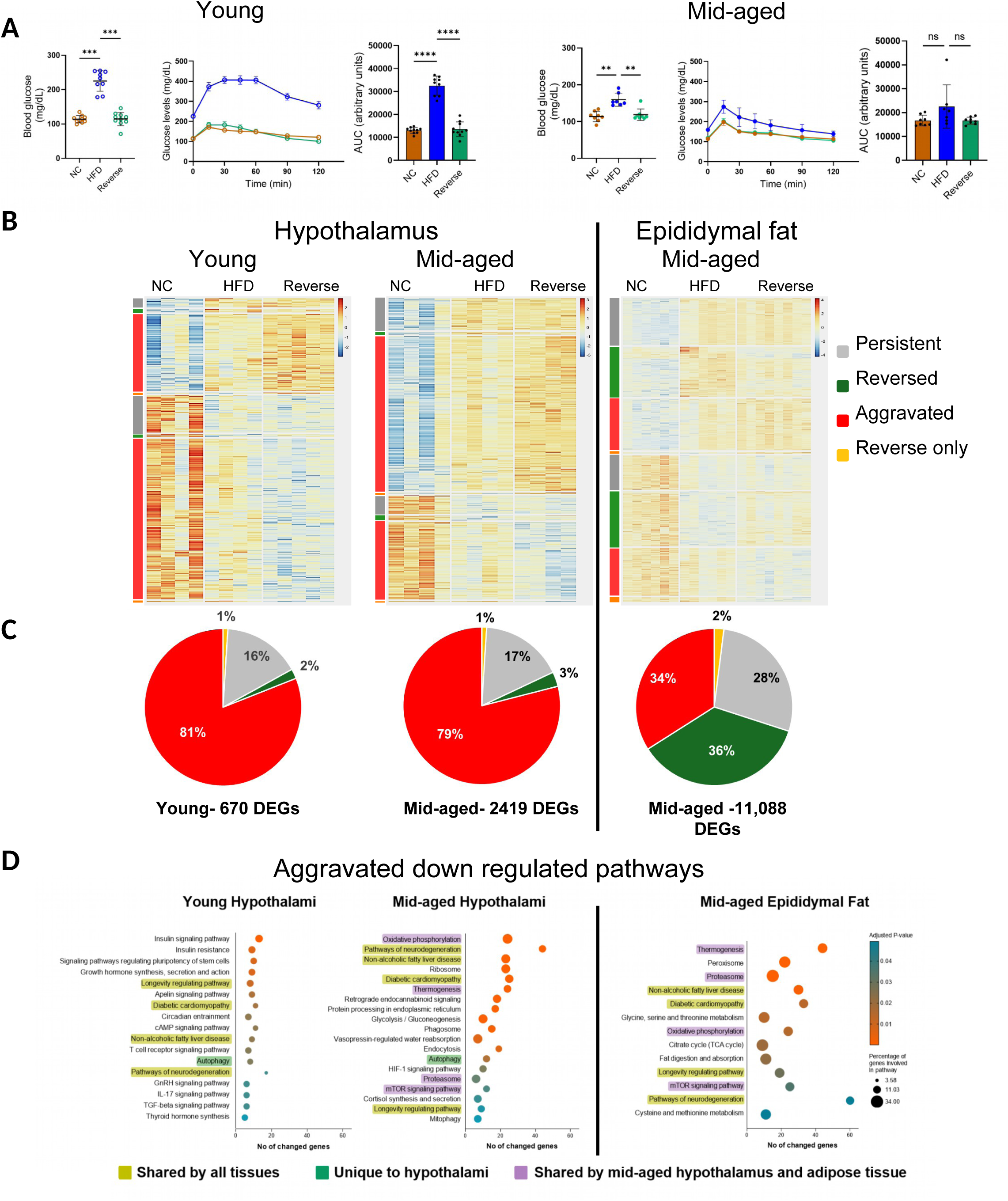
Early obesity reversal normalizes glucose tolerance but aggravates obesity-induced hypothalamic transcriptomic changes. **A.** From left to right: Fasting glucose levels, GTT curves and AUCs for young (left) and mid-aged (right) mice (mean± SD). *-p<0.05, ** - p<0.01, *** - p<0.001, ****-p<0.0001. **B.** Heatmap of DEGs (defined by FDR-adjusted p-value<0.05 in any contrast) from whole hypothalami bulk RNA-sequencing data from young (left) and mid-aged (middle) mice. Bulk RNA-sequencing from mid-aged mice epididymal fat (right). DEGs in each age group/tissue were divided into 8 predefined patterns of expression across the three diet groups (up-unchanged (uc); up-down; up-up; uc-up; down-uc; down-up; down-down; uc-down, see **Supplementary Methods** and **Supplemental Fig.1**), and genes within each pattern were hierarchically clustered. Patterns with equivalent but opposite trends (e.g. up-down and down-up) were depicted with the same color and further grouped for calculating their distribution within each tissue/age in Panel C **C.** Pie charts representing % of DEGs in 4 expression change patterns ( see **Supplemental Fig.1**), corresponding to the color code shown on the side of the heatmap, in B. **D.** KEGG pathway enrichment analysis of DEGs exhibiting downregulated aggravated pattern (down-down) as shown in **Supplemental Fig.1**, in hypothalami: young (n=367 genes) and mid-age (n=639 genes), and in epididymal adipose tissue (n=1813 genes), respectively. X-axis represents the number of DEGs involved in the pathway, circle diameter represents the percentage of DEGs out of the total genes in the pathway, and color corresponds to the adjusted p-value of the enrichment. KEGG pathways altered in young and mid-age hypothalami and in adipose tissue are highlighted in yellow, those common to hypothalami of young and mid-age mice are highlighted in green, and pathways common to hypothalami and adipose tissue only of mid-age mice in purple. Shown in the graph are the top enriched pathways sorted by FDR-adjustment p-value for each pathway. The full list of significantly enriched KEGG pathways (FDR-adjusted p-value<0.05( is provided in **Supplemental TableS4**.

To determine the effects of nutritional interventions on the hypothalamus in an unbiased manner, we used bulk RNA-sequencing to investigate the dynamic transcriptomic response of whole hypothalami to obesity and early obesity reversal in the two age groups. From mid-aged mice we also sequenced RNA from epididymal fat. In young mice’ hypothalami, 670 genes were differentially expressed in at least one of the pairwise comparisons between the groups. Remarkably, mid-aged mice hypothalami exhibited nearly 4-fold more genes (2,419 DEGs, **Fig. 2B,C**, **Table S2,S3**). By comparison, in mid-aged mice’ adipose tissue there were 11,088 DEGs. Notably, when DEGs were further subjected to patterns of change analysis (see ‘Research Design and Methods’), despite the difference in the number of DEGs, hypothalami of both age-groups exhibited ∼80% genes in the “aggravated” pattern, i.e., genes that were increased or decreased in the HFD vs. NC group, and further changed, in the same direction, in the Reversal group (**Supplementary Fig. 1A,B**). Interestingly, only a low percentage of genes exhibited the “reversed” gene expression pattern (2.2 and 2.6% of DEGs for young and mid-age mice, respectively) (**Fig.2B,C**, **Supplementary Fig. 1A,B**). This result may indicate a tissue-specific response pattern, as in adipose tissue only 34% of equally-defined DEGs exhibited the aggravated pattern, whereas 36% the reversed pattern (**Fig. 2B,C, Supplementary Fig. 1C**).

Since most hypothalamic DEGs exhibited the aggravated change pattern, pathway enrichment analysis was performed on this DEG group. Curiously, we did not detect enriched KEGG pathways among up-regulated-aggravated genes (i.e., increased in HFD vs. NC and further increased in the Reverse group) in the hypothalami of either age group, while in adipose tissue of mid-age mice up-regulated pathways expectedly signified obesity-related adipose tissue inflammation (**Supplementary Fig. 2A**). Yet, KEGG analysis of downregulated-aggravated genes (decrease in HFD vs. NC and further decreased in Reverse) revealed age-dependent differences in key pathways within the hypothalamus. In young mice, we observed a reduction in neuro-endocrine pathways related to insulin signaling, growth hormone synthesis, GnRH signaling, and circadian entrainment (and the related ‘circadian rhythm’ KEGG pathway was nearly significant (adjusted-p=0.06))(**Fig.2D, Supplementary TableS4**). In contrast, downregulated-aggravated genes in mid-aged mice were enriched in key metabolic pathways, including oxidative phosphorylation, glycolysis and gluconeogenesis, and mitophagy. Notably, oxidative phosphorylation, thermogenesis, mTOR signaling, and proteasome pathways were also enriched in epididymal adipose tissue of mid-aged mice, but not in the hypothalami of young mice (**Fig. 2D**). Interestingly, KEGG pathways of the aggravated down-regulated genes were more robustly altered (number and percentage of genes involved in the pathway, and the statistical significance) in the epididymal adipose tissue and hypothalami of mid-aged mice compared to KEGG pathway alterations in hypothalami of young mice (**Fig. 2D**).

Overall, pathway enrichment analysis indicated that early obesity reversal in both young and mid-aged mice further downregulated genes involved in key metabolic pathways (glycolysis, TCA cycle and oxidative phosphorylation), RNA surveillance, and mitophagy (**Table S2,S3**). Intriguingly, these genes were more robustly altered (in terms of the number of genes involved and statistical significance) in mid-aged mice than in young mice. Notably, while in hypothalami inflammatory pathways did not seem to be activated, such pathways were expectedly enriched among DEGs in the up-regulated adipose tissues of the same mid-age mice.

### Cytomorphometric analysis of hypothalamic microglia

Although we did not observe clear transcriptomic findings indicative of obesity-induced hypothalamic inflammation, we sought to determine whether the more pronounced transcriptomic changes in mid-age mice involve a more pronounced microglial reaction. Since the periventricular areas of the hypothalamus include nuclei involved in energy homeostasis such as the arcuate nucleus, we wished to characterize in detail microglial changes as the major immune cells in this region. Overall observation of the entire hypothalami revealed an elevation in Iba1 expression in obese mice, which was mostly evident in the ARC (**Fig. 3A,B**), suggesting spatially-distinct coordinates of hypothalamic inflammation. Upon acute obesity reversal, age seemed to affect the dynamics of the microglial response, with a minor reversal observed in younger mice versus an aggravated microglial pattern observed in mid-aged mice (**Fig. 3A,C,D**, versus **B,E,F,** respectively). Quantitative analysis revealed HFD-induced increases in both the number of microglia (percent of Iba1-positive cells), and the total volume occupied by Iba1-positive cells (**Fig. 3C-H**) consistent with previous reports in young mice ^1,3,5^. Two weeks of dietary switch back to NC in young mice were associated with relatively minor and statistically insignificant ∼24% reversal of hypothalamic microglial staining **(Fig. 3C,D),** while mid-aged mice exhibited reversal-induced further increase in percent microglia and their volume beyond the effect of HFD(**Fig. 3E,F**).

**Figure 3.**
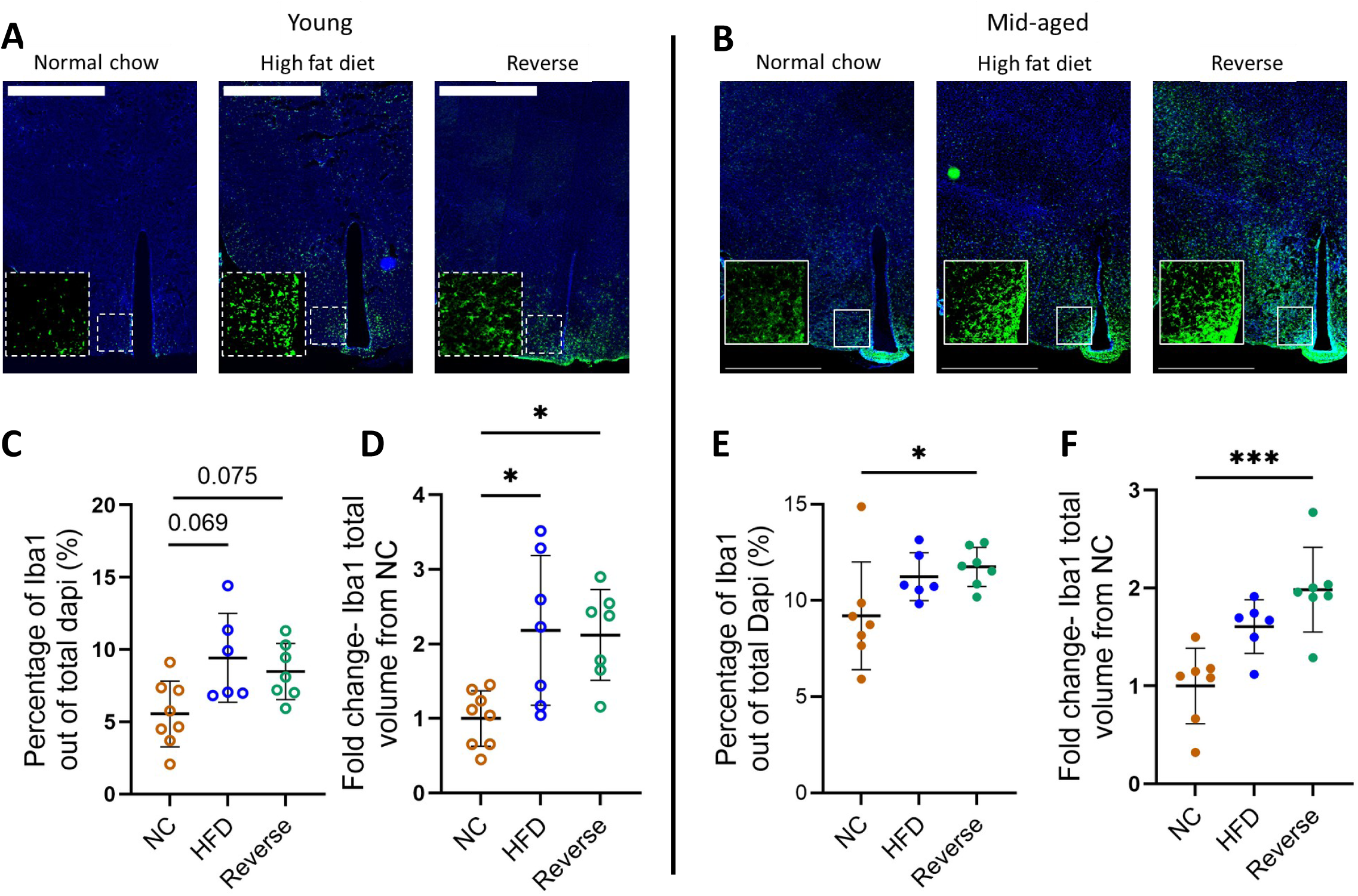
Hypothalamic microglia show age-dependent changes upon acute obesity reversal. A, **B.** Representative images of whole hypothalami scans from young (A) and mid-aged (B) mice (Dapi-blue, Iba1-green); scale bar, 1000 µm. **C, D.** Representative images of the ARC from young (left) and mid-aged (right) mice showing Iba1 expression (white) in response to the three diets. Scale bar, 100 µm. **E, G.** Young (E) and mid-aged (G) percentages of Iba1+ cells among the total identified nuclei in the ARC. (mean± SD). **F, H.** Total Iba1 volume (fold-change from NC) of young (F) and mid-aged (H) mice, compared with that in the NC group (mean ± SEM). *-p<0.05, ** - p<0.01, *** - p<0.001.

**Figure 4.**
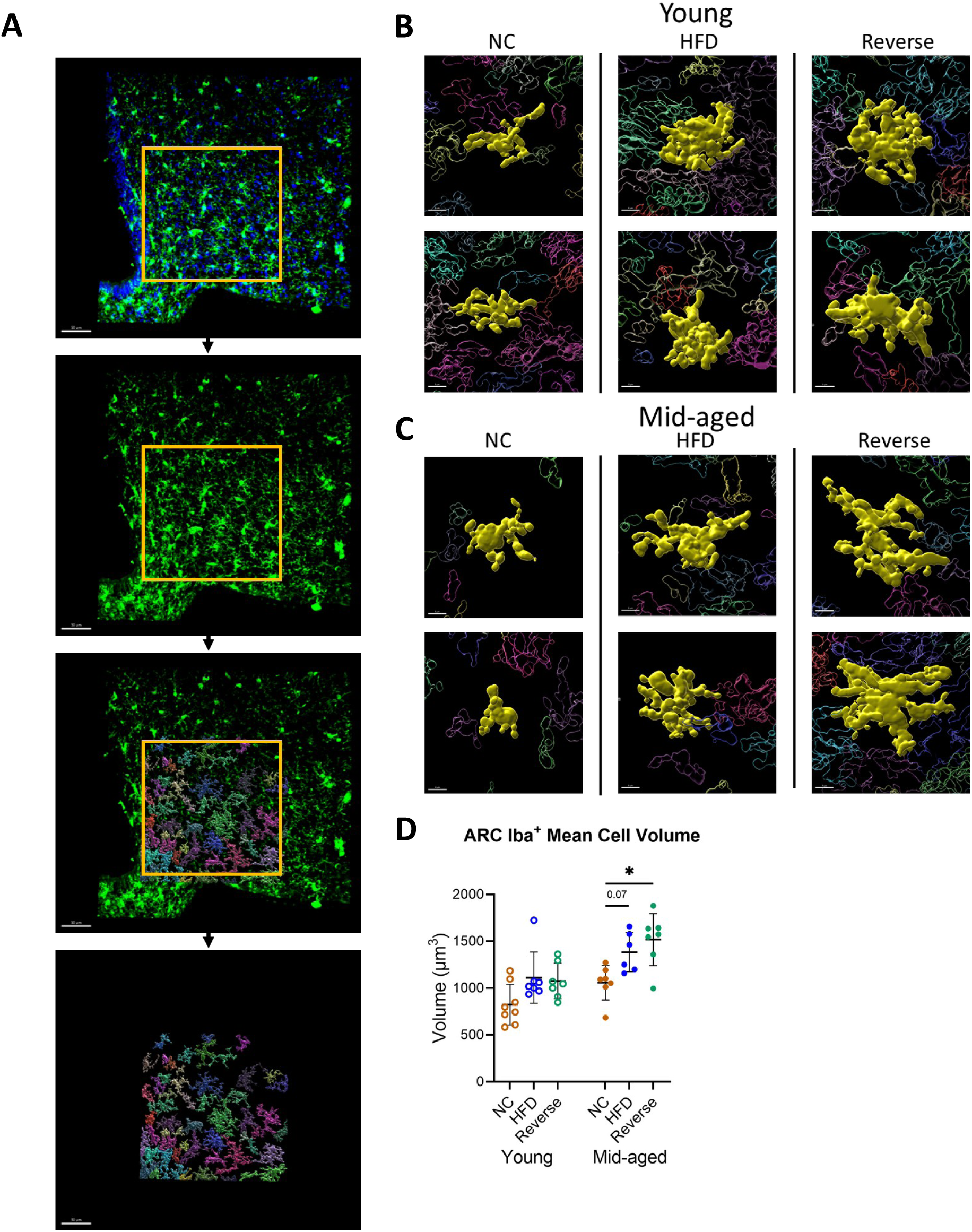
Single-cell microscopic volume analysis reveals age-dependent microglial aggravation upon acute obesity reversal. **A.** Scheme of Imaris confocal image-to-surface analysis transformation. **B, C.** Representative images of single microglia (yellow) in the ARC of young (B) and mid-aged (C) mice. Scale bar, 8 µm. **D.** Average volume of Iba1-positive cells in the ARC of young and mid-aged mice (mean ± SD). *-p<0.05.

To further increase the morphological resolution to the level of single microglia, we performedadvanced image analysis using Imaris (Methods). In equally selected areas of the arcuate nucleus, we assessed the 3-dimensional morphology of representative cells as exemplified in **Fig. 4A**. Compared with that in young NC mice, the mean volume of individual microglia seemed to increase and poorly reverse in the HFD and Reverse groups (**Fig. 4B,D**). In mid-aged mice, early obesity reversal-related augmentation of the cytomorphological changes were also readily apparent, with individual cells exhibiting larger somas and thicker branches than those of HFD-fed and NC hypothalamic microglia (**Fig. 4C,D**). Jointly, these cytomorphometric analyses are largely consistent with the notion that microglial changes do not resemble the dynamics of dysglycemia during weight-loss. Moreover, it signifies an age-dependent effect, wherein in mid-age mice further aggravation of the cytomorphological changes induced by obesity is induced by early obesity reversal.

### Obesity-induced NFκB activation in hypothalamic microglia in obesity reversal in mid-age

To molecularly characterize localized microglial changes in mid-aged mice, we performed immunofluorescence microscopy using anti-phosphorylated p65-NFκB antibodies. Such microscopic analysis could help determine if the cytomorphological changes represent an inflammation response, and can provide spatial coordinates for such changes within the hypothalamus. The total number of p-NFκB-positive nuclei, many of which were non-microglia cells, was increased in the ARC in HFD compared to NC, and was further higher in Rev (**Fig. 5A,B**). This trend was also observed among microglia (Iba1+/p-NFκB+ cells, **Fig. 5C**). Importantly, the volume of p-NFκB-positive microglia was markedly greater than that of p-NFκB-negative microglia under all three conditions (**Fig. 5D,E**). Jointly, these findings suggest that in mid-age mice, the increased total microglial volume in the ARC following obesity and its reversal is a combination of increased microglial number and larger volume of p-NFκB-positive microglial cells.

**Figure 5.**
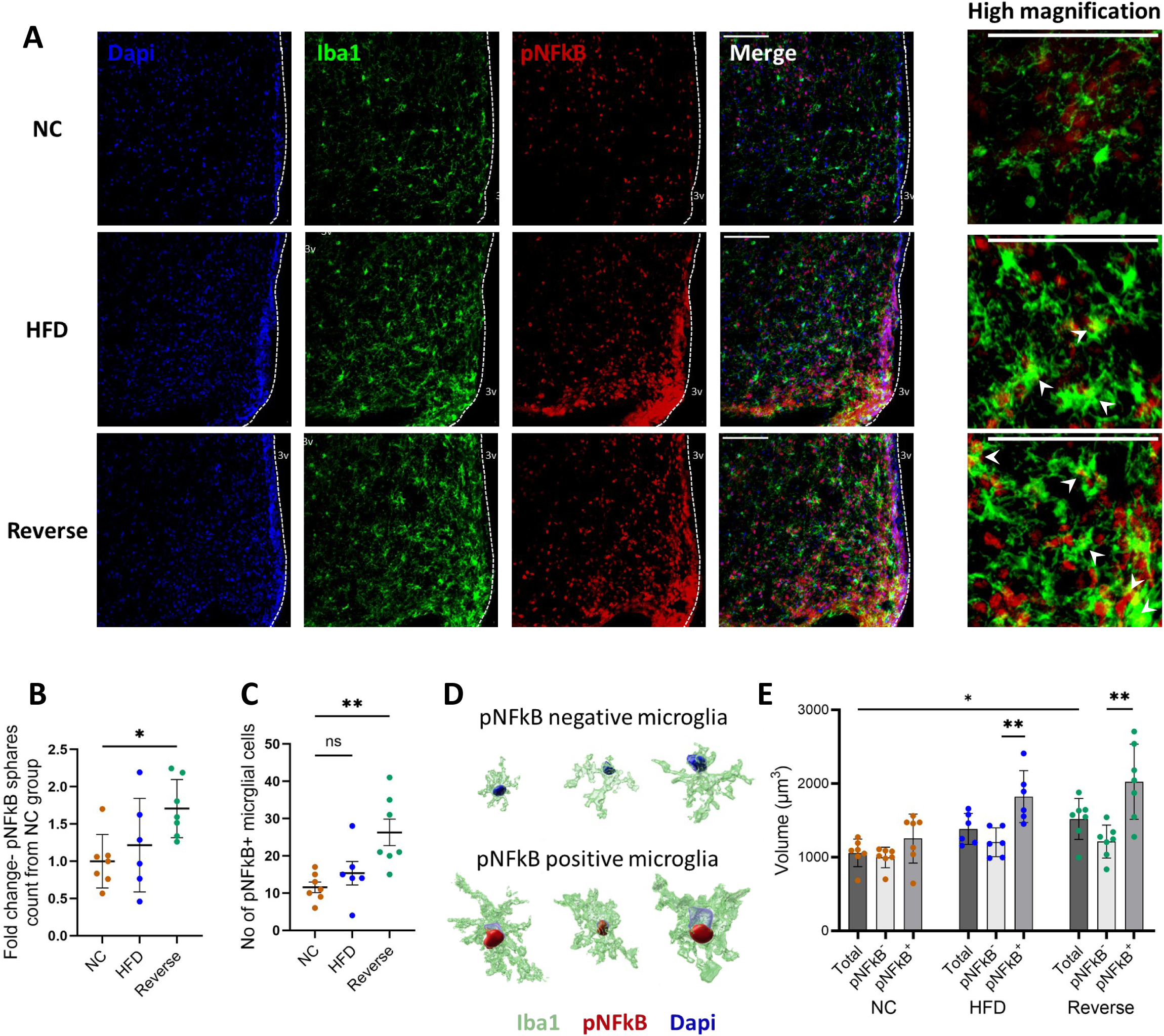
ARC microglia upregulate pNFkB expression upon acute obesity reversal in mid-aged mice. **A.** Representative images of mice ARC stained for Dapi (blue), Iba1 (green), and pNFkB (red). Scale bar, 100 µm. **B.** Fold change of pNFkB spheres number in the ARC of HFD and Reverse mid-aged mice compared with those in the NC group. **C.** Number of pNFkB+ microglia (mean ± SD). **D.** Representative images of single pNFkB- and pNFkB+ microglia from the three diet groups. **E.** Averaged volume of pNFkB+ or pNFkB-microglia in the ARC of mid-aged mice (mean ± SD). *-p<0.05, ** - p<0.01.

Finally, we wished to determine if hypothalamic changes in mid-aged mice could propose an adipose tissue-hypothalamic communication axis related. Previous obesity-reversal studies demonstrated in young mice the persistence of adipose tissue inflammation^16^, but how obesity-reversal affects adipose tissue inflammation in mid-aged mice is unknown. Histological examination of peri-epidydimal fat of mid-aged mice expectedly demonstrated adipocyte hypertrophy and increased crown-like structures induced by HFD (**Fig. 6A,B**). Obesity reversal trended to reverse adipocyte hypertrophy, but to increase CLS percentage (per 100 adipocytes, **Fig. 6B**). Consistent with previous results in young mice^16^, adipose tissue expression of inflammatory genes showed non-significant, if any, reversal (**Fig. 6C**). Intercorrelation analysis revealed strong association between adipose tissue inflammatory parameters, and in the hypothalamus – particularly between microglial volume and percent p-NFκB microglia. Intriguingly, microglia cell volume, though not hypothalamic p-NFκB-positive microglia percentage, significantly correlated (p≤0.02) with adipose tissue inflammatory parameters (**Fig. 6D**).

**Figure 6.**
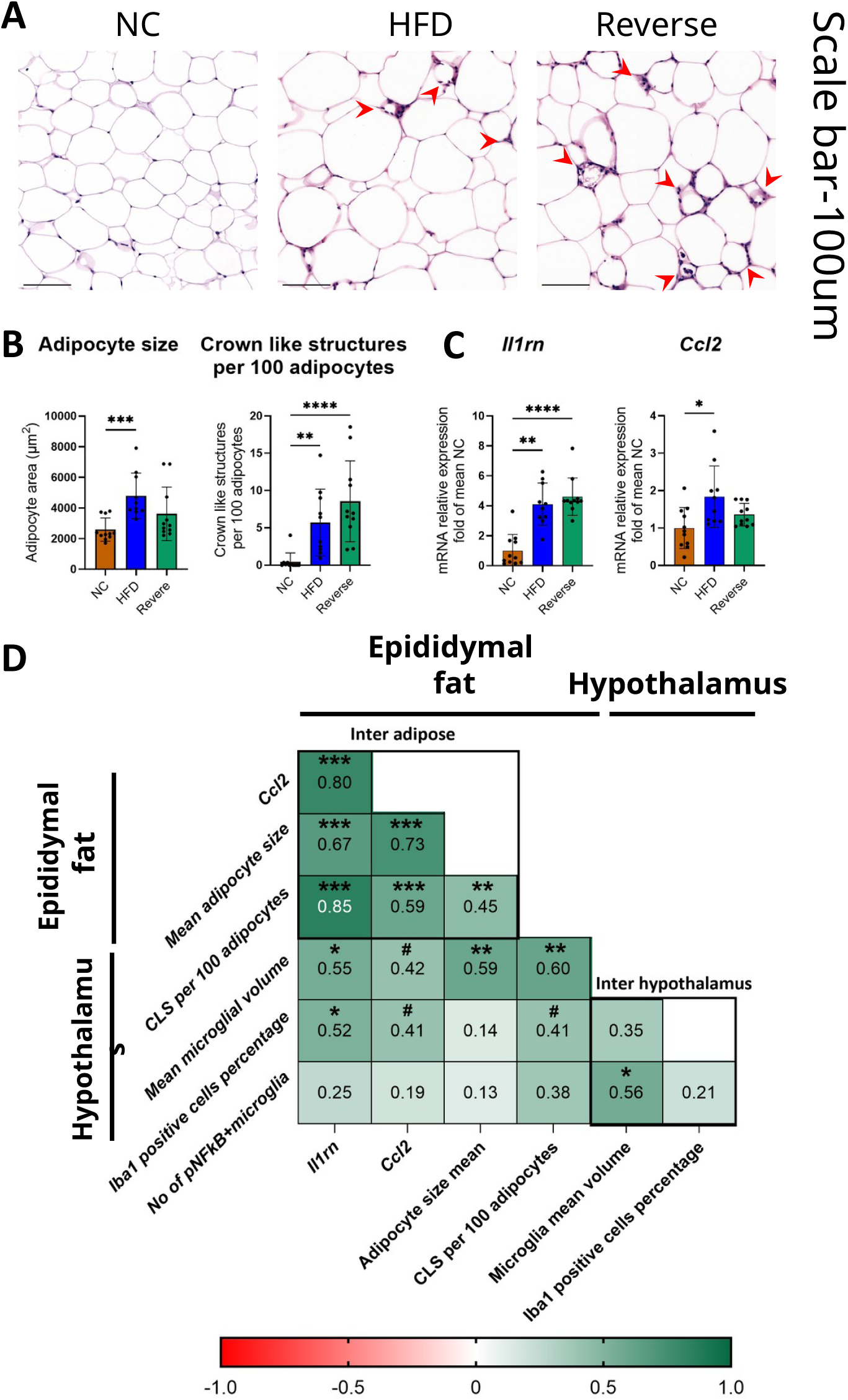
Acute weight loss induces an accumulation of CLS in the epididymal adipose tissue, positively correlating with hypothalamic microglial volume. A. Representative images of mice epididymal adipose tissue stained for H&E. Crown-like structures (CLS) marked with red arrows. Scale bar, 100 µm. B. Left: Adipocyte size (area) analysis (mean ± SD). Right: CLS count per 100 adipocytes (mean±SD). C. mRNA expression of *Il1rn* and *Ccl2* in mice epididymal adipose tissue (mean±SD). D. Correlation matrix analysis comparing epididymal adipose tissue and microscopic hypothalami analysis parameters of the same mice. *-p<0.05, ** - p<0.01, *** - p<0.001, ****-p<0.0001.

## Discussion

Hypothalamic inflammation, specifically involving microglia, has been proposed to link obesity with dysmetabolism^3,21^. In the present study, we hypothesized that metabolic improvement upon early weight-loss requires the reversibility of obesity-induced microglial changes. We also hypothesized that aging negatively affects the reversibility of changes induced by same-duration obesity, early in the course of its reversal. Our main findings show that compared to young mice, mid-aged mice exhibit somewhat blunted weight dynamics in response to HFD and switch back to NC, and less severe glucose intolerance in response to HFD-induced obesity. Nevertheless, as in young mice, dysglycemia is fully restored upon dietary switch. This remarkable “metabolic plasticity” during the early phase of obesity reversal stands out in contrast to the molecular changes in whole hypothalami and to the cytomorphological changes of hypothalamic (ARC) microglia: While younger mice exhibit minimal reversal of obesity-induced microglial cytomorphological changes upon acute weight-loss, mid-aged mice exhibit an aggravation of such changes. These changes correspond to weight-loss -associated aggravation of HFD-induced changes in gene expression during dietary switch back to NC, and to hypothalamic activation of NFκB. Our results provide several novel insights into the complex associations between weight dynamics, dysglycemia and hypothalamic microglial changes in obesity and its reversal, and into how age affects microglial adaptation to obesity and its reversal.

Central regulation of glucose homeostasis has been convincingly shown in rodent models since the 1990s^22^. Hypothalamic inflammation has been proposed to mediate metabolic dysregulation in obesity, but high glucose^23^ and/or FFA^24^ levels have also been proposed to induce hypothalamic microglial changes in obesity, potentially generating a “feed-forward” pathogenic feedback loop. Here, we investigated the associations between hypothalamic and microglial changes and dysglycemia during obesity reversal. Studies in both human and mouse have demonstrated that dysglycemia is rapidly corrected early during obesity reversal/weight-loss, clearly preceding the restoration of normal weight^16^. Microglial changes in response to obesity are also reversible, as shown in previous reports^20^, and as we also observed in young mice, but such reversal required 5 weeks of dietary switch from HFD to NC (data not shown). Since glycemia was fully-normalized as early as 1 week following the dietary switch, it occurred independently of significant reversal of whole hypothalamus obesity-induced transcriptomic changes or cytomorphological changes of hypothalamic microglia, contrasting our initial hypothesis. Moreover, while minor reversal of cytomorphometric microglial changes was observed in young mice, mid-age mice exhibited further aggravation of the obesity-induced microglial changes 2w into the dietary switch to NC. This corresponded to the transcriptomic changes in the hypothalamus, with most of the genes altered by obesity (up- or downregulated) exhibiting further aggravation of this effect early into the dietary switch. Interestingly, a recent study suggested that hypothalamic inflammation may be required to restore normoglycemia ^25^, and our results would suggest that this link may be more apparent in mid-age. It would be interesting to assess whether this represents an age-dependent difference in the function of microglia, as previously suggested^26^, and whether this augmented response to early weight-loss/dietary switch could explain the counter-intuitive, more severe impact of obesity and dysmetabolism in childhood and adolescence than in mid-age.

Microgliosis – a general term referring to the increased numbers (density) and morphological changes in microglia, has been documented in both humans and murine models of obesity^1–3^ and is generally considered a manifestation and/or a source/cause of hypothalamic inflammation during obesity. In early obesity reversal, our results demonstrate the complexity of deciphering hypothalamic inflammation: whole hypothalamus (bulk) RNA-seq demonstrated that this tissue exhibits much less pronounced transcriptomic alterations than adipose tissue does, and that among hypothalamic DEGs, inflammatory processes are not evidently enriched. This may indicate that the inflammatory response is more subtle, regulated mainly post-transcriptionally, and/or involves specific locations (hypothalamic nuclei) and/or specific cell types comprising the hypothalamus (microglia, astrocytes, neurons, etc.) – effects that are diluted when analyzing the whole hypothalamus. As for spatial considerations, indeed, MBH that include specific areas of the hypothalamus, such as the median eminence, ARC and PVN, seem to be more responsive to obesity and its reversal compared to more lateral regions (and nuclei) within the hypothalamus. These regions exhibit a greater number of microglial cells, a greater volume which they occupy, and a greater percentage of pNFκB-positive nuclei and total pNFκB signal intensity. Remarkably, under all conditions, cell volume of pNFκB-positive microglia was larger than that of pNFκB-negative microglia, suggesting a putative link between microglial cell volume and NFκB activation. These findings support the notion that microglial changes represent an inflammatory response, which likely also involves other cell types, as previously shown in obesity. Consistently, this may extend beyond the hypothalamus, as adipose tissue CLS correlated with hypothalamic microglia cell volume and pNFκB-positive cell percentage. The downstream effect of ARC microglial activation remains elusive, but as our experiment revealed a rapid metabolic improvement with further microglial activation, we suggest a possible beneficial effect of this acute, further microglial activation, in this context.

In conclusion, in mid-age mice, early during obesity reversal (weight-loss), rapid glycemic normalization occurs independently of reversal of hypothalamic microglial changes. Moreover, obesity-induced microglial changes are further aggravated early during weight-loss. This aggravation of acute obesity reversal response mirrors the whole hypothalamus transcriptome, which nevertheless does not strongly imply the activation of inflammatory signals. Identifying both the upstream signals for hypothalamic microgliosis, and its downstream consequences during weight-loss, could uncover possible ways to modulate the metabolic benefits, and possibly the ability to sustain, weight-loss in individuals living with obesity.

## Supporting information

all Supp data

## Acknowledgments

The authors wish to thank Prof. Rinke Stienstra (U. of Wageningen, The Netherlands), Prof. Martin Gericke (U of Leipzig, Germany), and Prof. Debbie Toiber (Ben-Gurion University, Israel), for helpful discussions. The work is part of A.Z and H.M’s Ph.D. Thesis (M.D.-Ph.D. track, Goldman school of Medical Sciences and The Kreitman school of graduate studies, BGU).

Conceptualization, A.Z., Y.H., U.Y., A.R., and A.M.; methodology, A.Z., Y.H., A.T., V.C-C., L.L.; investigation, A.Z., Y.H., A.T., V.C-C., Y.P., H.M., N.T.; writing – original draft, A.Z. and A.R.; writing – review and editing, A.Z., Y.H., V.C.-C., A.R., and A.M.; funding acquisition, A.R. and A.M.

