## Supplementary material for "Mid-aged mice rapidly normalize dysglycemia but aggravate obesity-induced hypothalamic and microglial changes upon dietary obesity reversal": all Supp data

#### Supplementary Extended Methods

***Bioinformatics Analysis of the RNA sequencing data***

Initial analysis of the raw sequence reads was carried out using the NeatSeq-Flow platform[1]. The sequences were quality trimmed and filtered using Trim Galore (v0.4.4_dev) and cutadapt (v1.15). Alignment of the reads to the mouse genome (GRCm38) was done using Bowtie2 (v2.3.4.3). The number of reads per gene per sample was counted using RSEM (v1.2.28). Quality assessment (QA) of the process was carried out using FASTQC (v0.11.8) and MultiQC (1.0.dev0). After trimming, each sample had an average of 34.4M reads with an average sequence length of 113bp. Subsequent statistical analyses were done using Neat_RNA-Seq (https://github.com/veredcc/Neat_RNA-Seq). Gene annotation was retrieved from Ensembl/BioMart web site. PCA of samples from both age groups revealed that the age effect was not very robust at the global transcriptome level. Therefore, we normalized and performed the statistical analysis for differential expression using all the samples together. In addition, the number of differentially expressed genes (DEGs) per comparison in the joint analysis was similar to that achieved when each age group was analyzed separately. Statistical testing for differential expression was done using the DESeq2 R package[2], considering 6 biological groups (design: "~ Diet_per_age") and including 6 contrasts (comparing groups within each age). In each age group, a list of DEGs was defined by unifying the genes having FDR adjusted p values<0.05 in any of the 3 contrasts in that age. Fold change (FC) is presented on a linear scale, where a minus sign denotes down-regulation. For visualization and clustering, expression values underwent variance stabilizing transformations in DESeq2.

*Clustering analyses* - Clustering analysis was done on DEGs from young and mid-age samples separately. Initially, we performed hierarchical clustering at the sample level with pheatmap. To help identify gene groups with expression patterns of interest, such as “obesity reversal" (i.e., when a HFD vs. NC difference – up- or down-regulation - was reversed in the Reverse vs. HFD comparison), we carried out partition clustering with two different methods: (i) Click/Expander using default and various manually-set "expected mean homogeneity" values (ii) eclust using group averages, where the number of clusters was set manually or estimated using the "gap" statistic with various k.max values. In both methods, with higher number of clusters we indeed detected the small clusters of interest, however this caused other (large) clusters to split to ostensibly meaningless subclusters. Therefore, we decided to divide the DEGs in each age to 8 predefined clusters, having the following trends between HFD vs. NC and Reverse vs. HFD: (i) up-unchanged (uc); (ii) up-down; (iii) up-up; (iv) uc-up; (v) down-uc; (vi) down-up; (vii) down-down; (viii) uc-down, where "up" means >= to FC_cutoff, "down" means <= -FC_cutoff, "uc" (unchanged) means between -FC_cutoff and FC_cutoff. In accordance with the values used in the unbiased clustering methods, the FC was calculated based on the z-score of the normalized Log2 expression values. To allow detection of mild changes in expression, as expected in hypothalamic regulatory processes, and considering that all DEGs have already passed a stringent cutoff of differential expression between at least one of the treatment groups, the FC cutoff was set to 1.2 (in linear scale). Clusters with equivalent but opposite patterns were unified as follows: i, v – "persistent"; ii, vi – "reversal"; iii, vii – "aggravated"; and iv, viii – "reverse only".

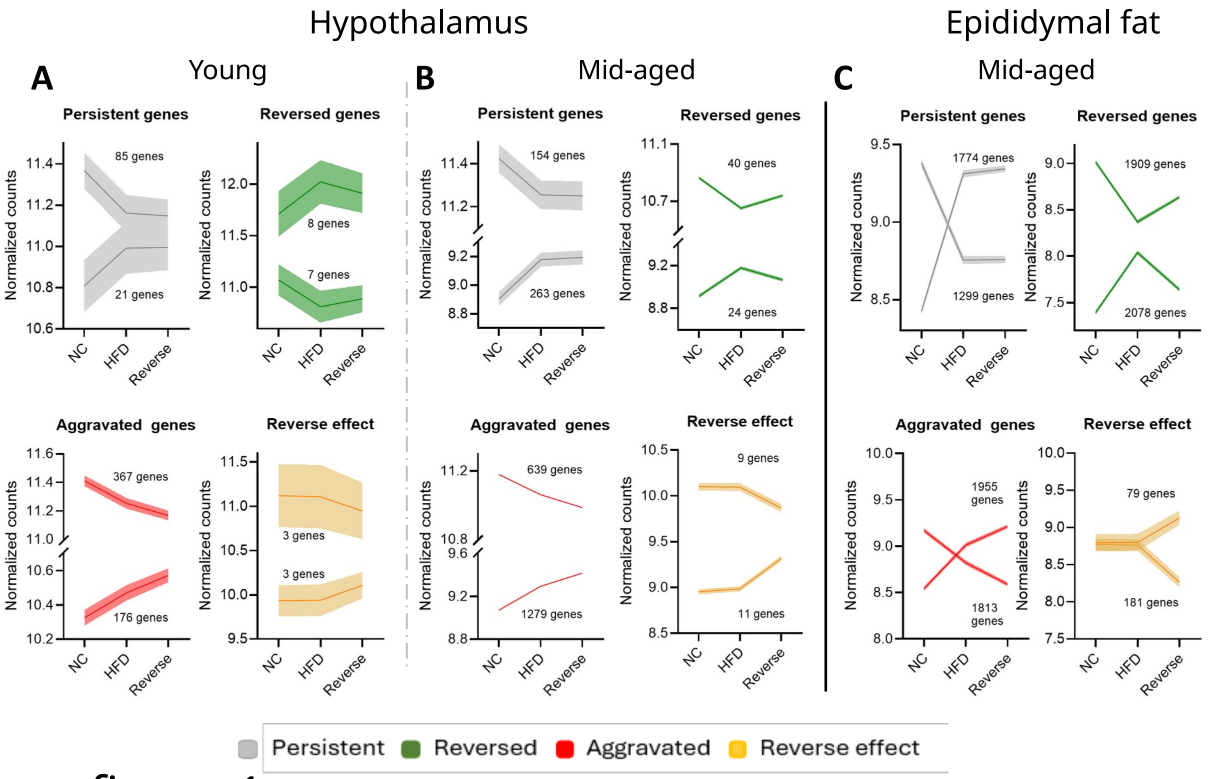

Supp. figure 1

### Adipose tissue KEGG pathway enrichment analysis: Aggravated-up genes (n=1955 genes)

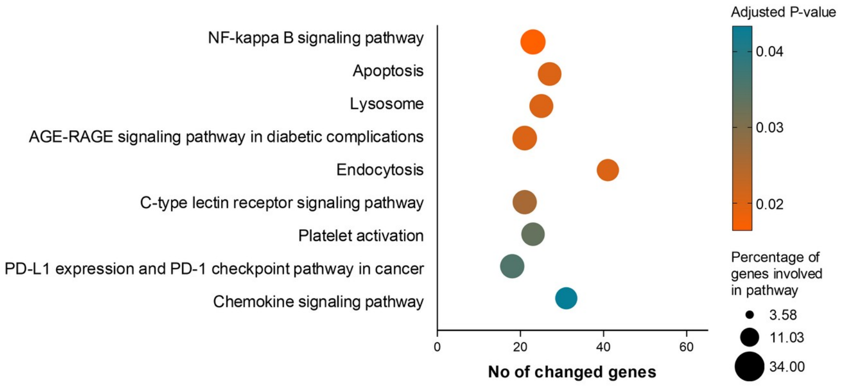

Supp. figure 2

| Antibody target/ Fluorophore | Dilution | Supplier | Catalog no. | Species | Application |
|---|---|---|---|---|---|
| Iba-1 | 1:500 | Abcam, United Kingdom | ab5076 | Goat | Primary antibody |
| pNFkB | 1:500 | Abcam, United Kingdom | ab86299 | Rabbit | Primary antibody |
| Alexa Fluor 488 | 1:500-250 | Abcam, United Kingdom | ab150073 | Donkey Anti Rabbit | Secondary antibody |
| Alexa Fluor 647 | 1:250 | Jackson ImmunoResearch Laboratories | 705-605-147 | Donkey Anti Goat | Secondary antibody |
| DAPI | 1:2500 | Biolegend, CA, USA | 422801 | - | Nuclear staining |

**Table S1: Used Antibodies**

**Supplemental table S2**

Gene information / Normalized counts per sample

| Ensembl_Gene_ID | Gene_Symbol | Gene_Title | Gene_Type | Young_01 (Y_NC) | Young_02 (Y_NC) | Young_03 (Y_NC) | Young_05 (Y_NC) | Young_06 (Y_HFD) | Young_07 (Y_HFD) | Young_08 (Y_HFD) | Young_09 (Y_HFD) | Young_11 (Y_Reverse) | Young_12 (Y_Reverse) | Young_13 (Y_Reverse) | Young_14 (Y_Reverse) | Young_15 (Y_Reverse) |
|---|---|---|---|---|---|---|---|---|---|---|---|---|---|---|---|---|
| ENSMUSG00000020889 | Nr1d1 | nuclear receptor subfan | protein_coding | 1,801 | 1,605 | 1,856 | 1,902 | 2,232 | 2,419 | 2,040 | 1,904 | 2,052 | 2,164 | 2,154 | 1,986 | 2,227 |
| ENSMUSG00000019027 | Dnah1 | dynein, axonemal, heav | protein_coding | 1,395 | 1,404 | 1,354 | 1,433 | 1,952 | 1,538 | 1,501 | 1,641 | 1,561 | 1,634 | 1,570 | 1,735 | 1,642 |
| ENSMUSG00000028137 | Celf3 | CUGBP, Elav-like family | protein_coding | 3,144 | 3,166 | 2,838 | 2,832 | 3,850 | 3,542 | 3,369 | 2,965 | 3,273 | 3,512 | 3,364 | 3,562 | 3,319 |
| ENSMUSG00000021432 | Slc35b3 | solute carrier family 35, | protein_coding | 207 | 265 | 266 | 212 | 327 | 304 | 311 | 323 | 278 | 330 | 333 | 298 | 382 |
| ENSMUSG00000029538 | Srsf9 | serine and arginine-rich | protein_coding | 612 | 623 | 666 | 616 | 744 | 687 | 656 | 731 | 685 | 736 | 677 | 715 | 750 |
| ENSMUSG00000056972 | Magel2 | MAGE family member L | protein_coding | 1,732 | 2,142 | 1,921 | 1,736 | 2,543 | 2,146 | 2,001 | 2,329 | 2,047 | 2,376 | 2,099 | 2,343 | 2,320 |
| ENSMUSG00000038141 | Tmem181a | transmembrane proteir | protein_coding | 1,584 | 1,704 | 1,776 | 1,781 | 1,836 | 1,881 | 1,885 | 1,890 | 1,828 | 1,838 | 1,844 | 1,966 | 1,884 |
| ENSMUSG00000039199 | Zdhhc1 | zinc finger, DHHC doma | protein_coding | 678 | 844 | 833 | 809 | 962 | 880 | 920 | 916 | 893 | 935 | 935 | 973 | 953 |
| ENSMUSG00000044308 | Ubr3 | ubiquitin protein ligase | protein_coding | 5,635 | 6,433 | 6,135 | 5,969 | 7,261 | 7,107 | 6,758 | 7,023 | 6,858 | 6,615 | 7,055 | 7,501 | 6,829 |
| ENSMUSG00000048310 | Pskh1 | protein serine kinase H: | protein_coding | 240 | 253 | 259 | 251 | 291 | 323 | 303 | 302 | 304 | 290 | 322 | 310 | 314 |
| ENSMUSG00000053754 | Chd8 | chromodomain helicase | protein_coding | 3,666 | 4,191 | 3,746 | 3,957 | 4,569 | 4,617 | 4,908 | 3,958 | 5,077 | 4,585 | 4,525 | 4,069 | 4,677 |
| ENSMUSG00000056342 | Usp34 | ubiquitin specific peptic | protein_coding | 7,870 | 8,195 | 8,190 | 8,479 | 8,801 | 9,100 | 9,289 | 9,080 | 9,822 | 9,171 | 8,689 | 8,860 | 8,767 |
| ENSMUSG00000029419 | Ajm1 | apical junction compon | protein_coding | 1,727 | 1,826 | 1,884 | 1,657 | 2,029 | 1,867 | 1,986 | 2,114 | 1,955 | 2,130 | 2,025 | 2,050 | 1,942 |
| ENSMUSG00000044700 | Tmem201 | transmembrane proteir | protein_coding | 1,836 | 1,811 | 2,055 | 1,773 | 2,083 | 2,004 | 2,070 | 2,229 | 2,081 | 2,140 | 2,129 | 2,165 | 1,994 |
| ENSMUSG00000035713 | Usp35 | ubiquitin specific peptic | protein_coding | 499 | 592 | 699 | 552 | 820 | 684 | 770 | 757 | 795 | 823 | 800 | 761 | 702 |
| ENSMUSG00000046020 | Pofut1 | protein O-fucosyltransf | protein_coding | 775 | 903 | 1,036 | 798 | 1,065 | 1,059 | 1,051 | 1,027 | 1,078 | 1,013 | 1,126 | 1,082 | 1,011 |
| ENSMUSG00000040327 | Cul9 | cullin 9 | protein_coding | 1,745 | 1,783 | 1,930 | 1,889 | 2,192 | 2,004 | 2,143 | 2,236 | 2,256 | 2,114 | 2,096 | 2,198 | 1,891 |
| ENSMUSG00000020802 | Ube2o | ubiquitin-conjugating e | protein_coding | 4,096 | 4,128 | 4,382 | 4,309 | 4,545 | 4,321 | 4,465 | 4,756 | 4,798 | 4,462 | 4,560 | 4,526 | 4,418 |
| ENSMUSG00000037224 | Zfyve28 | zinc finger, FYVE domai | protein_coding | 406 | 417 | 435 | 414 | 473 | 436 | 487 | 534 | 538 | 485 | 513 | 465 | 459 |
| ENSMUSG00000018401 | Mtmr4 | myotubularin related pr | protein_coding | 2,401 | 2,459 | 2,481 | 2,531 | 2,783 | 2,628 | 2,695 | 2,863 | 2,726 | 2,939 | 2,667 | 2,681 | 2,654 |
| ENSMUSG00000027006 | Dnajc10 | DnaJ heat shock proteir | protein_coding | 2,549 | 2,514 | 2,615 | 2,582 | 2,798 | 2,796 | 2,838 | 3,192 | 2,878 | 2,940 | 2,984 | 2,878 | 2,784 |
| ENSMUSG00000006342 | Susd2 | sushi domain containin | protein_coding | 1,031 | 1,104 | 1,055 | 1,111 | 1,401 | 1,191 | 1,203 | 1,235 | 1,186 | 1,236 | 1,210 | 1,146 | 1,222 |
| ENSMUSG00000021091 | Serpina3n | serine (or cysteine) pep | protein_coding | 1,310 | 1,266 | 870 | 1,188 | 2,572 | 1,726 | 1,700 | 2,009 | 1,288 | 1,692 | 1,386 | 1,468 | 1,462 |
| ENSMUSG00000051790 | Nlgn2 | neuroligin 2 | protein_coding | 8,719 | 8,253 | 8,040 | 8,227 | 9,370 | 8,813 | 9,116 | 9,523 | 8,521 | 8,715 | 8,637 | 9,770 | 8,950 |
| ENSMUSG00000026204 | Ptprn | protein tyrosine phosph | protein_coding | 13,125 | 13,921 | 13,475 | 13,578 | 15,894 | 14,426 | 14,740 | 15,964 | 14,723 | 15,403 | 14,567 | 15,643 | 14,240 |
| ENSMUSG00000045039 | Megf8 | multiple EGF-like-doma | protein_coding | 6,429 | 6,905 | 7,481 | 6,766 | 7,817 | 7,275 | 7,833 | 8,177 | 7,776 | 7,493 | 7,643 | 7,915 | 7,319 |
| ENSMUSG00000022761 | Lztr1 | leucine-zipper-like trans | protein_coding | 2,201 | 1,980 | 2,238 | 2,012 | 2,505 | 2,443 | 2,171 | 2,511 | 2,313 | 2,391 | 2,364 | 2,298 | 2,325 |
| ENSMUSG00000037727 | Avp | arginine vasopressin | protein_coding | 3,032 | 2,205 | 2,717 | 2,642 | 5,172 | 3,527 | 4,149 | 6,136 | 3,164 | 4,209 | 3,784 | 4,295 | 4,205 |
| ENSMUSG00000040859 | Bsdc1 | BSD domain containing | protein_coding | 1,456 | 1,448 | 1,461 | 1,445 | 1,686 | 1,534 | 1,635 | 1,757 | 1,641 | 1,587 | 1,629 | 1,545 | 1,634 |
| ENSMUSG00000004996 | Mri1 | methylthioribose-1-pho | protein_coding | 237 | 277 | 282 | 249 | 283 | 306 | 320 | 312 | 317 | 303 | 300 | 353 | 336 |
| ENSMUSG00000051747 | Ttn | titin | protein_coding | 56 | 87 | 200 | 116 | 311 | 287 | 492 | 125 | 180 | 362 | 317 | 575 | 399 |
| ENSMUSG00000001870 | Ltbp1 | latent transforming gro | protein_coding | 187 | 265 | 311 | 184 | 330 | 377 | 282 | 227 | 396 | 348 | 351 | 310 | 315 |
| ENSMUSG00000023826 | Prkn | parkin RBR E3 ubiquitin | protein_coding | 189 | 280 | 330 | 161 | 359 | 344 | 319 | 267 | 376 | 429 | 412 | 370 | 357 |
| ENSMUSG00000029168 | Dpysl5 | dihydropyrimidinase-lik | protein_coding | 1,984 | 2,492 | 2,537 | 2,096 | 2,696 | 2,436 | 2,498 | 2,372 | 2,683 | 2,805 | 2,721 | 2,602 | 2,611 |
| ENSMUSG00000045482 | Trrap | transformation/transcri | protein_coding | 4,750 | 5,791 | 5,745 | 4,915 | 6,329 | 5,946 | 6,055 | 5,421 | 6,238 | 6,097 | 6,279 | 6,034 | 5,998 |
| ENSMUSG00000053693 | Mast1 | microtubule associated | protein_coding | 3,260 | 3,744 | 3,628 | 3,567 | 3,848 | 3,839 | 3,926 | 3,667 | 4,017 | 4,142 | 4,009 | 3,850 | 3,649 |
| ENSMUSG00000033767 | Tmem131l | transmembrane 131 lik | protein_coding | 255 | 442 | 384 | 337 | 455 | 512 | 501 | 343 | 501 | 486 | 539 | 453 | 450 |
| [illegible] | [illegible] | [illegible] | [illegible] | [illegible] | [illegible] | [illegible] | [illegible] | [illegible] | [illegible] | [illegible] | [illegible] | [illegible] | [illegible] | [illegible] | [illegible] | [illegible] |

DESeq2 results for differential expression / Expression pattern

| Gene_Symbol | linearFC.Y_HF (Y_HFD.vs.NC) | pvalue.Y_HFD | padj.Y_HFD.v | pass.Y_HFD.v | linearFC.Y_Re (Y_Reverse.vs.NC) | pvalue.Y_Rev | padj.Y_Revers | pass.Y_Revers | linearFC.Y_Re (Y_Reverse.vs.HFD) | pvalue.Y_Rev | padj.Y_Revers | pass.Y_Revers | Pattern_No. | Clustering_or |
|---|---|---|---|---|---|---|---|---|---|---|---|---|---|---|
| Nr1d1 | 1.2 | 0.00142 | 0.122 | | 1.18 | 0.0021 | 0.0431 | up | -1.02 | 0.779 | 1 | | 1 | 1 |
| Dnah1 | 1.19 | 0.00126 | 0.121 | | 1.17 | 0.0024 | 0.0461 | up | -1.02 | 0.721 | 1 | | 1 | 2 |
| Celf3 | 1.15 | 0.00226 | 0.141 | | 1.14 | 0.00235 | 0.0459 | up | -1.01 | 0.864 | 1 | | 1 | 3 |
| Slc35b3 | 1.32 | 0.00079 | | | 1.36 | 0.000131 | 0.0103 | up | 1.02 | 0.75 | 1 | | 1 | 4 |
| Srsf9 | 1.12 | 0.00579 | | | 1.13 | 0.00182 | 0.04 | up | 1.01 | 0.801 | 1 | | 1 | 5 |
| Magel2 | 1.2 | 0.00245 | 0.146 | | 1.19 | 0.00237 | 0.046 | up | -1.01 | 0.883 | 1 | | 1 | 6 |
| Tmem181a | 1.09 | 0.00121 | 0.121 | | 1.09 | 0.000783 | 0.0262 | up | -1 | 0.986 | 1 | | 1 | 7 |
| Zdhhc1 | 1.16 | 0.0026 | 0.151 | | 1.18 | 0.000337 | 0.0174 | up | 1.02 | 0.66 | 1 | | 1 | 8 |
| Ubr3 | 1.16 | 0.000153 | 0.0477 | up | 1.15 | 0.000184 | 0.0125 | up | -1.01 | 0.806 | 1 | | 1 | 9 |
| Pskh1 | 1.21 | 0.00081 | | | 1.23 | 0.00024 | 0.0139 | up | 1.01 | 0.837 | 1 | | 1 | 10 |
| Chd8 | 1.16 | 0.00121 | 0.121 | | 1.18 | 0.000155 | 0.011 | up | 1.02 | 0.705 | 1 | | 1 | 11 |
| Usp34 | 1.11 | 0.000129 | 0.0477 | up | 1.11 | 0.0000673 | 0.00711 | up | -1 | 0.969 | 1 | | 1 | 12 |
| Ajm1 | 1.13 | 0.00136 | 0.121 | | 1.14 | 0.000257 | 0.0145 | up | 1.01 | 0.761 | 1 | | 1 | 13 |
| Tmem201 | 1.12 | 0.000906 | 0.11 | | 1.12 | 0.000391 | 0.0191 | up | 1 | 0.944 | 1 | | 1 | 14 |
| Usp35 | 1.29 | 0.00011 | | | 1.32 | 0.00000879 | 0.00173 | up | 1.02 | 0.692 | 1 | | 1 | 15 |
| Pofut1 | 1.19 | 0.0027 | 0.151 | | 1.21 | 0.000791 | 0.0262 | up | 1.01 | 0.834 | 1 | | 1 | 16 |
| Cul9 | 1.17 | 0.00145 | 0.123 | | 1.15 | 0.00267 | 0.0486 | up | -1.02 | 0.728 | 1 | | 1 | 17 |
| Ube2o | 1.07 | 0.00464 | 0.182 | | 1.08 | 0.00112 | 0.0311 | up | 1.01 | 0.769 | 1 | | 1 | 18 |
| Zfyve28 | 1.15 | 0.00736 | | | 1.17 | 0.00154 | 0.0367 | up | 1.02 | 0.699 | 1 | | 1 | 19 |
| Mtmr4 | 1.11 | 0.000108 | 0.0477 | up | 1.11 | 0.0000944 | 0.00878 | up | -1 | 0.885 | 1 | | 1 | 20 |
| Dnajc10 | 1.13 | 0.00131 | 0.121 | | 1.13 | 0.00118 | 0.0318 | up | -1 | 0.895 | 1 | | 1 | 21 |
| Susd2 | 1.17 | 0.000166 | 0.0477 | up | 1.11 | 0.00591 | 0.0709 | | -1.05 | 0.228 | 1 | | 2 | 22 |
| Serpina3n | 1.73 | 0.00000876 | 0.0343 | up | 1.26 | 0.0481 | 0.21 | | -1.37 | 0.0067 | 1 | | 2 | 23 |
| Nlgn2 | 1.11 | 0.00012 | 0.0477 | up | 1.07 | 0.00508 | 0.0654 | | -1.03 | 0.212 | 1 | | 2 | 24 |
| Ptprn | 1.13 | 0.0000977 | 0.0477 | up | 1.1 | 0.00087 | 0.0273 | up | -1.02 | 0.439 | 1 | | 2 | 25 |
| Megf8 | 1.13 | 0.0000327 | 0.0343 | up | 1.11 | 0.000239 | 0.0139 | up | -1.02 | 0.485 | 1 | | 2 | 26 |
| Lztr1 | 1.14 | 0.000223 | 0.0519 | | 1.11 | 0.00259 | 0.0479 | up | -1.03 | 0.384 | 1 | | 2 | 27 |
| Avp | 1.79 | 0.0000416 | 0.0343 | up | 1.48 | 0.00346 | 0.0549 | | -1.21 | 0.163 | 1 | | 2 | 28 |
| Bsdc1 | 1.14 | 0.000147 | 0.0477 | up | 1.11 | 0.00193 | 0.0411 | up | -1.03 | 0.38 | 1 | | 2 | 29 |
| Mri1 | 1.16 | 0.0265 | | | 1.22 | 0.00176 | 0.0394 | up | 1.05 | 0.405 | 1 | | 3 | 30 |
| Ttn | 2.65 | 0.00626 | | | 3.2 | 0.000587 | 0.0229 | up | 1.21 | 0.576 | 1 | | 3 | 31 |
| Ltbp1 | 1.28 | 0.0508 | | | 1.45 | 0.00202 | 0.0419 | up | 1.13 | 0.296 | 1 | | 3 | 32 |
| Prkn | 1.34 | 0.00875 | | | 1.61 | 0.00000524 | 0.00128 | up | 1.21 | 0.0686 | 1 | | 3 | 33 |
| Dpysl5 | 1.1 | 0.0673 | 0.39 | | 1.18 | 0.000639 | 0.0241 | up | 1.07 | 0.134 | 1 | | 3 | 34 |
| Trrap | 1.12 | 0.024 | 0.292 | | 1.16 | 0.00228 | 0.0452 | up | 1.03 | 0.5 | 1 | | 3 | 35 |
| Mast1 | 1.07 | 0.0244 | 0.293 | | 1.11 | 0.000841 | 0.0273 | up | 1.03 | 0.326 | 1 | | 3 | 36 |
| Tmem131l | 1.27 | 0.0159 | | | 1.37 | 0.000953 | 0.0289 | up | 1.08 | 0.438 | 1 | | 3 | 37 |
| [illegible] | [illegible] | [illegible] | [illegible] | [illegible] | [illegible] | [illegible] | [illegible] | [illegible] | [illegible] | [illegible] | [illegible] | [illegible] | [illegible] | [illegible] |

z-score expression values for heatmap visualization

| Gene_Symbol | Young_01.zsc (Y_NC) | Young_02.zsc (Y_NC) | Young_03.zsc (Y_NC) | Young_05.zsc (Y_NC) | Young_06.zsc (Y_HFD) | Young_07.zsc (Y_HFD) | Young_08.zsc (Y_HFD) | Young_09.zsc (Y_HFD) | Young_11.zsc (Y_Reverse) | Young_12.zsc (Y_Reverse) | Young_13.zsc (Y_Reverse) | Young_14.zsc (Y_Reverse) | Young_15.zsc (Y_Reverse) |
|---|---|---|---|---|---|---|---|---|---|---|---|---|---|
| Nr1d1 | -1.0364631 | -2.08150336 | -0.7624696 | -0.53644662 | 0.93998596 | 1.69164682 | 0.10619869 | -0.52727323 | 0.16275774 | 0.65229223 | 0.60933483 | -0.13838658 | 0.92032621 |
| Dnah1 | -1.10138655 | -1.03399338 | -1.38803263 | -0.83297033 | 2.25102844 | -0.13961618 | -0.37852281 | 0.50656893 | 0.01084451 | 0.46274609 | 0.06785662 | 1.06241798 | 0.51305931 |
| Celf3 | -0.44946635 | -0.37068504 | -1.56246307 | -1.58485616 | 1.77676163 | 0.85725131 | 0.30895022 | -1.08828526 | -0.00747239 | 0.76443235 | 0.29244121 | 0.91938164 | 0.14400991 |
| Slc35b3 | -1.88019263 | -0.55341195 | -0.54082514 | -1.76865891 | 0.66529944 | 0.22892737 | 0.35526573 | 0.57824334 | -0.28702012 | 0.72212796 | 0.77330782 | 0.10390766 | 1.60302944 |
| Srsf9 | -1.51577863 | -1.27010654 | -0.35136892 | -1.43789857 | 1.19612791 | 0.08424031 | -0.56275954 | 0.94989632 | 0.03075817 | 1.04954079 | -0.12250206 | 0.6410357 | 1.30881507 |
| Magel2 | -1.69682638 | 0.08391192 | -0.83587919 | -1.67880532 | 1.54636073 | 0.09797124 | -0.49122068 | 0.79554962 | -0.30207019 | 0.96321526 | -0.0892309 | 0.84459633 | 0.76242753 |
| Tmem181a | -2.53492957 | -1.2088164 | -0.45453664 | -0.40036977 | 0.14993242 | 0.60115 | 0.63660582 | 0.68887372 | 0.07735457 | 0.17261794 | 0.2283996 | 1.41299927 | 0.63071903 |
| Zdhhc1 | -2.67701217 | -0.48010648 | -0.61670073 | -0.90506426 | 0.88364084 | -0.04172279 | 0.41627137 | 0.36854102 | 0.10030746 | 0.58114172 | 0.57980025 | 1.00579781 | 0.78510597 |
| Ubr3 | -2.07284767 | -0.46955272 | -1.04329283 | -1.37682777 | 1.00357636 | 0.74280827 | 0.13052596 | 0.59705821 | 0.30803225 | -0.13012034 | 0.65348082 | 1.40055958 | 0.2565999 |
| Pskh1 | -1.75475178 | -1.27004376 | -1.03748782 | -1.33605026 | 0.08003423 | 1.13332927 | 0.48811281 | 0.45890149 | 0.51649575 | 0.05911561 | 1.09972922 | 0.72568254 | 0.83693269 |
| Chd8 | -1.59681635 | -0.31211002 | -1.39181145 | -0.86677162 | 0.52077259 | 0.62100737 | 1.2139886 | -0.8639738 | 1.54380814 | 0.55561827 | 0.42660359 | -0.59672703 | 0.7464117 |
| Usp34 | -1.82244185 | -1.14903482 | -1.15956857 | -0.58133475 | 0.0411161 | 0.59974193 | 0.94301767 | 0.56369536 | 1.87675367 | 0.72920017 | -0.17166308 | 0.15339729 | -0.02287911 |
| Ajm1 | -1.49889359 | -0.76019368 | -0.34284784 | -2.03972189 | 0.64691643 | -0.46450541 | 0.35531065 | 1.19860134 | 0.14782823 | 1.29506405 | 0.61664891 | 0.78571444 | 0.06007836 |
| Tmem201 | -1.36002399 | -1.55108914 | 0.21056413 | -1.84272331 | 0.40354113 | -0.14023906 | 0.31615694 | 1.35253204 | 0.38675638 | 0.78195814 | 0.70736009 | 0.94250265 | -0.207296 |
| Usp35 | -2.12415507 | -1.09868159 | -0.05752088 | -1.52310055 | 0.97181612 | -0.19700488 | 0.55854361 | 0.45304329 | 0.76491161 | 0.99041439 | 0.81008621 | 0.48371966 | -0.03207193 |
| Pofut1 | -2.13558378 | -0.84984295 | 0.33716765 | -1.89140619 | 0.57273182 | 0.52382232 | 0.46021407 | 0.25622773 | 0.68375077 | 0.14253708 | 1.06479545 | 0.71390532 | 0.12168072 |
| Cul9 | -1.73038066 | -1.48616208 | -0.58107667 | -0.82700897 | 0.88481869 | -0.14958618 | 0.62446073 | 1.11489293 | 1.21752992 | 0.46632559 | 0.366445 | 0.9153761 | -0.81563441 |
| Ube2o | -1.72316506 | -1.5565139 | -0.276864 | -0.63831286 | 0.50304963 | -0.57931128 | 0.12379471 | 1.4823148 | 1.67170229 | 0.10714424 | 0.57716667 | 0.41497601 | -0.10598125 |
| Zfyve28 | -1.40260516 | -1.141187 | -0.68767723 | -1.21921305 | 0.19108582 | -0.66443675 | 0.49016908 | 1.48406938 | 1.58236772 | 0.45725495 | 1.05745956 | -0.00939518 | -0.13789214 |
| Mtmr4 | -1.6615579 | -1.26015815 | -1.11250523 | -0.77789097 | 0.8234196 | -0.14338453 | 0.28159157 | 1.3003302 | 0.47608805 | 1.75009909 | 0.10655845 | 0.19593535 | 0.02147445 |
| Dnajc10 | -1.31251936 | -1.50846146 | -0.94291142 | -1.12164294 | 0.04264652 | 0.02949902 | 0.24498057 | 1.96368064 | 0.44960397 | 0.76097022 | 0.97627474 | 0.44798229 | -0.03010277 |
| Susd2 | -1.64489334 | -0.79328542 | -1.35736106 | -0.71934057 | 2.22525418 | 0.15206736 | 0.27853193 | 0.61959162 | 0.10492152 | 0.62284309 | 0.35710555 | -0.32915202 | 0.48371717 |
| Serpina3n | -0.47618286 | -0.60462104 | -1.94733547 | -0.83639801 | 2.10816119 | 0.55900348 | 0.50121072 | 1.14165682 | -0.53933431 | 0.48394923 | -0.26829733 | -0.05403964 | -0.06777278 |
| Nlgn2 | -0.17007756 | -1.10775455 | -1.55522596 | -1.16241932 | 1.06283964 | 0.01279307 | 0.59291231 | 1.33969955 | -0.56211265 | -0.176879 | -0.33084984 | 1.77866367 | 0.27841062 |
| Ptprn | -1.62889679 | -0.70820507 | -1.2165036 | -1.09782838 | 1.36751584 | -0.15077669 | 0.18731467 | 1.43631732 | 0.16876939 | 0.87555652 | 0.00170977 | 1.11783208 | -0.35280507 |
| Megf8 | -2.097069 | -1.06628823 | 0.09383742 | -1.35907554 | 0.73049024 | -0.31130048 | 0.76055263 | 1.38391131 | 0.65475035 | 0.11758056 | 0.40406999 | 0.91102626 | -0.22248551 |
| Lztr1 | -0.49244224 | -1.90655776 | -0.26831374 | -1.69053589 | 1.25086767 | 0.91231824 | -0.67420795 | 1.28043971 | 0.17092725 | 0.62184569 | 0.46805464 | 0.08345317 | 0.24415121 |
| Avp | -0.65266178 | -1.74563853 | -1.03298444 | -1.12846112 | 1.22786824 | -0.12567266 | 0.44533048 | 1.83929656 | -0.5053189 | 0.49628285 | 0.12149346 | 0.56774191 | 0.49272393 |
| Bsdc1 | -1.17875955 | -1.26749307 | -1.12255333 | -1.30027741 | 1.10478541 | -0.36860698 | 0.62502501 | 1.75023036 | 0.67707499 | 0.16170534 | 0.56477581 | -0.25683891 | 0.61093232 |
| Mri1 | -1.95519875 | -0.60673288 | -0.47261305 | -1.55546611 | -0.41565895 | 0.26972327 | 0.68907852 | 0.43916789 | 0.60552884 | 0.17188112 | 0.09818574 | 1.60581099 | 1.12629339 |
| Ttn | -1.70414388 | -1.28053002 | -0.25632354 | -0.96196799 | 0.42630476 | 0.29840289 | 1.23427033 | -0.87651782 | -0.40224462 | 0.68479062 | 0.45654834 | 1.52821931 | 0.85319163 |
| Ltbp1 | -1.74741067 | -0.42111778 | 0.25016925 | -1.79971382 | 0.49929545 | 1.08128314 | -0.16607638 | -1.02683096 | 1.30474645 | 0.72713197 | 0.77144019 | 0.23101144 | 0.2960717 |
| Prkn | -1.7144254 | -0.41556768 | 0.16864306 | -2.19233559 | 0.48389142 | 0.31925053 | 0.04446513 | -0.58078049 | 0.65515903 | 1.16928248 | 1.00834845 | 0.59473263 | 0.45933642 |
| Dpysl5 | -2.24426238 | 0.00034885 | 0.17949198 | -1.70964017 | 0.78535494 | -0.22639768 | 0.02391519 | -0.49141497 | 0.7371552 | 1.17958125 | 0.87528871 | 0.42804358 | 0.46253551 |
| Trrap | -2.18790539 | -0.00796114 | -0.09469443 | -1.81226239 | 0.97466169 | 0.28433999 | 0.48590465 | -0.73484203 | 0.81519955 | 0.56112272 | 0.88747303 | 0.44784358 | 0.38112017 |
| Mast1 | -2.33302297 | -0.12743453 | -0.62992804 | -0.90023458 | 0.30923885 | 0.27253302 | 0.63004804 | -0.46090077 | 0.99858026 | 1.4915164 | 0.96897027 | 0.32013783 | -0.53950377 |
| Tmem131l | -2.34934049 | 0.1405811 | -0.52771062 | -1.13099415 | 0.28833645 | 0.87564959 | 0.76400251 | -1.05806155 | 0.76007897 | 0.60688445 | 1.13075853 | 0.26670821 | 0.23310698 |
| [illegible] | [illegible] | [illegible] | [illegible] | [illegible] | [illegible] | [illegible] | [illegible] | [illegible] | [illegible] | [illegible] | [illegible] | [illegible] | [illegible] |

**Supplemental table S3**

Gene information

| Ensembl_Gene_ID | Gene_Symbol | Gene_Title | Gene_Type | Normalized counts per sample: M_NC Midaged_01 | M_NC Midaged_02 | M_NC Midaged_03 | M_NC Midaged_04 | M_HFD Midaged_05 | M_HFD Midaged_06 | M_HFD Midaged_07 | M_HFD Midaged_08 | M_Reverse Midaged_09 | M_Reverse Midaged_10 | M_Reverse Midaged_11 | M_Reverse Midaged_12 | DESeq2 results for differential expression: M_HFD.vs.NC linearFC | M_HFD.vs.NC pvalue | M_HFD.vs.NC padj | M_HFD.vs.NC pass | M_Reverse.vs.NC linearFC | M_Reverse.vs.NC pvalue | M_Reverse.vs.NC padj | M_Reverse.vs.NC pass | M_Reverse.vs.HFD linearFC | M_Reverse.vs.HFD pvalue | M_Reverse.vs.HFD padj | M_Reverse.vs.HFD pass | Expression pattern: Pattern No. | Clustering | z-score expression values for heatmap visualization: Midaged_01 | Midaged_02 | Midaged_03 | Midaged_04 | Midaged_05 | Midaged_06 | Midaged_07 | Midaged_08 | Midaged_09 | Midaged_10 | Midaged_11 | Midaged_12 |
|---|---|---|---|---|---|---|---|---|---|---|---|---|---|---|---|---|---|---|---|---|---|---|---|---|---|---|---|---|---|---|---|---|---|---|---|---|---|---|---|---|---|
| [illegible] | [illegible] | [illegible] | [illegible] | [illegible] | [illegible] | [illegible] | [illegible] | [illegible] | [illegible] | [illegible] | [illegible] | [illegible] | [illegible] | [illegible] | [illegible] | [illegible] | [illegible] | [illegible] | [illegible] | [illegible] | [illegible] | [illegible] | [illegible] | [illegible] | [illegible] | [illegible] | [illegible] | [illegible] | [illegible] | [illegible] | [illegible] | [illegible] | [illegible] | [illegible] | [illegible] | [illegible] | [illegible] | [illegible] | [illegible] | [illegible] | [illegible] |

| | | | | | | | |
|---|---|---|---|---|---|---|---|
| ENSMUSG00000111312 | Gm47205 | predicted gene, | TEC | 15 | 17 | 18 | 25 |
| ENSMUSG00000105852 | Gm42890 | predicted gene | lncRNA | 48 | 55 | 60 | 84 |
| ENSMUSG00000030243 | Recql | RecQ protein-lil | protein_coding | 446 | 462 | 533 | 476 |
| ENSMUSG00000106995 | Gm33167 | predicted gene, | TEC | 45 | 49 | 57 | 61 |
| ENSMUSG00000097352 | C920009B18Rik | RIKEN cDNA C9 | transcribed_unproces | 8 | 1 | 9 | 2 |
| ENSMUSG00000038126 | Mphosph9 | M-phase phosp | protein_coding | 617 | 490 | 522 | 705 |
| ENSMUSG00000032198 | Dock6 | dedicator of cyt | protein_coding | 509 | 482 | 474 | 632 |
| ENSMUSG00000098024 | Gm27003 | predicted gene, | lncRNA | 12 | 4 | 9 | 15 |
| ENSMUSG00000084964 | Gm15503 | predicted gene | lncRNA | 1 | 5 | 9 | 1 |
| ENSMUSG00000022369 | Mtbp | Mdm2, transfor | protein_coding | 36 | 56 | 48 | 52 |
| ENSMUSG00000022160 | Mettl3 | methyltransfera | protein_coding | 428 | 473 | 435 | 432 |
| ENSMUSG00000073274 | Gm14636 | predicted gene | lncRNA | 0 | 2 | 1 | 2 |
| ENSMUSG00000045455 | Gm9797 | predicted pseu | transcribed_processe | 0 | 3 | 0 | 0 |
| ENSMUSG00000106951 | 5930430L01Rik | RIKEN cDNA 59 | lncRNA | 96 | 104 | 83 | 103 |
| ENSMUSG00000098120 | Gm5914 | predicted gene | lncRNA | 17 | 17 | 16 | 17 |
| ENSMUSG00000100213 | Gm28151 | predicted gene | lncRNA | 48 | 44 | 41 | 41 |
| ENSMUSG00000104876 | Trdc | T cell receptor ( | TR_C_gene | 9 | 20 | 12 | 19 |
| ENSMUSG00000107096 | Gm43597 | predicted gene | lncRNA | 373 | 419 | 445 | 451 |
| ENSMUSG00000021481 | Zfp346 | zinc finger prot | protein_coding | 445 | 553 | 486 | 500 |
| ENSMUSG00000035158 | Mitf | melanogenesis | protein_coding | 137 | 157 | 141 | 154 |
| ENSMUSG00000091784 | Gm17022 | predicted gene | processed_pseudoge | 12 | 15 | 18 | 19 |
| ENSMUSG00000118310 | Gm50268 | predicted gene, | TEC | 5 | 8 | 6 | 13 |
| ENSMUSG00000052783 | Grk4 | G protein-coupl | protein_coding | 144 | 172 | 165 | 206 |
| ENSMUSG00000100183 | Gm28512 | predicted gene | lncRNA | 48 | 67 | 53 | 67 |
| ENSMUSG00000086907 | Gm15298 | predicted gene | lncRNA | 61 | 87 | 60 | 85 |
| ENSMUSG00000024006 | Stk38 | serine/threonin | protein_coding | 983 | 1,032 | 982 | 1,288 |
| ENSMUSG00000026305 | Lrrfip1 | leucine rich rep | protein_coding | 1,002 | 1,065 | 985 | 1,092 |
| ENSMUSG00000001768 | Rin2 | Ras and Rab int | protein_coding | 1,012 | 1,119 | 1,047 | 963 |
| ENSMUSG00000002205 | Vrk3 | vaccinia related | protein_coding | 456 | 470 | 441 | 380 |
| ENSMUSG00000079224 | Gm6565 | predicted gene | processed_pseudoge | 0 | 0 | 0 | 0 |
| ENSMUSG00000041498 | Kif14 | kinesin family n | protein_coding | 6 | 4 | 2 | 5 |
| ENSMUSG00000021209 | Ppp4r4 | protein phosph | protein_coding | 916 | 933 | 936 | 887 |
| ENSMUSG00000055296 | Tmem245 | transmembrane | protein_coding | 3,800 | 3,843 | 3,804 | 4,002 |
| ENSMUSG00000075113 | Or4c29 | olfactory recep | protein_coding | 4 | 7 | 6 | 13 |
| ENSMUSG00000033021 | Gmppa | GDP-mannose p | protein_coding | 479 | 476 | 433 | 456 |
| ENSMUSG00000073102 | Drc1 | dynein regulato | protein_coding | 483 | 394 | 420 | 434 |
| ENSMUSG00000005907 | Pex1 | peroxisomal bio | protein_coding | 796 | 718 | 727 | 786 |
| ENSMUSG00000036550 | Cnot1 | CCR4-NOT tran | protein_coding | 4,394 | 4,319 | 3,897 | 4,154 |
| ENSMUSG00000039254 | Pomt1 | protein-O-man | protein_coding | 398 | 415 | 316 | 362 |
| ENSMUSG00000029521 | Chek2 | checkpoint kina | protein_coding | 43 | 57 | 26 | 102 |
| ENSMUSG00000116908 | Gm49599 | predicted gene, | TEC | 60 | 69 | 57 | 92 |
| ENSMUSG00000056459 | Zbtb25 | zinc finger and | protein_coding | 323 | 326 | 366 | 414 |
| ENSMUSG00000070733 | Fryl | FRY like transcr | protein_coding | 2,547 | 2,511 | 2,693 | 2,840 |
| ENSMUSG00000083734 | Gm16209 | predicted gene | processed_pseudoge | 7 | 4 | 11 | 17 |
| ENSMUSG00000028017 | Egf | epidermal grow | protein_coding | 39 | 54 | 40 | 58 |
| ENSMUSG00000097207 | 6030443J06Rik | RIKEN cDNA 60 | lncRNA | 22 | 23 | 21 | 28 |
| ENSMUSG00000111854 | Gm48740 | predicted gene, | lncRNA | 10 | 25 | 15 | 23 |
| ENSMUSG00000055072 | Gm9964 | predicted gene | TEC | 46 | 53 | 35 | 59 |
| ENSMUSG00000034761 | Map4k5 | mitogen-activat | protein_coding | 1,134 | 1,133 | 1,082 | 1,401 |
| ENSMUSG00000087620 | 5330434G04Rik | RIKEN cDNA 53 | lncRNA | 412 | 485 | 430 | 541 |
| ENSMUSG00000108465 | Gm45110 | predicted gene | TEC | 46 | 65 | 50 | 76 |
| ENSMUSG00000091511 | Vmn2r87 | vomeronasal 2, | protein_coding | 41 | 32 | 26 | 36 |
| ENSMUSG00000115591 | Gm31282 | predicted gene, | lncRNA | 50 | 25 | 39 | 51 |
| ENSMUSG00000107571 | Gm43877 | predicted gene, | TEC | 4 | 0 | 0 | 3 |
| ENSMUSG00000110189 | Gm45665 | predicted gene | processed_pseudoge | 53 | 25 | 21 | 54 |
| ENSMUSG00000025878 | Uimc1 | ubiquitin intera | protein_coding | 416 | 387 | 374 | 414 |
| ENSMUSG00000009555 | Cdk9 | cyclin-depende | protein_coding | 1,414 | 1,325 | 1,388 | 1,397 |
| ENSMUSG00000027668 | Mfn1 | mitofusin 1 | protein_coding | 2,386 | 2,402 | 2,359 | 2,672 |
| ENSMUSG00000023170 | Gps2 | G protein pathw | protein_coding | 382 | 334 | 317 | 389 |
| ENSMUSG00000037364 | Srrt | serrate RNA eff | protein_coding | 3,046 | 2,920 | 2,890 | 3,369 |
| ENSMUSG00000042359 | Osbpl6 | oxysterol bindir | protein_coding | 2,405 | 2,393 | 2,350 | 2,511 |
| ENSMUSG00000021500 | Ddx46 | DEAD box helic | protein_coding | 2,190 | 2,100 | 2,075 | 2,512 |
| ENSMUSG00000031256 | Cstf2 | cleavage stimul | protein_coding | 3,887 | 3,941 | 3,641 | 4,656 |
| ENSMUSG00000037674 | Rfx7 | regulatory facto | protein_coding | 1,729 | 1,730 | 1,731 | 1,969 |
| ENSMUSG00000040867 | Begain | brain-enriched | protein_coding | 3,628 | 3,804 | 3,386 | 4,210 |
| ENSMUSG00000040840 | 4930579G18Rik | RIKEN cDNA 49 | lncRNA | 49 | 44 | 29 | 61 |
| ENSMUSG00000118501 | Gm53048 | predicted gene, | unprocessed_pseudo | 5 | 4 | 2 | 9 |
| ENSMUSG00000028820 | Sfpq | splicing factor p | protein_coding | 5,088 | 5,071 | 4,435 | 5,915 |
| ENSMUSG00000039860 | Srrm3 | serine/arginine | protein_coding | 1,185 | 1,110 | 1,043 | 1,409 |
| ENSMUSG00000094030 | Gm21833 | predicted gene, | protein_coding | 166 | 131 | 123 | 265 |
| ENSMUSG00000052752 | Traf7 | TNF receptor-a | protein_coding | 714 | 686 | 653 | 895 |
| ENSMUSG00000060445 | Sycp2 | synaptonemal c | protein_coding | 27 | 27 | 18 | 70 |
| ENSMUSG00000066760 | Psg16 | pregnancy spec | protein_coding | 1,176 | 1,133 | 1,212 | 1,452 |
| ENSMUSG00000074292 | Gm10660 | predicted gene | lncRNA | 15 | 18 | 24 | 67 |
| ENSMUSG00000028637 | Ccdc30 | coiled-coil dom | protein_coding | 519 | 489 | 537 | 629 |
| ENSMUSG00000052676 | Zmat1 | zinc finger, mat | protein_coding | 1,003 | 902 | 990 | 1,313 |
| ENSMUSG00000111436 | Gm47356 | predicted gene, | processed_pseudoge | 58 | 58 | 62 | 121 |
| ENSMUSG00000041685 | Fcho2 | FCH domain on | protein_coding | 1,382 | 1,416 | 1,430 | 1,646 |
| ENSMUSG00000033981 | Gria2 | glutamate rece | protein_coding | 11,724 | 11,633 | 11,714 | 13,735 |
| ENSMUSG00000111939 | Gm48177 | predicted gene, | processed_pseudoge | 123 | 135 | 135 | 284 |
| ENSMUSG00000000838 | Fmr1 | fragile X messe | protein_coding | 1,434 | 1,354 | 1,308 | 1,585 |
| ENSMUSG00000078868 | Gm14412 | predicted gene | protein_coding | 78 | 62 | 44 | 100 |
| ENSMUSG00000013236 | Ptprs | protein tyrosine | protein_coding | 8,842 | 8,729 | 7,441 | 10,645 |
| ENSMUSG00000028180 | Zranb2 | zinc finger, RAN | protein_coding | 4,289 | 3,883 | 3,716 | 4,958 |
| ENSMUSG00000020069 | Hnrnph3 | heterogeneous | protein_coding | 1,550 | 1,441 | 1,396 | 1,744 |
| ENSMUSG00000112158 | Gm47822 | predicted gene, | TEC | 72 | 65 | 53 | 142 |
| ENSMUSG00000018537 | Pcgf2 | polycomb grou | protein_coding | 751 | 669 | 674 | 844 |
| ENSMUSG00000021493 | Pdlim7 | PDZ and LIM do | protein_coding | 775 | 698 | 679 | 806 |
| ENSMUSG00000035299 | Mid1 | midline 1 | protein_coding | 364 | 304 | 317 | 392 |
| ENSMUSG00000004980 | Hnrnpa2b1 | heterogeneous | protein_coding | 12,487 | 12,713 | 12,800 | 14,119 |
| ENSMUSG00000037519 | Ppfia1 | protein tyrosine | protein_coding | 1,220 | 1,289 | 1,229 | 1,505 |
| ENSMUSG00000025261 | Huwe1 | HECT, UBA and | protein_coding | 17,042 | 16,487 | 17,072 | 22,030 |
| ENSMUSG00000096808 | | | protein_coding | 275 | 246 | 275 | 383 |
| ENSMUSG00000018379 | Srsf1 | serine and argir | protein_coding | 5,642 | 5,640 | 5,429 | 6,522 |
| ENSMUSG00000020464 | Pnpt1 | polyribonucleot | protein_coding | 763 | 739 | 702 | 885 |
| ENSMUSG00000020982 | Nemf | nuclear export | protein_coding | 2,797 | 2,682 | 2,631 | 3,122 |
| ENSMUSG00000104046 | Gm37567 | predicted gene, | TEC | 871 | 879 | 833 | 1,075 |
| ENSMUSG00000026116 | Tmem131 | transmembrane | protein_coding | 3,093 | 3,090 | 2,983 | 3,264 |
| ENSMUSG00000117286 | Gm1043 | predicted gene | protein_coding | 1,548 | 1,622 | 1,502 | 1,999 |
| ENSMUSG00000025747 | Tyms | thymidylate syn | protein_coding | 254 | 273 | 251 | 354 |
| ENSMUSG00000036057 | Ptpn23 | protein tyrosine | protein_coding | 2,177 | 2,142 | 1,989 | 2,323 |
| ENSMUSG00000025241 | Fyco1 | FYVE and coilec | protein_coding | 2,765 | 2,783 | 2,650 | 3,139 |
| ENSMUSG00000040225 | Prrc2c | proline-rich coil | protein_coding | 9,696 | 9,621 | 9,145 | 11,307 |
| ENSMUSG00000020640 | Itsn2 | intersectin 2 | protein_coding | 2,040 | 2,016 | 2,104 | 2,337 |
| ENSMUSG00000019996 | Map7 | microtubule-as | protein_coding | 2,264 | 1,992 | 1,978 | 2,579 |
| ENSMUSG00000034243 | Golgb1 | golgi autoantig | protein_coding | 7,363 | 6,832 | 7,086 | 7,894 |
| ENSMUSG00000111485 | Gm48443 | predicted gene, | processed_pseudoge | 742 | 645 | 730 | 1,042 |
| ENSMUSG00000029004 | Kmt2e | lysine (K)-specif | protein_coding | 4,475 | 3,957 | 4,084 | 5,186 |
| ENSMUSG00000053510 | Nrd1 | nardilysin, N-ar | protein_coding | 3,790 | 3,144 | 3,330 | 4,311 |
| ENSMUSG00000071054 | Safb | scaffold attachr | protein_coding | 3,442 | 3,264 | 3,396 | 4,130 |
| ENSMUSG00000104837 | Gm42520 | predicted gene | TEC | 81 | 83 | 80 | 136 |
| ENSMUSG00000069892 | 9930111J21Rik2 | RIKEN cDNA 99 | protein_coding | 469 | 419 | 402 | 553 |
| ENSMUSG00000102326 | Gm37788 | predicted gene, | TEC | 151 | 139 | 125 | 220 |
| ENSMUSG00000106631 | Gm42669 | predicted gene | protein_coding | 511 | 503 | 485 | 623 |
| ENSMUSG00000025272 | Tro | trophinin | protein_coding | 11,335 | 11,292 | 10,583 | 13,784 |
| ENSMUSG00000002028 | Kmt2a | lysine (K)-specif | protein_coding | 8,639 | 8,286 | 7,837 | 10,298 |
| ENSMUSG00000037541 | Shank2 | SH3 and multip | protein_coding | 2,666 | 2,683 | 2,512 | 3,094 |
| ENSMUSG00000038346 | Zfp384 | zinc finger prot | protein_coding | 858 | 846 | 815 | 1,108 |
| ENSMUSG00000040721 | Zfhx2 | zinc finger hom | protein_coding | 2,875 | 2,773 | 2,553 | 3,254 |
| ENSMUSG00000112122 | Gm47628 | predicted gene, | processed_pseudoge | 147 | 131 | 93 | 245 |
| ENSMUSG00000010608 | Rbm25 | RNA binding mo | protein_coding | 3,715 | 3,841 | 3,228 | 4,755 |
| ENSMUSG00000063511 | Snrnp70 | small nuclear ri | protein_coding | 6,884 | 6,983 | 6,008 | 8,935 |
| ENSMUSG00000028809 | Srrm1 | serine/arginine | protein_coding | 4,745 | 4,541 | 3,839 | 6,287 |
| ENSMUSG00000113795 | Gm48119 | predicted gene, | processed_pseudoge | 750 | 702 | 652 | 1,143 |
| ENSMUSG00000113393 | Gm48415 | predicted gene, | processed_pseudoge | 992 | 1,002 | 818 | 1,738 |
| ENSMUSG00000061331 | Gm17132 | predicted gene | unprocessed_pseudo | 552 | 568 | 478 | 831 |
| ENSMUSG00000112049 | Gm48269 | predicted gene, | processed_pseudoge | 289 | 359 | 280 | 597 |
| ENSMUSG00000022185 | Acin1 | apoptotic chror | protein_coding | 3,191 | 3,366 | 3,027 | 4,134 |
| ENSMUSG00000020863 | Luc7l3 | LUC7-like 3 (S. c | protein_coding | 3,881 | 4,007 | 3,778 | 4,754 |
| ENSMUSG00000020994 | Pnn | pinin | protein_coding | 2,290 | 2,352 | 2,193 | 3,025 |
| ENSMUSG00000108847 | B830042I05Rik | RIKEN cDNA B8 | lncRNA | 163 | 183 | 158 | 325 |
| ENSMUSG00000111472 | Gm47103 | predicted gene, | processed_pseudoge | 35 | 28 | 27 | 108 |
| ENSMUSG00000026339 | Ccdc93 | coiled-coil dom | protein_coding | 1,095 | 1,119 | 943 | 1,105 |
| ENSMUSG00000042909 | Or52h1 | olfactory recep | protein_coding | 24 | 38 | 20 | 35 |
| ENSMUSG00000014633 | Cmc2 | COX assembly r | protein_coding | 181 | 198 | 162 | 216 |
| ENSMUSG00000066838 | Zfp772 | zinc finger prot | protein_coding | 295 | 344 | 255 | 353 |
| ENSMUSG00000035051 | Dhx57 | DEAH (Asp-Glu- | protein_coding | 1,477 | 1,480 | 1,371 | 1,678 |
| ENSMUSG00000039205 | Ciz1 | CDKN1A interac | protein_coding | 1,067 | 1,009 | 880 | 1,276 |
| ENSMUSG00000113161 | Gm47457 | predicted gene, | TEC | 635 | 607 | 472 | 832 |
| ENSMUSG00000100750 | Gm29084 | predicted gene | lncRNA | 1,394 | 1,443 | 1,165 | 1,700 |
| ENSMUSG00000108173 | Gm44231 | predicted gene, | TEC | 850 | 936 | 747 | 1,089 |
| ENSMUSG00000030138 | Bms1 | BMS1, ribosom | protein_coding | 871 | 958 | 781 | 1,018 |
| ENSMUSG00000034799 | Unc13a | unc-13 homolo | protein_coding | 4,191 | 4,777 | 3,889 | 4,978 |
| ENSMUSG00000066129 | Kndc1 | kinase non-cata | protein_coding | 4,928 | 5,170 | 4,939 | 5,329 |
| ENSMUSG00000027411 | Vps16 | VSP16 CORVET/ | protein_coding | 930 | 923 | 985 | 1,093 |
| ENSMUSG00000031161 | Hdac6 | histone deacety | protein_coding | 1,154 | 1,129 | 1,319 | 1,505 |
| ENSMUSG00000003810 | Mast2 | microtubule as | protein_coding | 3,566 | 3,490 | 3,599 | 3,733 |
| ENSMUSG00000105694 | Gm43237 | predicted gene | TEC | 1 | 1 | 2 | 5 |
| ENSMUSG00000110170 | St6galnac2 | ST6 (alpha-N-ac | protein_coding | 21 | 8 | 14 | 21 |
| ENSMUSG00000019977 | Hbs1l | Hbs1-like (S. ce | protein_coding | 1,003 | 907 | 1,019 | 1,277 |
| ENSMUSG00000073436 | Eme2 | essential meiot | protein_coding | 413 | 425 | 445 | 520 |
| ENSMUSG00000111325 | Gm47140 | predicted gene, | lncRNA | 126 | 138 | 117 | 229 |
| ENSMUSG00000108803 | 4930533N22Rik | RIKEN cDNA 49 | lncRNA | 495 | 519 | 503 | 782 |
| ENSMUSG00000109887 | Gm28756 | predicted gene | lncRNA | 257 | 260 | 231 | 377 |
| ENSMUSG00000040250 | Ints13 | integrator comp | protein_coding | 532 | 558 | 547 | 723 |
| ENSMUSG00000086130 | Gm16211 | predicted gene | lncRNA | 255 | 312 | 295 | 393 |
| ENSMUSG00000055436 | Srsf11 | serine and argir | protein_coding | 2,967 | 3,315 | 2,933 | 3,914 |
| ENSMUSG00000097397 | Gm16861 | predicted gene, | lncRNA | 45 | 55 | 39 | 84 |
| ENSMUSG00000033004 | Mycbp2 | MYC binding pr | protein_coding | 15,070 | 16,360 | 14,872 | 18,191 |
| ENSMUSG00000091076 | Vmn2r115 | vomeronasal 2, | protein_coding | 278 | 357 | 288 | 571 |
| ENSMUSG00000021134 | Srsf5 | serine and argir | protein_coding | 6,516 | 7,117 | 6,635 | 7,961 |
| ENSMUSG00000103694 | Gm37530 | predicted gene, | TEC | 142 | 197 | 150 | 243 |
| ENSMUSG00000103234 | Gm37158 | predicted gene, | TEC | 236 | 285 | 215 | 438 |
| ENSMUSG00000117599 | Gm49971 | predicted gene, | TEC | 231 | 297 | 185 | 455 |
| ENSMUSG00000094472 | Gm21897 | predicted gene, | protein_coding | 400 | 575 | 391 | 853 |
| ENSMUSG00000111269 | Gm47933 | predicted gene, | processed_pseudoge | 169 | 219 | 138 | 385 |
| ENSMUSG00000108088 | Gm44229 | predicted gene, | TEC | 8 | 11 | 6 | 16 |
| ENSMUSG00000105471 | A430073D23Rik | RIKEN cDNA A4 | lncRNA | 17 | 24 | 11 | 36 |
| ENSMUSG00000111544 | 4930534H03Rik | RIKEN cDNA 49 | lncRNA | 64 | 75 | 47 | 121 |
| ENSMUSG00000090362 | Vmn2r79 | vomeronasal 2, | protein_coding | 31 | 44 | 25 | 51 |
| ENSMUSG00000102289 | Gm31258 | predicted gene, | TEC | 27 | 45 | 29 | 64 |
| ENSMUSG00000009079 | Ewsr1 | Ewing sarcoma | protein_coding | 3,997 | 4,620 | 3,715 | 5,248 |
| ENSMUSG00000097391 | Mirg | miRNA containi | lncRNA | 538 | 654 | 498 | 786 |
| ENSMUSG00000114995 | Gm49284 | predicted gene, | TEC | 163 | 157 | 159 | 181 |
| ENSMUSG00000034820 | Cpsf7 | cleavage and po | protein_coding | 2,459 | 2,474 | 2,406 | 2,789 |
| ENSMUSG00000072739 | Gm10408 | predicted gene | protein_coding | 0 | 0 | 0 | 11 |
| ENSMUSG00000103133 | Gm37303 | predicted gene, | TEC | 33 | 35 | 30 | 63 |
| ENSMUSG00000027201 | Myef2 | myelin basic pr | protein_coding | 2,747 | 2,871 | 2,747 | 3,157 |
| ENSMUSG00000085687 | Gm16153 | predicted gene | lncRNA | 58 | 59 | 59 | 100 |
| ENSMUSG00000097911 | Gm26691 | predicted gene, | lncRNA | 147 | 147 | 150 | 212 |
| ENSMUSG00000110317 | Gm45250 | predicted gene | lncRNA | 108 | 117 | 93 | 140 |
| ENSMUSG00000038943 | Prc1 | protein regulato | protein_coding | 60 | 81 | 56 | 86 |
| ENSMUSG00000091382 | Vmn1r18 | vomeronasal 1 | protein_coding | 100 | 108 | 90 | 148 |
| ENSMUSG00000045903 | Npas4 | neuronal PAS d | protein_coding | 81 | 118 | 110 | 166 |
| ENSMUSG00000114970 | Gm49069 | predicted gene, | TEC | 32 | 75 | 44 | 81 |
| ENSMUSG00000046650 | Or5an6 | olfactory recep | protein_coding | 14 | 18 | 15 | 24 |
| ENSMUSG00000105791 | Gm43341 | predicted gene | TEC | 3 | 6 | 2 | 9 |
| ENSMUSG00000109095 | Gm44799 | predicted gene | TEC | 7 | 14 | 5 | 19 |
| ENSMUSG00000001082 | Mfsd10 | major facilitato | protein_coding | 252 | 268 | 215 | 319 |
| ENSMUSG00000027981 | Rnpc3 | RNA-binding re | protein_coding | 1,149 | 1,316 | 1,046 | 1,635 |
| ENSMUSG00000108713 | Gm33027 | predicted gene, | lncRNA | 105 | 125 | 87 | 163 |
| ENSMUSG00000105476 | Gm35439 | predicted gene, | lncRNA | 70 | 89 | 71 | 145 |
| ENSMUSG00000021767 | Kat6b | K(lysine) acetyl | protein_coding | 1,615 | 1,757 | 1,592 | 1,838 |
| ENSMUSG00000116718 | Gm49668 | predicted gene, | lncRNA | 102 | 137 | 99 | 186 |
| ENSMUSG00000021611 | Tert | telomerase rev | protein_coding | 84 | 114 | 85 | 210 |
| ENSMUSG00000109390 | Gm45718 | predicted gene | TEC | 53 | 67 | 54 | 100 |
| ENSMUSG00000115276 | 9930017N22Rik | RIKEN cDNA 99 | TEC | 44 | 56 | 51 | 88 |
| ENSMUSG00000035151 | Elmod2 | ELMO/CED-12 c | protein_coding | 791 | 883 | 832 | 1,097 |
| ENSMUSG00000051832 | E230016K23Rik | RIKEN cDNA E2 | lncRNA | 8 | 21 | 15 | 41 |
| ENSMUSG00000036403 | Cep135 | centrosomal pr | protein_coding | 356 | 384 | 365 | 494 |
| ENSMUSG00000064043 | Trerf1 | transcriptional | protein_coding | 1,085 | 1,098 | 1,063 | 1,445 |
| ENSMUSG00000033540 | Idua | iduronidase, alp | protein_coding | 461 | 533 | 484 | 677 |
| ENSMUSG00000110311 | Gm45410 | predicted gene | TEC | 100 | 111 | 105 | 167 |
| ENSMUSG00000104416 | Gm37067 | predicted gene, | TEC | 19 | 28 | 29 | 65 |
| ENSMUSG00000114722 | Gm31392 | predicted gene, | lncRNA | 60 | 82 | 75 | 129 |
| ENSMUSG00000021666 | Gfm2 | G elongation fa | protein_coding | 588 | 622 | 556 | 852 |
| ENSMUSG00000026171 | Rnf25 | ring finger prot | protein_coding | 854 | 929 | 906 | 1,154 |
| ENSMUSG00000002949 | Timm44 | translocase of i | protein_coding | 737 | 828 | 746 | 904 |
| ENSMUSG00000056211 | R3hdm1 | R3H domain co | protein_coding | 3,991 | 4,032 | 3,898 | 4,520 |
| ENSMUSG00000072647 | Adam1a | a disintegrin an | protein_coding | 467 | 620 | 441 | 737 |
| ENSMUSG00000111585 | Gm47355 | predicted gene, | processed_pseudoge | 45 | 66 | 38 | 104 |
| ENSMUSG00000002266 | Zim1 | zinc finger, imp | protein_coding | 2,814 | 2,973 | 2,652 | 3,482 |
| ENSMUSG00000034163 | Zfc3h1 | zinc finger, C3H | protein_coding | 1,285 | 1,460 | 1,254 | 1,749 |
| ENSMUSG00000069565 | Dazap1 | DAZ associated | protein_coding | 1,273 | 1,305 | 1,199 | 1,545 |
| ENSMUSG00000028307 | Aldob | aldolase B, fruc | protein_coding | 31 | 44 | 27 | 69 |
| ENSMUSG00000070495 | Ctcfl | CCCTC-binding | protein_coding | 113 | 143 | 104 | 180 |
| ENSMUSG00000040165 | Cd209c | CD209c antigen | protein_coding | 86 | 94 | 69 | 120 |
| ENSMUSG00000046056 | Sbsn | suprabasin | protein_coding | 152 | 179 | 125 | 252 |
| ENSMUSG00000113328 | Gm47260 | predicted gene, | TEC | 187 | 221 | 164 | 272 |
| ENSMUSG00000020070 | Rufy2 | RUN and FYVE c | protein_coding | 2,065 | 2,195 | 1,959 | 2,646 |
| ENSMUSG00000043411 | Usp48 | ubiquitin specif | protein_coding | 2,302 | 2,385 | 2,092 | 3,207 |
| ENSMUSG00000030231 | Plekha5 | pleckstrin homo | protein_coding | 1,042 | 1,098 | 995 | 1,361 |
| ENSMUSG00000054142 | Vmn1r236 | vomeronasal 1 | protein_coding | 58 | 72 | 50 | 113 |
| ENSMUSG00000101906 | Mrgprc2-ps | MAS-related GF | unprocessed_pseudo | 21 | 39 | 20 | 65 |
| ENSMUSG00000113388 | Gm48111 | predicted gene, | processed_pseudoge | 386 | 423 | 287 | 1,040 |
| ENSMUSG00000114367 | Gm48216 | predicted gene, | processed_pseudoge | 243 | 221 | 152 | 374 |
| ENSMUSG00000117670 | Gm50270 | predicted gene, | lncRNA | 13 | 8 | 2 | 44 |
| ENSMUSG00000028064 | Sema4a | sema domain, i | protein_coding | 1,482 | 1,535 | 1,425 | 1,916 |
| ENSMUSG00000106060 | Gm42488 | predicted gene | lncRNA | 29 | 32 | 27 | 51 |
| ENSMUSG00000031441 | Atp11a | ATPase, class VI | protein_coding | 3,202 | 3,317 | 2,951 | 3,951 |
| ENSMUSG00000038569 | Rad9b | RAD9 checkpoi | protein_coding | 281 | 292 | 248 | 380 |
| ENSMUSG00000032580 | Rbm5 | RNA binding mo | protein_coding | 3,713 | 4,239 | 3,952 | 5,588 |
| ENSMUSG00000075144 | Or5l13 | olfactory recep | protein_coding | 51 | 53 | 46 | 96 |
| ENSMUSG00000027751 | Supt20 | SPT20 SAGA co | protein_coding | 984 | 1,104 | 1,090 | 1,403 |
| ENSMUSG00000040407 | Akap9 | A kinase (PRKA) | protein_coding | 5,477 | 5,774 | 5,715 | 6,885 |
| ENSMUSG00000055341 | Zfp457 | zinc finger prot | protein_coding | 70 | 71 | 83 | 164 |
| ENSMUSG00000028649 | Macf1 | microtubule-ac | protein_coding | 23,367 | 25,820 | 23,001 | 29,488 |
| ENSMUSG00000083889 | E530001F21Rik | RIKEN cDNA E5 | processed_pseudoge | 36 | 58 | 39 | 109 |
| ENSMUSG00000113517 | Gm47905 | predicted gene, | TEC | 23 | 60 | 44 | 85 |
| ENSMUSG00000037965 | Zc3h7a | zinc finger CCCH | protein_coding | 726 | 1,135 | 939 | 1,384 |

| | | | | | | | | | | | | | | | | | | | | | | | | | | | | | | | | | | | | | | |
|---|---|---|---|---|---|---|---|---|---|---|---|---|---|---|---|---|---|---|---|---|---|---|---|---|---|---|---|---|---|---|---|---|---|---|---|---|---|---|
| ENSMUSG00000042308 | Setd1a | SET domain cor | protein_coding | 1,761 | 2,178 | 2,006 | 2,612 | 2,455 | 2,662 | 2,277 | 2,303 | 2,511 | 2,486 | 2,694 | 2,601 | 1.13 | 0.0286 | 0.213 | 1.2 | 0.00111 | 0.018 up | 1.06 | 0.287 | 1 | 3 | 477 | -2.28186755 | -0.63985916 | -1.27887606 | 0.78927708 | 0.30012633 | 0.93806949 | -0.29050556 | -0.20384749 | 0.47887404 | 0.39897645 | 1.03497026 | 0.75466218 |
| ENSMUSG00000108884 | Gm45792 | predicted gene | lncRNA | 70 | 120 | 102 | 197 | 179 | 183 | 141 | 174 | 181 | 174 | 208 | 203 | 1.38 | 0.0186 | 0.177 | 1.56 | 0.00093 | 0.016 up | 1.13 | 0.346 | 1 | 3 | 478 | -2.2692264 | -0.85600923 | -1.30714912 | 0.78466769 | 0.43178053 | 0.52445849 | -0.34996008 | 0.34667215 | 0.48427001 | 0.35134461 | 0.969516 | 0.88963534 |
| ENSMUSG00000027676 | Ccdc39 | coiled-coil dom | protein_coding | 658 | 826 | 666 | 931 | 983 | 1,047 | 796 | 989 | 1,006 | 875 | 1,194 | 976 | 1.24 | 0.0141 | 0.155 | 1.31 | 0.00159 | 0.0222 up | 1.06 | 0.489 | 1 | 3 | 479 | -1.72730736 | -0.49298974 | -1.66376269 | 0.1835121 | 0.49013429 | 0.85352293 | -0.69381082 | 0.52622794 | 0.62368346 | -0.17057864 | 1.61929509 | 0.45207344 |
| ENSMUSG00000026131 | Dst | dystonin | protein_coding | 25,427 | 26,744 | 26,413 | 30,098 | 30,060 | 30,120 | 27,727 | 29,500 | 30,099 | 28,701 | 32,158 | 30,755 | 1.08 | 0.049 | 0.27 | 1.12 | 0.00387 | 0.039 up | 1.04 | 0.358 | 1 | 3 | 480 | -1.82697882 | -1.11043841 | -1.28715812 | 0.56771285 | 0.54975713 | 0.57821193 | -0.59801014 | 0.28285991 | 0.56820054 | -0.10754526 | 1.50881027 | 0.87457814 |
| ENSMUSG00000114290 | 4930435F18Rik | RIKEN cDNA 49 | lncRNA | 33 | 51 | 42 | 84 | 89 | 88 | 63 | 74 | 80 | 79 | 107 | 88 | 1.49 | 0.0061 | 0.105 | 1.67 | 0.000211 | 0.00615 up | 1.13 | 0.351 | 1 | 3 | 481 | -1.98911272 | -0.97374523 | -1.44233178 | 0.52761291 | 0.72987373 | 0.67289862 | -0.37571537 | 0.10253684 | 0.36721247 | 0.33012651 | 1.38097032 | 0.6696737 |
| ENSMUSG00000044519 | Zfp488 | zinc finger prot | protein_coding | 481 | 550 | 545 | 662 | 762 | 779 | 647 | 715 | 717 | 781 | 826 | 756 | 1.3 | 0.0000435 | 0.0204 up | 1.38 | 0.000000333 | 0.000148 up | 1.06 | 0.332 | 1 | 3 | 482 | -1.96647736 | -1.21823866 | -1.26341478 | -0.14687799 | 0.69690148 | 0.834003 | -0.27736779 | 0.31678855 | 0.33438054 | 0.84799532 | 1.19003632 | 0.65227135 |
| ENSMUSG00000054702 | Ap1s3 | adaptor-relatec | protein_coding | 311 | 444 | 373 | 602 | 658 | 726 | 479 | 598 | 648 | 682 | 780 | 756 | 1.42 | 0.0017 | 0.063 | 1.65 | 0.00000579 | 0.000721 up | 1.16 | 0.165 | 1 | 3 | 483 | -1.99013924 | -0.8634643 | -1.42363144 | 0.18131845 | 0.49693701 | 0.85673908 | -0.61135778 | 0.15824975 | 0.44464143 | 0.62469926 | 1.12006991 | 1.00593788 |
| ENSMUSG00000066263 | Or51k1 | olfactory recep | protein_coding | 228 | 314 | 264 | 415 | 467 | 550 | 383 | 425 | 506 | 494 | 560 | 507 | 1.49 | 0.000166 | 0.0296 up | 1.69 | 0.000000575 | 0.000193 up | 1.13 | 0.223 | 1 | 3 | 484 | -1.95375939 | -0.96175004 | -1.50787787 | -0.00372227 | 0.41899423 | 1.03392829 | -0.28331525 | 0.07995145 | 0.72203717 | 0.63312567 | 1.09853479 | 0.72385322 |
| ENSMUSG00000059729 | Or2y1 | olfactory recep | protein_coding | 141 | 166 | 164 | 244 | 253 | 301 | 211 | 257 | 301 | 264 | 316 | 283 | 1.43 | 0.000417 | 0.0378 up | 1.62 | 0.000000791 | 0.000227 up | 1.14 | 0.166 | 1 | 3 | 485 | -1.83380797 | -1.28957781 | -1.3250656 | 0.10947305 | 0.25662058 | 0.949525 | -0.4323148 | 0.30763291 | 0.95775476 | 0.42739197 | 1.16479863 | 0.70756929 |
| ENSMUSG00000022961 | Son | Son DNA bindin | protein_coding | 10,461 | 11,557 | 10,778 | 13,085 | 13,368 | 14,423 | 12,647 | 13,068 | 13,444 | 13,428 | 14,568 | 13,831 | 1.17 | 0.000338 | 0.0348 up | 1.2 | 0.0000136 | 0.00111 up | 1.03 | 0.447 | 1 | 3 | 486 | -1.90617626 | -0.97466188 | -1.62704068 | 0.18919602 | 0.3897108 | 1.10273121 | -0.13061925 | 0.17685249 | 0.44263532 | 0.43148657 | 1.19680162 | 0.70908403 |
| ENSMUSG00000089719 | Gm15758 | predicted gene | lncRNA | 16 | 28 | 23 | 50 | 50 | 73 | 46 | 48 | 55 | 53 | 67 | 60 | 1.85 | 0.003 | 0.0792 | 2.02 | 0.000522 | 0.0111 up | 1.09 | 0.643 | 1 | 3 | 487 | -2.03531074 | -1.06252284 | -1.40629844 | 0.2498636 | 0.21316748 | 1.29339518 | 0.01027565 | 0.11718646 | 0.47655294 | 0.39531757 | 1.02337695 | 0.72499618 |
| ENSMUSG00000107524 | Gm44136 | predicted gene, | TEC | 15 | 21 | 17 | 36 | 35 | 47 | 28 | 35 | 38 | 37 | 44 | 38 | 1.64 | 0.0116 | | 1.76 | 0.00288 | 0.0325 up | 1.07 | 0.686 | 1 | 3 | 488 | -1.85886215 | -1.05906442 | -1.603646 | 0.38393395 | 0.3155893 | 1.26133439 | -0.32134839 | 0.33289934 | 0.57461575 | 0.46377191 | 0.99286479 | 0.51791153 |
| ENSMUSG00000115801 | Gm55330 | predicted gene, | lncRNA | 1,935 | 2,089 | 1,909 | 2,574 | 2,447 | 2,701 | 2,293 | 2,493 | 2,618 | 2,509 | 2,722 | 2,574 | 1.17 | 0.00762 | 0.115 | 1.22 | 0.000439 | 0.01 up | 1.05 | 0.403 | 1 | 3 | 489 | -1.6877069 | -1.07920044 | -1.79512559 | 0.59800628 | 0.19052138 | 0.99053327 | -0.33534936 | 0.33898278 | 0.73493043 | 0.3932422 | 1.0537547 | 0.59741126 |
| ENSMUSG00000035266 | Helq | helicase, POLQ- | protein_coding | 250 | 294 | 222 | 328 | 380 | 341 | 277 | 419 | 366 | 364 | 382 | 406 | 1.29 | 0.00477 | 0.0941 | 1.38 | 0.000292 | 0.00763 up | 1.07 | 0.443 | 1 | 3 | 490 | -1.38161759 | -0.60445534 | -1.93064687 | -0.05505604 | 0.70943247 | 0.14434229 | -0.89700503 | 1.22753264 | 0.50828319 | 0.4815356 | 0.73489885 | 1.06275581 |
| ENSMUSG00000108617 | Gm31749 | predicted gene, | lncRNA | 7 | 17 | 6 | 17 | 24 | 18 | 14 | 31 | 22 | 20 | 31 | 29 | 1.88 | 0.0214 | | 2.16 | 0.0035 | 0.0367 up | 1.15 | 0.542 | 1 | 3 | 491 | -1.70504677 | -0.20624264 | -1.82504092 | -0.1547107 | 0.54663367 | -0.0373024 | -0.55150083 | 1.17996803 | 0.36898995 | 0.1405665 | 1.23567631 | 1.00800979 |
| ENSMUSG00000104037 | Gm37776 | predicted gene, | TEC | 29 | 32 | 20 | 59 | 53 | 52 | 42 | 70 | 56 | 58 | 63 | 61 | 1.56 | 0.0134 | 0.152 | 1.69 | 0.00257 | 0.0301 up | 1.08 | 0.616 | 1 | 3 | 492 | -1.28953713 | -1.05092416 | -2.13025238 | 0.59761997 | 0.2723674 | 0.24896155 | -0.36400357 | 1.16891869 | 0.46057311 | 0.5413828 | 0.81672448 | 0.72816925 |
| ENSMUSG00000112549 | Gm35818 | predicted gene, | lncRNA | 54 | 56 | 51 | 97 | 98 | 89 | 77 | 114 | 84 | 109 | 104 | 104 | 1.46 | 0.0177 | 0.173 | 1.55 | 0.00514 | 0.0463 up | 1.06 | 0.695 | 1 | 3 | 493 | -1.47577459 | -1.3769915 | -1.6633272 | 0.5143543 | 0.54253134 | 0.17762752 | -0.35394835 | 1.13128874 | -0.02784332 | 0.97489572 | 0.78774429 | 0.76944304 |
| ENSMUSG00000043987 | Cep164 | centrosomal pr | protein_coding | 536 | 670 | 512 | 754 | 780 | 813 | 616 | 919 | 966 | 814 | 1,017 | 921 | 1.27 | 0.0155 | 0.162 | 1.5 | 0.0000255 | 0.00165 up | 1.19 | 0.0742 | 1 | 3 | 494 | -1.52375549 | -0.57315552 | -1.71022592 | -0.05094248 | 0.09973036 | 0.29064981 | -0.93538595 | 0.85036297 | 1.07941178 | 0.29493972 | 1.31824635 | 0.86012437 |
| ENSMUSG00000106841 | Gm42656 | predicted gene | TEC | 24 | 28 | 24 | 41 | 48 | 51 | 32 | 61 | 53 | 57 | 62 | 61 | 1.65 | 0.00467 | 0.0933 | 1.97 | 0.0000614 | 0.0028 up | 1.2 | 0.238 | 1 | 3 | 495 | -1.51042136 | -1.14716034 | -1.49773301 | -0.16651333 | 0.24848863 | 0.45018777 | -0.83506683 | 1.03998844 | 0.54946439 | 0.78786685 | 1.05883757 | 1.02206121 |
| ENSMUSG00000019971 | Cep290 | centrosomal pr | protein_coding | 1,012 | 1,000 | 968 | 1,316 | 1,273 | 1,230 | 1,084 | 1,352 | 1,348 | 1,317 | 1,502 | 1,345 | 1.15 | 0.0137 | 0.153 | 1.28 | 0.00000912 | 0.000926 up | 1.11 | 0.0506 | 1 | 3 | 496 | -1.27972869 | -1.35885239 | -1.58537468 | 0.53691564 | 0.30185112 | 0.06368331 | -0.81231543 | 0.72790211 | 0.70444181 | 0.53963797 | 1.47145389 | 0.69038534 |
| ENSMUSG00000028245 | Nsmaf | neutral sphingo | protein_coding | 1,083 | 1,242 | 1,068 | 1,603 | 1,479 | 1,609 | 1,274 | 1,626 | 1,666 | 1,747 | 1,775 | 1,648 | 1.2 | 0.0176 | 0.172 | 1.37 | 0.0000355 | 0.00206 up | 1.14 | 0.0788 | 1 | 3 | 497 | -1.64858935 | -0.91150971 | -1.72182236 | 0.49684403 | 0.04735073 | 0.51635705 | -0.77227799 | 0.57669512 | 0.71413285 | 0.97987934 | 1.07064576 | 0.65229452 |
| ENSMUSG00000029168 | Dpysl5 | dihydropyrimid | protein_coding | 2,027 | 2,085 | 1,890 | 2,361 | 2,337 | 2,385 | 2,195 | 2,509 | 2,493 | 2,509 | 2,713 | 2,560 | 1.13 | 0.0184 | 0.177 | 1.23 | 0.0000478 | 0.00241 up | 1.09 | 0.0899 | 1 | 3 | 498 | -1.27804241 | -1.02133126 | -1.92476688 | 0.13308653 | 0.03810477 | 0.22750981 | -0.5450995 | 0.70196789 | 0.64045322 | 0.70207496 | 1.43653389 | 0.88950898 |
| ENSMUSG00000107021 | Gm42853 | predicted gene | TEC | 29 | 35 | 21 | 60 | 47 | 56 | 46 | 71 | 75 | 81 | 84 | 67 | 1.51 | 0.0186 | 0.177 | 2.09 | 0.0000121 | 0.00106 up | 1.38 | 0.0383 | 1 | 3 | 499 | -1.32688026 | -0.96315251 | -1.89806402 | 0.27836218 | -0.35048764 | 0.11551627 | -0.36982697 | 0.72620078 | 0.885422 | 1.0986186 | 1.2281821 | 0.57610947 |
| ENSMUSG00000042156 | Dzip1 | DAZ interacting | protein_coding | 3,587 | 4,012 | 3,819 | 4,544 | 4,259 | 4,268 | 3,925 | 4,520 | 4,375 | 4,338 | 4,462 | 4,619 | 1.06 | 0.0927 | 0.358 | 1.12 | 0.00298 | 0.0333 up | 1.05 | 0.2 | 1 | 3 | 500 | -2.05233821 | -0.63149218 | -1.25797277 | 0.95478383 | 0.12923592 | 0.15423 | -0.90979541 | 0.8891412 | 0.47217979 | 0.3628384 | 0.72424792 | 1.16494152 |
| ENSMUSG00000085527 | Gm15535 | predicted gene | lncRNA | 25 | 52 | 56 | 88 | 76 | 76 | 55 | 105 | 104 | 93 | 120 | 96 | 1.42 | 0.0747 | 0.325 | 1.86 | 0.00116 | 0.0184 up | 1.31 | 0.139 | 1 | 3 | 501 | -2.22804499 | -0.89470208 | -0.74144544 | 0.38920677 | 0.01272128 | 0.00844309 | -0.7792709 | 0.90231348 | 0.85727974 | 0.54080582 | 1.29603097 | 0.63666224 |
| ENSMUSG00000113427 | Gm46378 | predicted gene, | lncRNA | 96 | 172 | 128 | 217 | 187 | 220 | 178 | 246 | 256 | 193 | 271 | 261 | 1.36 | 0.0119 | 0.144 | 1.6 | 0.0000929 | 0.00365 up | 1.17 | 0.167 | 1 | 3 | 502 | -2.16695386 | -0.46316305 | -1.37151336 | 0.34072189 | -0.17163605 | 0.38332852 | -0.33970187 | 0.78400402 | 0.93537701 | -0.07334497 | 1.14183485 | 1.00104688 |
| ENSMUSG00000074507 | Gm14340 | predicted gene | lncRNA | 60 | 90 | 63 | 137 | 102 | 131 | 101 | 135 | 128 | 128 | 163 | 136 | 1.34 | 0.0673 | 0.31 | 1.58 | 0.00351 | 0.0368 up | 1.18 | 0.278 | 1 | 3 | 503 | -1.84865715 | -0.69698238 | -1.70596015 | 0.71675321 | -0.30807507 | 0.5459961 | -0.32736756 | 0.65568679 | 0.47760928 | 0.4598031 | 1.36209854 | 0.6690953 |
| ENSMUSG00000061601 | Pclo | piccolo (presyn | protein_coding | 10,597 | 11,761 | 10,711 | 14,018 | 13,153 | 13,705 | 11,501 | 13,550 | 13,485 | 13,258 | 15,219 | 14,272 | 1.1 | 0.0937 | 0.361 | 1.19 | 0.00226 | 0.0279 up | 1.08 | 0.168 | 1 | 3 | 504 | -1.66321777 | -0.76791786 | -1.57169948 | 0.74343834 | 0.19480958 | 0.54899622 | -0.96023895 | 0.45094003 | 0.40962376 | 0.26343429 | 1.45301683 | 0.89881502 |
| ENSMUSG00000097583 | 6430590A07Rik | RIKEN cDNA 64 | lncRNA | 102 | 130 | 94 | 177 | 157 | 169 | 132 | 168 | 169 | 157 | 199 | 176 | 1.24 | 0.0708 | 0.317 | 1.39 | 0.00558 | 0.0489 up | 1.11 | 0.342 | 1 | 3 | 505 | -1.63807248 | -0.62846996 | -1.95264107 | 0.76116349 | 0.19051444 | 0.52976199 | -0.57410649 | 0.52138971 | 0.53670934 | 0.20044256 | 1.32237884 | 0.73092963 |
| ENSMUSG00000042508 | Dmtf1 | cyclin D binding | protein_coding | 1,416 | 1,627 | 1,493 | 2,004 | 1,726 | 1,865 | 1,723 | 1,789 | 1,976 | 1,831 | 2,095 | 1,996 | 1.09 | 0.173 | 0.48 | 1.21 | 0.00176 | 0.0236 up | 1.11 | 0.0774 | 1 | 3 | 506 | -1.89447292 | -0.76396955 | -1.46733869 | 0.96772711 | -0.27723493 | 0.36545241 | -0.29188216 | 0.02075844 | 0.84923673 | 0.21527701 | 1.34200238 | 0.93444416 |
| ENSMUSG00000111092 | Gm17875 | predicted gene, | processed_pseudoge | 74 | 150 | 63 | 213 | 145 | 186 | 124 | 156 | 215 | 168 | 232 | 217 | 1.22 | 0.274 | 0.593 | 1.65 | 0.00413 | 0.0404 up | 1.36 | 0.0748 | 1 | 3 | 507 | -1.67246997 | -0.08639249 | -1.97745854 | 0.87113455 | -0.18930045 | 0.48142037 | -0.56155885 | -0.00021463 | 0.89751565 | 0.19872631 | 1.11991396 | 0.91868409 |
| ENSMUSG00000063804 | Lin28b | lin-28 homolog | protein_coding | 281 | 363 | 253 | 483 | 503 | 452 | 443 | 471 | 515 | 463 | 510 | 546 | 1.35 | 0.00331 | 0.0809 | 1.47 | 0.000148 | 0.00498 up | 1.09 | 0.4 | 1 | 3 | 508 | -1.71708226 | -0.72882239 | -2.11810917 | 0.4780179 | 0.6535542 | 0.18479907 | 0.10542044 | 0.369215 | 0.75413969 | 0.2886971 | 0.71012662 | 1.0200438 |
| ENSMUSG00000107535 | Gm44183 | predicted gene, | TEC | 206 | 284 | 221 | 415 | 404 | 368 | 350 | 382 | 391 | 361 | 420 | 455 | 1.33 | 0.00985 | 0.13 | 1.44 | 0.000879 | 0.0154 up | 1.08 | 0.466 | 1 | 3 | 509 | -1.99645067 | -0.81213718 | -1.74601831 | 0.73514255 | 0.61799377 | 0.22981351 | 0.01890627 | 0.38420515 | 0.48806529 | 0.15307903 | 0.78506042 | 1.14234016 |
| ENSMUSG00000059851 | Kmt5c | lysine methyltr | protein_coding | 322 | 374 | 295 | 507 | 508 | 473 | 414 | 447 | 492 | 438 | 564 | 511 | 1.23 | 0.0329 | 0.225 | 1.34 | 0.00236 | 0.0286 up | 1.09 | 0.371 | 1 | 3 | 510 | -1.54884032 | -0.82548317 | -1.9559484 | 0.7417083 | 0.76138451 | 0.37216361 | -0.30634652 | 0.07786693 | 0.58665332 | -0.01850607 | 1.32584317 | 0.78950466 |
| ENSMUSG00000038056 | Kmt2c | lysine (K)-specif | protein_coding | 5,321 | 5,807 | 5,215 | 6,503 | 6,400 | 6,433 | 6,315 | 6,247 | 6,683 | 6,285 | 6,849 | 6,803 | 1.11 | 0.0195 | 0.18 | 1.16 | 0.000699 | 0.0133 up | 1.05 | 0.294 | 1 | 3 | 511 | -1.75010893 | -0.76799672 | -1.97457082 | 0.50767191 | 0.32584664 | 0.38409092 | 0.17629182 | 0.05467102 | 0.81452159 | 0.12161297 | 1.09184298 | 1.01612661 |
| ENSMUSG00000111696 | Gm47932 | predicted gene, | processed_pseudoge | 57 | 86 | 57 | 136 | 156 | 135 | 129 | 116 | 151 | 127 | 181 | 150 | 1.59 | 0.00164 | 0.0618 | 1.8 | 0.0000441 | 0.00231 up | 1.13 | 0.36 | 1 | 3 | 512 | -1.81862571 | -0.88686622 | -1.81876213 | 0.37992023 | 0.7998936 | 0.35499606 | 0.2313383 | -0.07386754 | 0.69172859 | 0.17858131 | 1.29273426 | 0.66892926 |
| ENSMUSG00000107041 | Gm42735 | predicted gene | TEC | 20 | 39 | 21 | 65 | 55 | 46 | 59 | 51 | 70 | 66 | 79 | 68 | 1.44 | 0.0858 | 0.347 | 1.93 | 0.00157 | 0.0221 up | 1.33 | 0.145 | 1 | 3 | 513 | -1.90241535 | -0.61609225 | -1.81328358 | 0.63851494 | 0.1942339 | -0.26849507 | 0.35971881 | 0.00046696 | 0.82254657 | 0.65649102 | 1.18728521 | 0.74102885 |
| ENSMUSG00000084336 | Or4x6 | olfactory recep | protein_coding | 17 | 49 | 18 | 57 | 55 | 48 | 53 | 54 | 67 | 65 | 73 | 69 | 1.48 | 0.0224 | 0.191 | 1.92 | 0.0000607 | 0.00278 up | 1.3 | 0.0786 | 1 | 3 | 514 | -2.03935664 | -0.04471204 | -1.97484436 | 0.33639253 | 0.24432009 | -0.08099643 | 0.11494182 | 0.18444591 | 0.75974224 | 0.66878064 | 0.99010812 | 0.84117811 |
| [illegible] | | | | | | | | | | | | | | | | | | | | | | | | | | | | | | | | | | | | | | |

| | | | | | | | |
|---|---|---|---|---|---|---|---|
| ENSMUSG00000112149 | Gm47611 | predicted gene, | processed_pseudoge | 287 | 350 | 243 | 514 |
| ENSMUSG00000104871 | Gm42639 | predicted gene | TEC | 5 | 15 | 0 | 28 |
| ENSMUSG00000116283 | Gm49412 | predicted gene, | TEC | 48 | 55 | 33 | 75 |
| ENSMUSG00000110390 | Gm45869 | predicted gene | lncRNA | 17 | 32 | 9 | 35 |
| ENSMUSG00000113978 | Gm48260 | predicted gene, | processed_pseudoge | 64 | 103 | 42 | 129 |
| ENSMUSG00000112117 | Rmst | rhabdomyosarc | lncRNA | 347 | 441 | 273 | 393 |
| ENSMUSG00000036916 | Zfp280c | zinc finger prot | protein_coding | 559 | 701 | 505 | 695 |
| ENSMUSG00000078485 | Plekhn1 | pleckstrin hom | protein_coding | 1,053 | 1,201 | 989 | 1,176 |
| ENSMUSG00000054412 | Gm4793 | predicted gene | lncRNA | 27 | 45 | 23 | 63 |
| ENSMUSG00000106714 | Gm42546 | predicted gene | TEC | 166 | 181 | 129 | 243 |
| ENSMUSG00000105270 | Gm42863 | predicted gene | TEC | 158 | 208 | 132 | 272 |
| ENSMUSG00000112689 | Gm47476 | predicted gene, | processed_pseudoge | 194 | 263 | 140 | 376 |
| ENSMUSG00000067949 | Vmn1r-ps144 | vomeronasal 1 | transcribed_unproce | 167 | 216 | 151 | 252 |
| ENSMUSG00000107182 | Gm43268 | predicted gene | TEC | 223 | 294 | 174 | 345 |
| ENSMUSG00000107996 | Gm44228 | predicted gene, | TEC | 8 | 18 | 0 | 35 |
| ENSMUSG00000090211 | Gm16050 | predicted gene | lncRNA | 7 | 10 | 2 | 13 |
| ENSMUSG00000111358 | 2900078I11Rik | RIKEN cDNA 29 | TEC | 5 | 7 | 0 | 16 |
| ENSMUSG00000041415 | Dicer1 | dicer 1, ribonuc | protein_coding | 2,607 | 2,709 | 2,339 | 2,785 |
| ENSMUSG00000038816 | Ctnnal1 | catenin (cadher | protein_coding | 469 | 540 | 411 | 623 |
| ENSMUSG00000112409 | C030006N10Rik | RIKEN cDNA C0 | TEC | 1 | 4 | 0 | 7 |
| ENSMUSG00000038733 | Wdr26 | WD repeat dom | protein_coding | 4,257 | 3,967 | 3,716 | 4,132 |
| ENSMUSG00000086796 | 4932702P03Rik | RIKEN cDNA 49 | lncRNA | 60 | 58 | 43 | 56 |
| ENSMUSG00000110717 | 1700047G07Rik | RIKEN cDNA 17 | lncRNA | 12 | 7 | 5 | 9 |
| ENSMUSG00000089798 | 1700028K03Rik | RIKEN cDNA 17 | protein_coding | 106 | 116 | 35 | 73 |
| ENSMUSG00000026447 | Pik3c2b | phosphatidylin | protein_coding | 939 | 909 | 791 | 890 |
| ENSMUSG00000042423 | Fbrs | fibrosin | protein_coding | 1,600 | 1,728 | 1,521 | 1,729 |
| ENSMUSG00000086602 | Gm15609 | predicted gene | lncRNA | 60 | 75 | 56 | 72 |
| ENSMUSG00000029420 | Rimbp2 | RIMS binding p | protein_coding | 2,775 | 2,928 | 2,429 | 3,029 |
| ENSMUSG00000115690 | Gm48924 | predicted gene, | lncRNA | 44 | 41 | 23 | 40 |
| ENSMUSG00000020516 | Rps6kb1 | ribosomal prot | protein_coding | 1,226 | 1,123 | 1,028 | 1,398 |
| ENSMUSG00000022429 | Dmc1 | DNA meiotic re | protein_coding | 28 | 19 | 13 | 42 |
| ENSMUSG00000104432 | A430027C01Rik | RIKEN cDNA A4 | TEC | 24 | 21 | 15 | 39 |
| ENSMUSG00000115955 | 9530056E24Rik | RIKEN cDNA 95 | TEC | 5 | 3 | 0 | 9 |
| ENSMUSG00000042672 | Dcst1 | DC-STAMP dom | protein_coding | 88 | 102 | 75 | 101 |
| ENSMUSG00000104433 | Gm37426 | predicted gene, | TEC | 7 | 6 | 5 | 12 |
| ENSMUSG00000026213 | Stk11ip | serine/threonir | protein_coding | 465 | 467 | 459 | 553 |
| ENSMUSG00000106855 | 5330437M03Rik | RIKEN cDNA 53 | TEC | 5 | 11 | 3 | 20 |
| ENSMUSG00000113941 | Gm35755 | predicted gene, | processed_pseudoge | 8 | 14 | 8 | 32 |
| ENSMUSG00000021725 | Parp8 | poly (ADP-ribos | protein_coding | 1,003 | 994 | 975 | 1,080 |
| ENSMUSG00000110110 | Gm30978 | predicted gene, | lncRNA | 24 | 24 | 15 | 32 |
| ENSMUSG00000112972 | Gm48417 | predicted gene, | processed_pseudoge | 85 | 67 | 51 | 151 |
| ENSMUSG00000024949 | Sf1 | splicing factor 1 | protein_coding | 4,524 | 4,376 | 4,216 | 4,741 |
| ENSMUSG00000116590 | Gm53028 | predicted gene, | lncRNA | 19 | 20 | 15 | 40 |
| ENSMUSG00000085328 | Gm17131 | predicted gene | unprocessed_pseudo | 1,250 | 1,203 | 1,211 | 1,643 |
| ENSMUSG00000099608 | 4933411E06Rik | RIKEN cDNA 49 | lncRNA | 35 | 27 | 35 | 61 |
| ENSMUSG00000103821 | D430013B06Rik | RIKEN cDNA D4 | TEC | 27 | 22 | 17 | 52 |
| ENSMUSG00000028399 | Ptprd | protein tyrosine | protein_coding | 5,987 | 6,022 | 5,727 | 7,069 |
| ENSMUSG00000112013 | Gm47967 | predicted gene, | processed_pseudoge | 56 | 44 | 47 | 109 |
| ENSMUSG00000085246 | Gm15893 | predicted gene | lncRNA | 70 | 60 | 59 | 109 |
| ENSMUSG00000112000 | Gm47956 | predicted gene, | lncRNA | 84 | 67 | 59 | 113 |
| ENSMUSG00000042104 | Uggt2 | UDP-glucose gl | protein_coding | 1,041 | 1,041 | 973 | 1,725 |
| ENSMUSG00000078902 | Gm14443 | predicted gene | protein_coding | 153 | 139 | 120 | 250 |
| ENSMUSG00000113395 | Gm48365 | predicted gene, | processed_pseudoge | 299 | 287 | 234 | 566 |
| ENSMUSG00000090386 | Mir99ahg | Mir99a and Mir | lncRNA | 184 | 114 | 107 | 205 |
| ENSMUSG00000114228 | Gm47605 | predicted gene, | processed_pseudoge | 8 | 1 | 0 | 24 |
| ENSMUSG00000054309 | Cpsf3 | cleavage and p | protein_coding | 1,331 | 1,373 | 1,245 | 1,475 |
| ENSMUSG00000113523 | Gm48366 | predicted gene, | processed_pseudoge | 65 | 66 | 39 | 99 |
| ENSMUSG00000060149 | BC002059 | cDNA sequence | lncRNA | 235 | 217 | 171 | 265 |
| ENSMUSG00000073907 | Or56b35 | olfactory recep | protein_coding | 3 | 4 | 0 | 8 |
| ENSMUSG00000103070 | Gm37903 | predicted gene, | TEC | 226 | 242 | 198 | 281 |
| ENSMUSG00000105207 | Gm42927 | predicted gene | processed_pseudoge | 332 | 443 | 204 | 1,004 |
| ENSMUSG00000086231 | Rapgef4os3 | Rap guanine nu | lncRNA | 32 | 30 | 14 | 59 |
| ENSMUSG00000112788 | Gm47821 | predicted gene, | TEC | 336 | 368 | 228 | 555 |
| ENSMUSG00000106446 | Gm42970 | predicted gene | TEC | 9 | 21 | 6 | 31 |
| ENSMUSG00000109082 | Gm44586 | predicted gene | TEC | 3 | 8 | 0 | 21 |
| ENSMUSG00000024943 | Smc5 | structural main | protein_coding | 1,065 | 1,182 | 900 | 1,273 |
| ENSMUSG00000109044 | Gm44680 | predicted gene | TEC | 1 | 4 | 0 | 11 |
| ENSMUSG00000117238 | | novel transcript | lncRNA | 212 | 267 | 146 | 334 |
| ENSMUSG00000025558 | Dock9 | dedicator of cy | protein_coding | 3,595 | 3,621 | 3,419 | 4,050 |
| ENSMUSG00000061665 | Cd2ap | CD2-associated | protein_coding | 1,378 | 1,333 | 1,338 | 1,560 |
| ENSMUSG00000020280 | Pus10 | pseudouridylat | protein_coding | 228 | 275 | 199 | 324 |
| ENSMUSG00000027692 | Tnik | TRAF2 and NCK | protein_coding | 5,193 | 5,643 | 5,359 | 6,630 |
| ENSMUSG00000031502 | Col4a1 | collagen, type I | protein_coding | 1,002 | 1,123 | 991 | 1,529 |
| ENSMUSG00000117183 | Gm20008 | predicted gene, | processed_pseudoge | 236 | 264 | 213 | 331 |
| ENSMUSG00000112596 | Gm48804 | predicted gene, | TEC | 32 | 48 | 33 | 67 |
| ENSMUSG00000034269 | Setd5 | SET domain cor | protein_coding | 3,888 | 4,119 | 3,733 | 4,247 |
| ENSMUSG00000057378 | Ryr3 | ryanodine rece | protein_coding | 1,327 | 1,905 | 1,279 | 2,243 |
| ENSMUSG00000026141 | Col19a1 | collagen, type X | protein_coding | 183 | 255 | 191 | 366 |
| ENSMUSG00000110393 | Gm36445 | predicted gene, | lncRNA | 53 | 52 | 45 | 85 |
| ENSMUSG00000019518 | Ap4m1 | adaptor-relate | protein_coding | 448 | 444 | 441 | 409 |
| ENSMUSG00000085311 | Gm15445 | predicted gene | lncRNA | 9 | 17 | 6 | 7 |
| ENSMUSG00000029727 | Cyp3a13 | cytochrome P4 | protein_coding | 11 | 15 | 9 | 17 |
| ENSMUSG00000032593 | Amigo3 | adhesion molec | protein_coding | 41 | 48 | 33 | 39 |
| ENSMUSG00000100833 | Gm28988 | predicted gene | lncRNA | 9 | 13 | 9 | 17 |
| ENSMUSG00000112352 | Gm40617 | predicted gene, | lncRNA | 5 | 7 | 5 | 12 |
| ENSMUSG00000092274 | Neat1 | nuclear paraspe | lncRNA | 3,769 | 3,881 | 3,404 | 4,127 |
| ENSMUSG00000089706 | B230216N24Rik | RIKEN cDNA B2 | lncRNA | 133 | 168 | 108 | 192 |
| ENSMUSG00000082705 | Gm15616 | predicted gene | processed_pseudoge | 1 | 6 | 2 | 7 |
| ENSMUSG00000087644 | Gm14703 | predicted gene | lncRNA | 54 | 63 | 48 | 75 |
| ENSMUSG00000028344 | Invs | inversin | protein_coding | 415 | 467 | 391 | 483 |
| ENSMUSG00000036893 | Ehmt1 | euchromatic his | protein_coding | 836 | 919 | 813 | 925 |
| ENSMUSG00000046169 | Adamts6 | a disintegrin-lik | protein_coding | 239 | 291 | 233 | 293 |
| ENSMUSG00000086735 | Gm13977 | predicted gene | lncRNA | 27 | 51 | 24 | 55 |
| ENSMUSG00000102443 | Gm37102 | predicted gene, | processed_pseudoge | 37 | 65 | 51 | 84 |
| ENSMUSG00000104641 | Gm43290 | predicted gene | TEC | 8 | 24 | 17 | 29 |
| ENSMUSG00000067276 | Capn6 | calpain 6 | protein_coding | 48 | 71 | 62 | 80 |
| ENSMUSG00000105286 | Gm43140 | predicted gene | lncRNA | 1 | 3 | 2 | 7 |
| ENSMUSG00000018068 | Ints2 | integrator com | protein_coding | 509 | 689 | 595 | 624 |
| ENSMUSG00000115814 | Gm49181 | predicted gene, | unprocessed_pseudo | 4 | 15 | 11 | 16 |
| ENSMUSG00000082286 | Pisd-ps1 | phosphatidylse | transcribed_unproce | 88 | 115 | 97 | 131 |
| ENSMUSG00000105322 | Gm43751 | predicted gene | TEC | 3 | 11 | 8 | 12 |
| ENSMUSG00000106251 | Gm42658 | predicted gene | lncRNA | 9 | 14 | 12 | 23 |
| ENSMUSG00000085269 | Gm15777 | predicted gene | lncRNA | 16 | 31 | 24 | 36 |
| ENSMUSG00000055670 | Zzef1 | zinc finger, ZZ-t | protein_coding | 2,036 | 2,527 | 2,207 | 2,546 |
| ENSMUSG00000024186 | Rgs11 | regulator of G- | protein_coding | 307 | 429 | 334 | 512 |
| ENSMUSG00000036391 | Sec24a | Sec24 related g | protein_coding | 752 | 870 | 788 | 943 |
| ENSMUSG00000020620 | Abca8b | ATP-binding ca | protein_coding | 804 | 915 | 890 | 895 |
| ENSMUSG00000087623 | Gm12404 | predicted gene | lncRNA | 21 | 38 | 33 | 41 |
| ENSMUSG00000105592 | Gm42527 | predicted gene | TEC | 3 | 6 | 6 | 8 |
| ENSMUSG00000060503 | Or2d2 | olfactory recep | protein_coding | 24 | 53 | 38 | 47 |
| ENSMUSG00000097267 | 1700109K24Rik | RIKEN cDNA 17 | lncRNA | 21 | 38 | 36 | 45 |
| ENSMUSG00000109087 | Gm45105 | predicted gene | processed_pseudoge | 21 | 39 | 36 | 44 |
| ENSMUSG00000104211 | Gm37985 | predicted gene, | TEC | 33 | 87 | 39 | 47 |
| ENSMUSG00000109775 | Gm45297 | predicted gene | TEC | 9 | 21 | 9 | 16 |
| ENSMUSG00000115756 | Gm49519 | predicted gene, | lncRNA | 70 | 116 | 86 | 88 |
| ENSMUSG00000026458 | Ppfia4 | protein tyrosine | protein_coding | 2,868 | 3,094 | 2,786 | 2,986 |
| ENSMUSG00000091345 | Col6a5 | collagen, type V | protein_coding | 170 | 280 | 139 | 204 |
| ENSMUSG00000111758 | Gm30373 | predicted gene, | lncRNA | 21 | 92 | 38 | 51 |
| ENSMUSG00000029552 | Tes | testin LIM dom | protein_coding | 27 | 64 | 38 | 48 |
| ENSMUSG00000114623 | Gm34788 | predicted gene, | lncRNA | 9 | 36 | 19 | 21 |
| ENSMUSG00000066440 | Zfyve26 | zinc finger, FYV | protein_coding | 554 | 848 | 622 | 652 |
| ENSMUSG00000023473 | Celsr3 | cadherin, EGF L | protein_coding | 2,004 | 3,218 | 2,484 | 2,916 |
| ENSMUSG00000058979 | Hdhd5 | haloacid dehal | protein_coding | 431 | 627 | 468 | 568 |
| ENSMUSG00000022948 | Setd4 | SET domain cor | protein_coding | 108 | 185 | 119 | 162 |
| ENSMUSG00000041954 | Tnfrsf18 | tumor necrosis | protein_coding | 82 | 145 | 77 | 129 |
| ENSMUSG00000115088 | Gm49165 | predicted gene, | lncRNA | 58 | 93 | 56 | 88 |
| ENSMUSG00000032558 | Nphp3 | nephronophthi | protein_coding | 276 | 508 | 393 | 427 |
| ENSMUSG00000025377 | Tepsin | TEPSIN, adapto | protein_coding | 244 | 314 | 242 | 286 |
| ENSMUSG00000118340 | Gm50301 | predicted gene, | lncRNA | 5 | 7 | 2 | 9 |
| ENSMUSG00000024561 | Mbd1 | methyl-CpG bin | protein_coding | 1,143 | 1,239 | 1,192 | 1,171 |
| ENSMUSG00000049041 | Or1r1 | olfactory recep | protein_coding | 15 | 35 | 23 | 27 |
| ENSMUSG00000002109 | Ddb2 | damage specifi | protein_coding | 189 | 236 | 234 | 236 |
| ENSMUSG00000102413 | Gm36937 | predicted gene, | lncRNA | 4 | 13 | 9 | 15 |
| ENSMUSG00000105868 | Gm43766 | predicted gene | TEC | 81 | 98 | 71 | 72 |
| ENSMUSG00000022568 | Scrib | scribbled plana | protein_coding | 1,262 | 1,300 | 1,222 | 1,254 |
| ENSMUSG00000036026 | Tmem63b | transmembran | protein_coding | 2,777 | 2,845 | 2,641 | 2,737 |
| ENSMUSG00000027346 | Gpcpd1 | glycerophosph | protein_coding | 1,427 | 1,497 | 1,492 | 1,479 |
| ENSMUSG00000053441 | Adamts19 | a disintegrin-lik | protein_coding | 30 | 49 | 35 | 50 |
| ENSMUSG00000063652 | Slc22a21 | solute carrier f | protein_coding | 65 | 89 | 91 | 96 |
| ENSMUSG00000005682 | Pan2 | PAN2 poly(A) s | protein_coding | 388 | 518 | 425 | 592 |
| ENSMUSG00000074340 | Ovgp1 | oviductal glyco | protein_coding | 85 | 164 | 129 | 192 |
| ENSMUSG00000065968 | Ifitm7 | interferon indu | protein_coding | 7 | 27 | 14 | 36 |
| ENSMUSG00000027710 | Acad9 | acyl-Coenzyme | protein_coding | 1,068 | 1,251 | 1,198 | 1,385 |
| ENSMUSG00000105095 | 8430422M14Rik | RIKEN cDNA 84 | TEC | 27 | 48 | 38 | 60 |
| ENSMUSG00000039307 | Hexdc | hexosaminidas | protein_coding | 653 | 934 | 847 | 942 |
| ENSMUSG00000031328 | Flna | filamin, alpha | protein_coding | 1,496 | 2,071 | 1,713 | 1,966 |
| ENSMUSG00000106924 | Gm42857 | predicted gene | processed_pseudoge | 304 | 463 | 340 | 490 |
| ENSMUSG00000016493 | Cd46 | CD46 antigen, c | protein_coding | 187 | 263 | 175 | 320 |
| ENSMUSG00000032308 | Ulk3 | unc-51-like kin | protein_coding | 295 | 369 | 313 | 379 |
| ENSMUSG00000104149 | Gm37138 | predicted gene, | TEC | 39 | 73 | 56 | 81 |
| ENSMUSG00000112998 | Gm47656 | predicted gene, | TEC | 45 | 84 | 48 | 79 |
| ENSMUSG00000110080 | Gm6145 | predicted gene | lncRNA | 636 | 734 | 634 | 756 |
| ENSMUSG00000079710 | Dynlt2a2 | dynein light ch | protein_coding | 67 | 76 | 69 | 89 |
| ENSMUSG00000086753 | Gm15751 | predicted gene | lncRNA | 48 | 55 | 48 | 64 |
| ENSMUSG00000038011 | Dnah10 | dynein, axonem | protein_coding | 207 | 209 | 215 | 272 |
| ENSMUSG00000115261 | Gm35769 | predicted gene, | lncRNA | 13 | 15 | 17 | 23 |
| ENSMUSG00000026726 | Cubn | cubilin (intrinsi | protein_coding | 61 | 70 | 72 | 68 |
| ENSMUSG00000047642 | D930020B18Rik | RIKEN cDNA D9 | protein_coding | 41 | 52 | 40 | 35 |
| ENSMUSG00000096351 | Samd11 | sterile alpha m | protein_coding | 117 | 164 | 121 | 149 |
| ENSMUSG00000113459 | Gm30655 | predicted gene, | lncRNA | 20 | 37 | 17 | 36 |
| ENSMUSG00000001062 | Vps9d1 | VPS9 domain c | protein_coding | 621 | 634 | 649 | 683 |
| ENSMUSG00000030400 | Ercc2 | excision repair | protein_coding | 514 | 637 | 512 | 605 |
| ENSMUSG00000102618 | Gm37848 | predicted gene, | TEC | 4 | 8 | 3 | 3 |
| ENSMUSG00000106204 | Gm21009 | predicted gene, | processed_pseudoge | 1 | 3 | 0 | 0 |
| ENSMUSG00000106150 | 4921527H02Rik | RIKEN cDNA 49 | TEC | 12 | 15 | 6 | 9 |
| ENSMUSG00000114536 | Gm48837 | predicted gene, | lncRNA | 25 | 41 | 24 | 24 |
| ENSMUSG00000023009 | Nckap5l | NCK-associated | protein_coding | 473 | 495 | 450 | 505 |
| ENSMUSG00000030091 | Nup210 | nucleoporin 21 | protein_coding | 695 | 893 | 693 | 797 |
| ENSMUSG00000102579 | Gm37965 | predicted gene, | TEC | 0 | 3 | 0 | 1 |
| ENSMUSG00000105990 | Gm43307 | predicted gene | TEC | 45 | 80 | 54 | 75 |
| ENSMUSG00000034621 | Gpatch8 | G patch domair | protein_coding | 2,130 | 2,354 | 2,138 | 2,226 |
| ENSMUSG00000087014 | Gm16364 | predicted gene | lncRNA | 21 | 28 | 24 | 27 |
| ENSMUSG00000086889 | Phf2os1 | PHD finger prot | lncRNA | 13 | 21 | 11 | 23 |
| ENSMUSG00000044037 | Als2cl | ALS2 C-termina | protein_coding | 335 | 426 | 327 | 411 |
| ENSMUSG00000106965 | Gm18290 | predicted gene, | processed_pseudoge | 11 | 18 | 11 | 16 |
| ENSMUSG00000108701 | Gm44616 | predicted gene | TEC | 15 | 27 | 12 | 19 |
| ENSMUSG00000045275 | Lca5l | Leber congenit | protein_coding | 128 | 189 | 133 | 158 |
| ENSMUSG00000100937 | Nscme3l | NSE3 homolog, | protein_coding | 15 | 37 | 14 | 16 |
| ENSMUSG00000021494 | Ddx41 | DEAD box helic | protein_coding | 479 | 528 | 452 | 500 |
| ENSMUSG00000109679 | Gm45342 | predicted gene | TEC | 12 | 20 | 9 | 15 |
| ENSMUSG00000036864 | Proser3 | proline and seri | protein_coding | 87 | 149 | 117 | 113 |
| ENSMUSG00000073973 | Or52k2 | olfactory recep | protein_coding | 66 | 93 | 90 | 76 |
| ENSMUSG00000100868 | Gm28095 | predicted gene | lncRNA | 0 | 4 | 5 | 1 |
| ENSMUSG00000102504 | Gm21955 | predicted gene, | TEC | 301 | 447 | 326 | 444 |
| ENSMUSG00000051331 | Cacna1c | calcium channe | protein_coding | 2,206 | 2,562 | 2,343 | 2,531 |
| ENSMUSG00000026199 | Ankzf1 | ankyrin repeat | protein_coding | 404 | 616 | 460 | 533 |
| ENSMUSG00000113921 | Gm48054 | predicted gene, | processed_pseudoge | 57 | 139 | 74 | 152 |
| ENSMUSG00000037363 | Letm2 | leucine zipper-E | protein_coding | 502 | 576 | 508 | 658 |
| ENSMUSG00000011751 | Sptbn4 | spectrin beta, n | protein_coding | 3,787 | 4,039 | 3,678 | 4,196 |
| ENSMUSG00000022228 | Zscan26 | zinc finger and | protein_coding | 3,529 | 3,554 | 3,382 | 3,748 |
| ENSMUSG00000028197 | Col24a1 | collagen, type X | protein_coding | 49 | 66 | 71 | 80 |
| ENSMUSG00000104348 | Gm37691 | predicted gene, | TEC | 41 | 55 | 50 | 72 |
| ENSMUSG00000028693 | Nasp | nuclear autoan | protein_coding | 376 | 428 | 345 | 402 |
| ENSMUSG00000029701 | Rbm28 | RNA binding m | protein_coding | 1,102 | 1,117 | 1,051 | 1,187 |
| ENSMUSG00000033253 | Szt2 | SZT2 subunit of | protein_coding | 1,108 | 1,660 | 1,193 | 1,357 |
| ENSMUSG00000034579 | Pla2g3 | phospholipase | protein_coding | 41 | 71 | 35 | 56 |
| ENSMUSG00000106087 | Gm43609 | predicted gene | TEC | 12 | 22 | 9 | 21 |
| ENSMUSG00000108227 | 4933412L11Rik | RIKEN cDNA 49 | TEC | 1 | 15 | 3 | 5 |
| ENSMUSG00000027284 | Cdan1 | congenital dyse | protein_coding | 448 | 659 | 497 | 660 |
| ENSMUSG00000039233 | Tbce | tubulin-specific | protein_coding | 562 | 927 | 654 | 857 |
| ENSMUSG00000040938 | Slc16a11 | solute carrier f | protein_coding | 375 | 500 | 394 | 447 |
| ENSMUSG00000104010 | Gm37366 | predicted gene, | TEC | 15 | 28 | 15 | 24 |
| ENSMUSG00000024764 | Naa40 | N(alpha)-acetyl | protein_coding | 533 | 716 | 551 | 680 |
| ENSMUSG00000114134 | Gm47408 | predicted gene, | lncRNA | 180 | 318 | 175 | 269 |
| ENSMUSG00000084846 | A730011C13Rik | RIKEN cDNA A7 | lncRNA | 34 | 42 | 27 | 44 |
| ENSMUSG00000094365 | Gm21982 | predicted gene | lncRNA | 0 | 7 | 0 | 8 |
| ENSMUSG00000109243 | Gm45867 | predicted gene | TEC | 25 | 39 | 21 | 35 |
| ENSMUSG00000103046 | Gm37309 | predicted gene, | TEC | 31 | 42 | 33 | 39 |
| ENSMUSG00000107495 | Gm44215 | predicted gene, | TEC | 497 | 689 | 557 | 614 |
| ENSMUSG00000053137 | Mapk11 | mitogen-activa | protein_coding | 472 | 575 | 486 | 538 |
| ENSMUSG00000106767 | Gm42727 | predicted gene | TEC | 9 | 32 | 21 | 28 |
| ENSMUSG00000000600 | Krit1 | KRIT1, ankyrin | protein_coding | 2,048 | 2,348 | 1,887 | 2,433 |
| ENSMUSG00000072572 | Slc39a2 | solute carrier f | protein_coding | 79 | 149 | 72 | 164 |
| ENSMUSG00000113890 | Gm48368 | predicted gene, | processed_pseudoge | 892 | 1,185 | 816 | 1,238 |
| ENSMUSG00000001415 | Smg5 | SMG5 nonsens | protein_coding | 2,066 | 2,209 | 1,933 | 2,255 |
| ENSMUSG00000044712 | Slc38a6 | solute carrier f | protein_coding | 3,725 | 4,667 | 3,451 | 4,890 |
| ENSMUSG00000102307 | Gm38194 | predicted gene, | TEC | 935 | 1,274 | 864 | 1,567 |
| ENSMUSG00000107962 | Gm43980 | predicted gene, | TEC | 404 | 589 | 350 | 767 |
| ENSMUSG00000023952 | Gtpbp2 | GTP binding pr | protein_coding | 647 | 856 | 606 | 963 |
| ENSMUSG00000027709 | Mccc1 | methylcrotono | protein_coding | 712 | 935 | 699 | 1,136 |
| ENSMUSG00000038658 | Ric1 | RAB6A GEF com | protein_coding | 1,335 | 1,511 | 1,341 | 1,656 |
| ENSMUSG00000022842 | Ece2 | endothelin con | protein_coding | 2,224 | 2,704 | 2,067 | 3,039 |
| ENSMUSG00000030846 | Tial1 | Tia1 cytotoxic g | protein_coding | 3,390 | 4,005 | 2,875 | 4,457 |
| ENSMUSG00000103839 | Gm37607 | predicted gene, | TEC | 259 | 287 | 209 | 329 |
| ENSMUSG00000020866 | Cacna1g | calcium channe | protein_coding | 2,506 | 2,823 | 2,269 | 3,349 |
| ENSMUSG00000062372 | Otof | otoferlin | protein_coding | 858 | 973 | 800 | 1,067 |
| ENSMUSG00000112038 | Gm47056 | predicted gene, | processed_pseudoge | 159 | 239 | 156 | 283 |
| ENSMUSG00000032621 | Srek1 | splicing regulat | protein_coding | 1,923 | 2,441 | 1,776 | 2,759 |
| ENSMUSG00000102700 | Gm38312 | predicted gene, | TEC | 289 | 366 | 263 | 400 |
| ENSMUSG00000108064 | Gm44423 | predicted gene, | TEC | 25 | 41 | 15 | 51 |
| ENSMUSG00000024045 | Akap8 | A kinase (PRKA) | protein_coding | 2,082 | 2,617 | 1,807 | 3,061 |
| ENSMUSG00000028248 | Pnisr | PNN interacting | protein_coding | 4,479 | 5,738 | 4,020 | 6,748 |
| ENSMUSG00000104245 | Gm38155 | predicted gene, | lncRNA | 28 | 59 | 26 | 55 |
| ENSMUSG00000111619 | Gm48348 | predicted gene, | processed_pseudoge | 177 | 247 | 173 | 261 |
| ENSMUSG00000021495 | Fam193b | family with seq | protein_coding | 1,633 | 2,047 | 1,668 | 2,151 |
| ENSMUSG00000107632 | Gm44070 | predicted gene, | TEC | 90 | 131 | 83 | 160 |
| ENSMUSG00000014355 | Anapc1 | anaphase prom | protein_coding | 2,513 | 3,190 | 2,524 | 3,437 |
| ENSMUSG00000025574 | Tk1 | thymidine kina | protein_coding | 65 | 115 | 65 | 135 |
| ENSMUSG00000034206 | Polq | polymerase (DN | protein_coding | 37 | 74 | 40 | 87 |
| ENSMUSG00000085913 | Gm15601 | predicted gene | lncRNA | 46 | 66 | 37 | 86 |

| | | | | | | | | | | | | | | | | | | | | | | | | | | | | | | | | | | | | | | | |
|---|---|---|---|---|---|---|---|---|---|---|---|---|---|---|---|---|---|---|---|---|---|---|---|---|---|---|---|---|---|---|---|---|---|---|---|---|---|---|---|
| ENSMUSG00000114148 | Gm47701 | predicted gene, | lncRNA | 56 | 77 | 42 | 83 | 103 | 118 | 103 | 99 | 116 | 104 | 141 | 155 | 1.63 | 0.00391 | 0.0863 | 1.99 | 0.0000325 | 0.00194 | up | 1.22 | 0.205 | 1 | 3 | 957 | -1.4110796 | -0.6108096 | -2.01433931 | -0.4320132 | 0.1920339 | 0.59973795 | 0.17505824 | 0.06509442 | 0.54237399 | 0.22361751 | 1.17605835 | 1.49426734 |
| ENSMUSG00000106864 | Gtf3c2 | general transcri | protein_coding | 964 | 1,291 | 930 | 1,275 | 1,356 | 1,339 | 1,322 | 1,338 | 1,488 | 1,403 | 1,638 | 1,641 | 1.2 | 0.011 | 0.138 | 1.38 | 0.00000617 | 0.000738 | up | 1.15 | 0.0485 | 1 | 3 | 958 | -1.76528481 | -0.1118578 | -1.96394321 | -0.18616118 | 0.17038788 | 0.09796323 | 0.02316779 | 0.09103492 | 0.71042578 | 0.36852796 | 1.27795006 | 1.2877894 |
| ENSMUSG00000110077 | Gm45632 | predicted gene | TEC | 24 | 45 | 15 | 45 | 60 | 59 | 47 | 67 | 72 | 67 | 74 | 87 | 1.81 | 0.00364 | 0.0838 | 2.31 | 0.0000214 | 0.00147 | up | 1.28 | 0.174 | 1 | 3 | 959 | -1.51115259 | -0.34973227 | -2.16760689 | -0.33329426 | 0.31469291 | 0.25832328 | -0.23763545 | 0.57554977 | 0.76972639 | 0.58242176 | 0.82909278 | 1.26961456 |
| ENSMUSG00000109031 | Gm45201 | predicted gene | TEC | 24 | 28 | 11 | 27 | 29 | 46 | 37 | 39 | 46 | 41 | 52 | 49 | 1.68 | 0.0146 | | 2.08 | 0.000334 | 0.00831 | up | 1.24 | 0.246 | 1 | 3 | 960 | -0.86329102 | -0.49636032 | -2.33762935 | -0.61847436 | -0.43017809 | 0.79139013 | 0.20222045 | 0.34128596 | 0.79829594 | 0.4468476 | 1.15407046 | 1.0118226 |
| ENSMUSG00000004565 | Pnpla6 | patatin-like phc | protein_coding | 1,644 | 1,790 | 1,577 | 1,834 | 1,834 | 2,110 | 1,936 | 1,960 | 1,994 | 1,979 | 2,225 | 2,166 | 1.15 | 0.0201 | 0.183 | 1.22 | 0.000545 | 0.0115 | up | 1.07 | 0.262 | 1 | 3 | 961 | -1.43962992 | -0.6340111 | -1.83396111 | -0.39972666 | -0.40014217 | 0.95050707 | 0.11676692 | 0.23710036 | 0.40197171 | 0.3313434 | 1.46457497 | 1.20520652 |
| ENSMUSG00000107796 | Gm44068 | predicted gene, | TEC | 3 | 6 | 2 | 7 | 9 | 17 | 9 | 16 | 12 | 11 | 18 | 19 | 3.07 | 0.00263 | | 3.62 | 0.000365 | 0.00879 | up | 1.18 | 0.538 | 1 | 3 | 962 | -1.44828748 | -0.71896722 | -1.84612963 | -0.51218158 | -0.1359265 | 0.99362398 | -0.10825052 | 0.85078049 | 0.33756457 | 0.21844978 | 1.13369045 | 1.23563366 |
| ENSMUSG00000049676 | Catsperg1 | cation channel : | protein_coding | 67 | 141 | 43 | 115 | 133 | 200 | 93 | 140 | 160 | 148 | 177 | 165 | 1.54 | 0.0189 | 0.178 | 1.78 | 0.0016 | 0.0223 | up | 1.15 | 0.428 | 1 | 3 | 963 | -1.42438729 | 0.27846967 | -2.18239512 | -0.2285104 | 0.13011653 | 1.26233119 | -0.73055932 | 0.25891138 | 0.61869141 | 0.40410364 | 0.90665428 | 0.70657403 |
| ENSMUSG00000112319 | Gm47221 | predicted gene, | lncRNA | 125 | 214 | 93 | 185 | 209 | 325 | 151 | 209 | 266 | 259 | 279 | 304 | 1.45 | 0.0302 | 0.218 | 1.79 | 0.000521 | 0.0111 | up | 1.24 | 0.193 | 1 | 3 | 964 | -1.33447913 | 0.04599999 | -1.97455193 | -0.34539932 | -0.02053496 | 1.31025195 | -0.87323895 | -0.02132711 | 0.68049142 | 0.6102619 | 0.82865662 | 1.09386951 |
| ENSMUSG00000034656 | Cacna1a | calcium channe | protein_coding | 3,366 | 3,971 | 3,244 | 4,054 | 3,908 | 4,358 | 3,782 | 4,122 | 4,359 | 4,360 | 4,846 | 4,464 | 1.1 | 0.0744 | 0.324 | 1.23 | 0.000181 | 0.0056 | up | 1.11 | 0.0501 | 1 | 3 | 965 | -1.58216989 | -0.16183625 | -1.89808772 | 0.01605149 | -0.29978941 | 0.64196992 | -0.58193752 | 0.16069354 | 0.64482212 | 0.64708917 | 1.56353293 | 0.84966162 |
| ENSMUSG00000097536 | 2610037D02Rik | RIKEN cDNA 26 | lncRNA | 78 | 121 | 78 | 113 | 108 | 144 | 98 | 121 | 128 | 136 | 154 | 142 | 1.2 | 0.105 | 0.38 | 1.43 | 0.00114 | 0.0183 | up | 1.19 | 0.101 | 1 | 3 | 966 | -1.7134692 | 0.15950013 | -1.70369124 | -0.15830834 | -0.3441118 | 0.99077871 | -0.79629195 | 0.15462818 | 0.41865605 | 0.72468117 | 1.33556488 | 0.93206342 |
| ENSMUSG00000033526 | Ppip5k1 | diphosphoinosi | protein_coding | 3,482 | 3,811 | 3,154 | 3,697 | 3,525 | 3,862 | 3,487 | 3,572 | 3,980 | 3,883 | 4,035 | 3,942 | 1.02 | 0.595 | 0.831 | 1.12 | 0.00377 | 0.0384 | up | 1.1 | 0.0176 | 1 | 3 | 967 | -0.81426627 | 0.42871018 | -2.16794608 | 0.01033822 | -0.64546937 | 0.61280395 | -0.79264868 | -0.46235874 | 1.02943214 | 0.68638741 | 1.21773793 | 0.8972793 |
| ENSMUSG00000004896 | Rrnad1 | ribosomal RNA | protein_coding | 679 | 776 | 608 | 839 | 754 | 815 | 741 | 751 | 822 | 829 | 954 | 859 | 1.05 | 0.384 | 0.692 | 1.19 | 0.00274 | 0.0314 | up | 1.13 | 0.032 | 1 | 3 | 968 | -1.1977139 | -0.06116393 | -2.11463299 | 0.61079067 | -0.30300213 | 0.35752713 | -0.46081834 | -0.34291846 | 0.43499849 | 0.51041529 | 1.74378306 | 0.82273511 |
| ENSMUSG00000024290 | Rock1 | Rho-associated | protein_coding | 1,433 | 1,564 | 1,153 | 1,630 | 1,593 | 1,725 | 1,540 | 1,614 | 1,705 | 1,697 | 1,911 | 1,731 | 1.12 | 0.0491 | 0.27 | 1.22 | 0.00055 | 0.0115 | up | 1.09 | 0.138 | 1 | 3 | 969 | -0.87805495 | -0.17569448 | -2.58080798 | 0.15748081 | -0.02676717 | 0.61812505 | -0.29907896 | 0.0807063 | 0.52340345 | 0.48294609 | 1.45295979 | 0.64478205 |
| ENSMUSG00000027130 | Slc12a6 | solute carrier fa | protein_coding | 3,872 | 4,220 | 3,629 | 4,283 | 4,131 | 4,362 | 4,008 | 4,315 | 4,421 | 4,449 | 4,524 | 4,363 | 1.05 | 0.174 | 0.48 | 1.11 | 0.00403 | 0.0399 | up | 1.06 | 0.131 | 1 | 3 | 970 | -1.28425307 | 0.04508969 | -2.28442783 | 0.27614626 | -0.28377332 | 0.56200235 | -0.75294316 | 0.39082462 | 0.76965955 | 0.86893857 | 1.12712096 | 0.56561538 |
| ENSMUSG00000031503 | Col4a2 | collagen, type I | protein_coding | 1,417 | 2,349 | 1,532 | 2,486 | 2,554 | 2,937 | 1,864 | 2,357 | 2,739 | 2,813 | 2,786 | 3,223 | 1.25 | 0.0612 | 0.297 | 1.49 | 0.0008 | 0.0144 | up | 1.19 | 0.139 | 1 | 3 | 971 | -1.94673255 | -0.01764932 | -1.65457111 | 0.20338799 | 0.3098904 | 0.86158997 | -0.9115092 | -0.00433371 | 0.58467081 | 0.6909915 | 0.65301794 | 1.23124728 |
| ENSMUSG00000037300 | Ttc13 | tetratricopeptic | protein_coding | 764 | 949 | 794 | 1,141 | 1,128 | 1,252 | 912 | 1,062 | 1,121 | 1,135 | 1,262 | 1,291 | 1.19 | 0.0426 | 0.25 | 1.32 | 0.0014 | 0.0206 | up | 1.11 | 0.245 | 1 | 3 | 972 | -1.81084022 | -0.60673335 | -1.59979374 | 0.44856274 | 0.38353684 | 0.99555191 | -0.83113366 | 0.03534025 | 0.34855884 | 0.42155678 | 1.04159164 | 1.17380197 |
| ENSMUSG00000024287 | Thoc1 | THO complex 1 | protein_coding | 788 | 1,059 | 852 | 1,296 | 1,234 | 1,425 | 1,019 | 1,146 | 1,272 | 1,365 | 1,482 | 1,409 | 1.21 | 0.0314 | 0.222 | 1.38 | 0.000186 | 0.0057 | up | 1.15 | 0.114 | 1 | 3 | 973 | -1.92864631 | -0.52172819 | -1.56270034 | 0.47970246 | 0.23377675 | 0.95644315 | -0.70713896 | -0.13404016 | 0.38472665 | 0.74113176 | 1.15687317 | 0.90160001 |
| ENSMUSG00000028869 | Gnl2 | guanine nucleo | protein_coding | 1,286 | 1,466 | 1,315 | 1,600 | 1,534 | 1,711 | 1,421 | 1,511 | 1,586 | 1,623 | 1,649 | 1,680 | 1.09 | 0.0795 | 0.335 | 1.15 | 0.00299 | 0.0333 | up | 1.06 | 0.23 | 1 | 3 | 974 | -1.84420827 | -0.44092686 | -1.60137938 | 0.51119633 | 0.05318925 | 1.24543095 | -0.77464633 | -0.11545337 | 0.41678607 | 0.66723372 | 0.84121343 | 1.04156446 |
| ENSMUSG00000018986 | Slfn3 | schlafen 3 | protein_coding | 41 | 65 | 45 | 73 | 74 | 93 | 64 | 67 | 88 | 71 | 109 | 92 | 1.33 | 0.0391 | 0.243 | 1.6 | 0.000306 | 0.0079 | up | 1.21 | 0.121 | 1 | 3 | 975 | -1.80547742 | -0.37213928 | -1.53821891 | 0.03695552 | 0.10422646 | 0.95650082 | -0.42649539 | -0.22283588 | 0.75771426 | -0.04964456 | 1.61517175 | 0.94424263 |
| ENSMUSG00000027966 | Col11a1 | collagen, type X | protein_coding | 450 | 770 | 589 | 952 | 1,055 | 1,186 | 750 | 867 | 1,245 | 1,013 | 1,375 | 1,230 | 1.4 | 0.0425 | 0.25 | 1.76 | 0.000569 | 0.0118 | up | 1.26 | 0.156 | 1 | 3 | 976 | -2.06373585 | -0.5453635 | -1.32621583 | 0.10039418 | 0.42320831 | 0.79519854 | -0.62484468 | -0.18666149 | 0.94971384 | 0.29476366 | 1.27226318 | 0.91127964 |
| ENSMUSG00000097545 | Mir124a-1hg | Mir124-1 host g | lncRNA | 1,117 | 1,543 | 1,122 | 1,647 | 1,790 | 1,930 | 1,515 | 1,659 | 1,842 | 1,767 | 2,086 | 2,050 | 1.27 | 0.00498 | 0.0952 | 1.43 | 0.0000265 | 0.00169 | up | 1.12 | 0.165 | 1 | 3 | 977 | -1.86786167 | -0.31704844 | -1.84675776 | 0.00290921 | 0.41606465 | 0.79140221 | -0.40623832 | 0.03887482 | 0.55788535 | 0.35284529 | 1.18298547 | 1.09493919 |
| ENSMUSG00000113290 | A530058O07Rik | RIKEN cDNA A5 | TEC | 37 | 77 | 45 | 84 | 87 | 113 | 64 | 78 | 103 | 95 | 122 | 126 | 1.4 | 0.0454 | 0.258 | 1.83 | 0.000209 | 0.00613 | up | 1.31 | 0.0853 | 1 | 3 | 978 | -1.92542307 | -0.20520598 | -1.52645028 | 0.02739538 | 0.11879978 | 0.9139928 | -0.70150324 | -0.17274512 | 0.63098786 | 0.39614915 | 1.1639618 | 1.28004092 |
| ENSMUSG00000105663 | Gm43606 | predicted gene | TEC | 54 | 83 | 57 | 96 | 97 | 113 | 87 | 89 | 119 | 115 | 131 | 143 | 1.32 | 0.0495 | 0.27 | 1.74 | 0.0000532 | 0.00259 | up | 1.32 | 0.0358 | 1 | 3 | 979 | -1.77039756 | -0.5115449 | -1.63985461 | -0.01762227 | 0.01046583 | 0.55505953 | -0.34895557 | -0.26210428 | 0.76513353 | 0.61727877 | 1.12401706 | 1.47852446 |
| ENSMUSG00000094840 | Muc3a | mucin 3A, cell s | protein_coding | 12 | 30 | 18 | 40 | 45 | 56 | 28 | 41 | 54 | 60 | 70 | 74 | 1.71 | 0.0133 | 0.152 | 2.58 | 0.00000464 | 0.000636 | up | 1.51 | 0.0289 | 1 | 3 | 980 | -1.89004085 | -0.62947285 | -1.38358828 | -0.07033097 | 0.17594314 | 0.66084297 | -0.70808545 | -0.02647609 | 0.54304961 | 0.80101199 | 1.18378425 | 1.34336253 |
| ENSMUSG00000032637 | Atxn2l | ataxin 2-like | protein_coding | 3,407 | 4,118 | 3,553 | 4,458 | 4,521 | 4,800 | 4,134 | 4,526 | 4,790 | 4,786 | 5,192 | 5,156 | 1.16 | 0.00944 | 0.128 | 1.28 | 0.00000918 | 0.000926 | up | 1.11 | 0.0663 | 1 | 3 | 981 | -1.94630784 | -0.53274995 | -1.63406779 | 0.06406418 | 0.1691916 | 0.62161693 | -0.50299803 | 0.17841273 | 0.60594821 | 0.59955887 | 1.21525855 | 1.16207254 |
| [illegible] | | | | | | | | | | | | | | | | | | | | | | | | | | | | | | | | | | | | | | | |

| | | | |
|---|---|---|---|
| ENSMUSG00000110997 | 2610028D06Rik | RIKEN cDNA 26 | TEC |
| ENSMUSG00000038371 | Sbf2 | SET binding fac | protein_coding |
| ENSMUSG00000109505 | Gm44677 | predicted gene | TEC |
| ENSMUSG00000029823 | Luc7l2 | LUC7-like 2 (S. c | protein_coding |
| ENSMUSG00000041488 | Stx3 | syntaxin 3 | protein_coding |
| ENSMUSG00000102729 | Gm32444 | predicted gene, | TEC |
| ENSMUSG00000008200 | Fnbp4 | formin binding | protein_coding |
| ENSMUSG00000117434 | 1810014P07Rik | RIKEN cDNA 18 | lncRNA |
| ENSMUSG00000049571 | Cfap46 | cilia and flagell | protein_coding |
| ENSMUSG00000104237 | Gm33533 | predicted gene, | lncRNA |
| ENSMUSG00000053889 | Kirrel3os | kirre like nephr | lncRNA |
| ENSMUSG00000102602 | A930004J17Rik | RIKEN cDNA A9 | TEC |
| ENSMUSG00000109394 | A230057D06Rik | RIKEN cDNA A2 | lncRNA |
| ENSMUSG00000113476 | Gm48309 | predicted gene, | TEC |
| ENSMUSG00000051965 | Nanos2 | nanos C2HC-typ | protein_coding |
| ENSMUSG00000114253 | Gm47798 | predicted gene, | lncRNA |
| ENSMUSG00000106665 | Gm43389 | predicted gene | TEC |
| ENSMUSG00000105454 | Gm43830 | predicted gene | TEC |
| ENSMUSG00000113146 | Gm47406 | predicted gene, | processed_pseudoge |
| ENSMUSG00000062296 | Trank1 | tetratricopeptic | protein_coding |
| ENSMUSG00000097767 | Miat | myocardial infa | lncRNA |
| ENSMUSG00000102858 | Gm37086 | predicted gene, | TEC |
| ENSMUSG00000031706 | Rfx1 | regulatory facto | protein_coding |
| ENSMUSG00000109196 | Gm44715 | predicted gene | processed_pseudoge |
| ENSMUSG00000052566 | Hook2 | hook microtubu | protein_coding |
| ENSMUSG00000104156 | Gm38102 | predicted gene, | lncRNA |
| ENSMUSG00000058740 | Kcnt1 | potassium chan | protein_coding |
| ENSMUSG00000111264 | E030022I16Rik | RIKEN cDNA E0 | TEC |
| ENSMUSG00000075590 | Nrbp2 | nuclear recepto | protein_coding |
| ENSMUSG00000110227 | Gm45634 | predicted gene | TEC |
| ENSMUSG00000103867 | 9630010A21Rik | RIKEN cDNA 96 | TEC |
| ENSMUSG00000006471 | Ndor1 | NADPH depend | protein_coding |
| ENSMUSG00000033439 | Trmt13 | tRNA methyltra | protein_coding |
| ENSMUSG00000033902 | Mapkbp1 | mitogen-activat | protein_coding |
| ENSMUSG00000108802 | Gm44769 | predicted gene | TEC |
| ENSMUSG00000115220 | Gm49768 | predicted gene, | lncRNA |
| ENSMUSG00000041879 | Ipo9 | importin 9 | protein_coding |
| ENSMUSG00000029428 | Stx2 | syntaxin 2 | protein_coding |
| ENSMUSG00000029389 | Ddx55 | DEAD box helic | protein_coding |
| ENSMUSG00000115031 | Gm49051 | predicted gene, | processed_pseudoge |
| ENSMUSG00000046613 | Vwa5b2 | von Willebrand | protein_coding |
| ENSMUSG00000015377 | Dennd6b | DENN/MADD d | protein_coding |
| ENSMUSG00000102776 | Gm38162 | predicted gene, | TEC |
| ENSMUSG00000108072 | Gm20362 | predicted gene, | lncRNA |
| ENSMUSG00000038685 | Rtel1 | regulator of tel | protein_coding |
| ENSMUSG00000059326 | Csf2ra | colony stimulat | protein_coding |
| ENSMUSG00000031310 | Zmym3 | zinc finger, MYI | protein_coding |
| ENSMUSG00000031398 | Plxna3 | plexin A3 | protein_coding |
| ENSMUSG00000114268 | | | |
| ENSMUSG00000031295 | Phka2 | phosphorylase | protein_coding |
| ENSMUSG00000112496 | Gm48764 | predicted gene, | lncRNA |
| ENSMUSG00000026088 | Mitd1 | MIT, microtubu | protein_coding |
| ENSMUSG00000028188 | Spata1 | spermatogenes | protein_coding |
| ENSMUSG00000025188 | Hps1 | HPS1, biogenes | protein_coding |
| ENSMUSG00000110246 | C130073E24Rik | RIKEN cDNA C1 | lncRNA |
| ENSMUSG00000025816 | Sec61a2 | Sec61, alpha su | protein_coding |
| ENSMUSG00000112392 | Gm35240 | predicted gene, | lncRNA |
| ENSMUSG00000048108 | Tmem72 | transmembrane | protein_coding |
| ENSMUSG00000102302 | Gm38190 | predicted gene, | lncRNA |
| ENSMUSG00000115267 | Vmn1r76 | vomeronasal 1 | protein_coding |
| ENSMUSG00000115238 | Gm19236 | predicted gene, | processed_pseudoge |
| ENSMUSG00000039801 | Cplane1 | ciliogenesis anc | protein_coding |
| ENSMUSG00000085396 | Firre | functional inter | lncRNA |
| ENSMUSG00000085741 | 5430405H02Rik | RIKEN cDNA 54 | lncRNA |
| ENSMUSG00000114413 | | | |
| ENSMUSG00000029068 | Ccnl2 | cyclin L2 | protein_coding |
| ENSMUSG00000004677 | Myo9b | myosin IXb | protein_coding |
| ENSMUSG00000026784 | Pdss1 | prenyl (solanes | protein_coding |
| ENSMUSG00000015568 | Lpl | lipoprotein lipa | protein_coding |
| ENSMUSG00000112516 | Gm47968 | predicted gene, | processed_pseudoge |
| ENSMUSG00000112142 | Gm47022 | predicted gene, | TEC |
| ENSMUSG00000041354 | Rgl2 | ral guanine nuc | protein_coding |
| ENSMUSG00000086228 | Ubap1l | ubiquitin-assoc | protein_coding |
| ENSMUSG00000085180 | AI838599 | expressed sequ | lncRNA |
| ENSMUSG00000058084 | Or9k2 | olfactory recep | protein_coding |
| ENSMUSG00000073879 | Gm54215 | predicted gene, | unprocessed_pseudo |
| ENSMUSG00000029055 | Plch2 | phospholipase | protein_coding |
| ENSMUSG00000036555 | Iqce | IQ motif contai | protein_coding |
| ENSMUSG00000038535 | Zfp280d | zinc finger prot | protein_coding |
| ENSMUSG00000001052 | Sec24b | Sec24 related g | protein_coding |
| ENSMUSG00000110637 | Gm7807 | predicted gene | processed_pseudoge |
| ENSMUSG00000117440 | Gm50087 | predicted gene, | lncRNA |
| ENSMUSG00000074024 | 4632427E13Rik | RIKEN cDNA 46 | lncRNA |
| ENSMUSG00000048191 | | | |
| ENSMUSG00000110266 | Gm32742 | predicted gene, | protein_coding |
| ENSMUSG00000041215 | Yeats2 | YEATS domain | protein_coding |
| ENSMUSG00000022475 | Hdac7 | histone deacety | protein_coding |
| ENSMUSG00000097379 | Gm26873 | predicted gene, | lncRNA |
| ENSMUSG00000101872 | Gm29237 | predicted gene | lncRNA |
| ENSMUSG00000056724 | Nbeal2 | neurobeachin-l | protein_coding |
| ENSMUSG00000102960 | Gm37943 | predicted gene, | TEC |
| ENSMUSG00000027715 | Ccna2 | cyclin A2 | protein_coding |
| ENSMUSG00000067356 | B430203G13Rik | RIKEN cDNA B4 | TEC |
| ENSMUSG00000103857 | Gm37249 | predicted gene, | TEC |
| ENSMUSG00000032739 | Pram1 | PML-RAR alpha | protein_coding |
| ENSMUSG00000036292 | Gramd1c | GRAM domain | protein_coding |
| ENSMUSG00000021027 | Ralgapa1 | Ral GTPase acti | protein_coding |
| ENSMUSG00000113893 | Gm48800 | predicted gene, | processed_pseudoge |
| ENSMUSG00000114407 | Gm48765 | predicted gene, | lncRNA |
| ENSMUSG00000106870 | Gm43681 | predicted gene | TEC |
| ENSMUSG00000019734 | Tmc4 | transmembrane | protein_coding |
| ENSMUSG00000023912 | Slc25a27 | solute carrier fa | protein_coding |
| ENSMUSG00000038759 | Nup205 | nucleoporin 20 | protein_coding |
| ENSMUSG00000020212 | Mdm1 | transformed m | protein_coding |
| ENSMUSG00000042185 | Nfrkb | nuclear factor r | protein_coding |
| ENSMUSG00000075569 | Rsph10b | radial spoke he | protein_coding |
| ENSMUSG00000100572 | 2410021H03Rik | RIKEN cDNA 24 | lncRNA |
| ENSMUSG00000001750 | Tcirg1 | T cell, immune | protein_coding |
| ENSMUSG00000047050 | Or8b50 | olfactory recep | protein_coding |
| ENSMUSG00000015659 | Serac1 | serine active sit | protein_coding |
| ENSMUSG00000105742 | Gm42748 | predicted gene | TEC |
| ENSMUSG00000047594 | Or10ag57 | olfactory recep | protein_coding |
| ENSMUSG00000055629 | B4galnt4 | beta-1,4-N-acet | protein_coding |
| ENSMUSG00000046959 | Slc26a1 | solute carrier fa | protein_coding |
| ENSMUSG00000091531 | Or12d2 | olfactory recep | protein_coding |
| ENSMUSG00000103313 | Gm38357 | predicted gene, | TEC |
| ENSMUSG00000103662 | Gm34294 | predicted gene, | TEC |
| ENSMUSG00000103214 | Gm38286 | predicted gene, | TEC |
| ENSMUSG00000026349 | Ccnt2 | cyclin T2 | protein_coding |
| ENSMUSG00000092032 | Vmn2r59 | vomeronasal 2, | protein_coding |
| ENSMUSG00000046101 | Mcmdc2 | minichromoson | protein_coding |
| ENSMUSG00000097680 | Gm26642 | predicted gene, | lncRNA |
| ENSMUSG00000029439 | Sfswap | splicing factor S | protein_coding |
| ENSMUSG00000106904 | Gm43029 | predicted gene | TEC |
| ENSMUSG00000097286 | Gm26684 | predicted gene, | lncRNA |
| ENSMUSG00000078624 | Or51ab3 | olfactory recep | protein_coding |
| ENSMUSG00000095928 | Or5ac22 | olfactory recep | protein_coding |
| ENSMUSG00000108298 | Gm44226 | predicted gene, | lncRNA |
| ENSMUSG00000097617 | Gm10687 | predicted gene | lncRNA |
| ENSMUSG00000109909 | Gm45663 | predicted gene | TEC |
| ENSMUSG00000078808 | Vmn1r58 | vomeronasal 1 | protein_coding |
| ENSMUSG00000118049 | Gm50103 | predicted gene, | lncRNA |
| ENSMUSG00000026637 | Traf5 | TNF receptor-a | protein_coding |
| ENSMUSG00000106962 | Gm43633 | predicted gene | TEC |
| ENSMUSG00000102610 | Gm37846 | predicted gene, | TEC |
| ENSMUSG00000085811 | Cep112it | centrosomal pr | lncRNA |
| ENSMUSG00000103183 | Gm37090 | predicted gene, | TEC |
| ENSMUSG00000028047 | Thbs3 | thrombospondi | protein_coding |
| ENSMUSG00000113673 | Gm47486 | predicted gene, | lncRNA |
| ENSMUSG00000021013 | Ttc8 | tetratricopeptic | protein_coding |
| ENSMUSG00000109028 | Gm35842 | predicted gene, | lncRNA |
| ENSMUSG00000072889 | Nfxl1 | nuclear transcri | protein_coding |
| ENSMUSG00000112355 | Gm48285 | predicted gene, | TEC |
| ENSMUSG00000041144 | Dnah7b | dynein, axonem | protein_coding |
| ENSMUSG00000115816 | Gm34589 | predicted gene, | lncRNA |
| ENSMUSG00000039396 | Neil3 | nei like 3 (E. col | protein_coding |
| ENSMUSG00000048697 | Vmn1r26 | vomeronasal 1 | protein_coding |
| ENSMUSG00000100768 | Gm29055 | predicted gene | lncRNA |
| ENSMUSG00000040651 | Tasor | transcription ac | protein_coding |
| ENSMUSG00000084136 | Vmn1r207 | vomeronasal 1 | protein_coding |
| ENSMUSG00000102224 | 4930447F24Rik | RIKEN cDNA 49 | lncRNA |
| ENSMUSG00000108297 | Gm44167 | predicted gene, | lncRNA |
| ENSMUSG00000085998 | AW822252 | expressed sequ | transcribed_unproce |
| ENSMUSG00000109575 | Gm44831 | predicted gene | TEC |
| ENSMUSG00000113055 | Gm48700 | predicted gene, | TEC |
| ENSMUSG00000063730 | Hsd3b2 | hydroxy-delta-5 | protein_coding |
| ENSMUSG00000107094 | Gm43294 | predicted gene | TEC |
| ENSMUSG00000039531 | Zup1 | zinc finger cont | protein_coding |
| ENSMUSG00000059866 | Tnip2 | TNFAIP3 intera | protein_coding |
| ENSMUSG00000022483 | Col2a1 | collagen, type I | protein_coding |
| ENSMUSG00000097825 | 9630001P10Rik | RIKEN cDNA 96 | lncRNA |
| ENSMUSG00000112925 | Gm31763 | predicted gene, | lncRNA |
| ENSMUSG00000084968 | Gm12743 | predicted gene | lncRNA |
| ENSMUSG00000089941 | Gm16168 | predicted gene | lncRNA |
| ENSMUSG00000040287 | Stac3 | SH3 and cystein | protein_coding |
| ENSMUSG00000024940 | Ltbp3 | latent transforr | protein_coding |
| ENSMUSG00000056856 | Jakmip3 | janus kinase an | protein_coding |
| ENSMUSG00000067736 | Gm10222 | predicted gene | unprocessed_pseudo |
| ENSMUSG00000112008 | Gm47218 | predicted gene, | TEC |
| ENSMUSG00000117102 | Gm55377 | predicted gene, | TEC |
| ENSMUSG00000108402 | 9430064I24Rik | RIKEN cDNA 94 | TEC |
| ENSMUSG00000014418 | Hps5 | HPS5, biogenes | protein_coding |
| ENSMUSG00000105052 | Gm21655 | predicted gene, | lncRNA |
| ENSMUSG00000028107 | Tars2 | threonyl-tRNA | protein_coding |
| ENSMUSG00000051510 | Mafg | v-maf musculoa | protein_coding |
| ENSMUSG00000090655 | Vmn2r120 | vomeronasal 2, | protein_coding |
| ENSMUSG00000021713 | Ppwd1 | peptidylprolyl i | protein_coding |
| ENSMUSG00000053644 | Aldh7a1 | aldehyde dehyc | protein_coding |
| ENSMUSG00000037926 | Ssh2 | slingshot protei | protein_coding |
| ENSMUSG00000090093 | Gm14399 | predicted gene | protein_coding |
| ENSMUSG00000094893 | Gm4791 | predicted gene | lncRNA |
| ENSMUSG00000033031 | Cip2a | cell proliferatio | protein_coding |
| ENSMUSG00000110945 | Gm9856 | predicted gene | lncRNA |
| ENSMUSG00000075225 | Ccdc162 | coiled-coil dom | protein_coding |
| ENSMUSG00000023007 | Prpf40b | pre-mRNA proc | protein_coding |
| ENSMUSG00000040838 | Gm11639 | predicted gene | protein_coding |
| ENSMUSG00000078173 | Lenep | lens epithelial p | protein_coding |
| ENSMUSG00000021758 | Ddx4 | DEAD box helic | protein_coding |
| ENSMUSG00000030680 | Pagr1a | PAXIP1 associat | protein_coding |
| ENSMUSG00000048677 | Tpcn2 | two pore segm | protein_coding |
| ENSMUSG00000113800 | 2210039B01Rik | RIKEN cDNA 22 | lncRNA |
| ENSMUSG00000056900 | Usp13 | ubiquitin specif | protein_coding |
| ENSMUSG00000108766 | Gm44760 | predicted gene | TEC |
| ENSMUSG00000078919 | Dpm1 | dolichol-phosph | protein_coding |
| ENSMUSG00000033760 | Rbm4b | RNA binding m | protein_coding |
| ENSMUSG00000067928 | Zfp760 | zinc finger prot | protein_coding |
| ENSMUSG00000063087 | Gm10125 | predicted gene | lncRNA |
| ENSMUSG00000102516 | Gm38340 | predicted gene, | TEC |
| ENSMUSG00000104420 | E230020A03Rik | RIKEN cDNA E2 | TEC |
| ENSMUSG00000113894 | Gm48838 | predicted gene, | TEC |
| ENSMUSG00000026333 | Gin1 | gypsy retrotran | protein_coding |
| ENSMUSG00000114605 | C130051F05Rik | RIKEN cDNA C1 | TEC |
| ENSMUSG00000035694 | Caps2 | calcyphosphine | protein_coding |
| ENSMUSG00000105282 | Gm42981 | predicted gene | TEC |
| ENSMUSG00000105931 | Gm43014 | predicted gene | TEC |
| ENSMUSG00000110702 | Gm45767 | predicted gene | lncRNA |
| ENSMUSG00000034906 | Ncaph | non-SMC conde | protein_coding |
| ENSMUSG00000057092 | Fxyd3 | FXYD domain-c | protein_coding |
| ENSMUSG00000060212 | Pcnx2 | pecanex homol | protein_coding |
| ENSMUSG00000031998 | 1700128F08Rik | RIKEN cDNA 17 | transcribed_processe |
| ENSMUSG00000109731 | Gm45425 | predicted gene | lncRNA |
| ENSMUSG00000113167 | Gm48482 | predicted gene, | TEC |
| ENSMUSG00000053161 | Daw1 | dynein assembl | protein_coding |
| ENSMUSG00000021843 | Ktn1 | kinectin 1 | protein_coding |
| ENSMUSG00000027014 | Cwc22 | CWC22 spliceos | protein_coding |
| ENSMUSG00000112805 | Gm34574 | predicted gene, | lncRNA |
| ENSMUSG00000105332 | Gm43316 | predicted gene | TEC |
| ENSMUSG00000107300 | Gm43279 | predicted gene | TEC |
| ENSMUSG00000109481 | Gm45130 | predicted gene | TEC |
| ENSMUSG00000034022 | Cpsf1 | cleavage and p | protein_coding |
| ENSMUSG00000109032 | Gm7972 | predicted gene | transcribed_unproce |
| ENSMUSG00000078867 | Gm14418 | predicted gene | protein_coding |
| ENSMUSG00000099724 | 4930554C24Rik | RIKEN cDNA 49 | lncRNA |
| ENSMUSG00000019464 | Ptger1 | prostaglandin E | protein_coding |
| ENSMUSG00000113918 | Gm6566 | predicted gene | lncRNA |
| ENSMUSG00000068115 | Ninl | ninein-like | protein_coding |
| ENSMUSG00000085566 | A730017L22Rik | RIKEN cDNA A7 | lncRNA |
| ENSMUSG00000024883 | Rin1 | Ras and Rab int | protein_coding |
| ENSMUSG00000005237 | Dnah2 | dynein, axonem | protein_coding |
| ENSMUSG00000083899 | Gm12346 | predicted gene | transcribed_processe |
| ENSMUSG00000109644 | 0610005C13Rik | RIKEN cDNA 06 | lncRNA |
| ENSMUSG00000028608 | Czib | CXXC motif con | protein_coding |
| ENSMUSG00000017713 | Tha1 | threonine aldol | protein_coding |
| ENSMUSG00000032498 | Mlh1 | mutL homolog | protein_coding |
| ENSMUSG00000049123 | Catsperg2 | cation channel | protein_coding |
| ENSMUSG00000044211 | Gm7887 | predicted gene | processed_pseudoge |
| ENSMUSG00000114501 | Gm48582 | predicted gene, | TEC |
| ENSMUSG00000026977 | Marchf7 | membrane asso | protein_coding |
| ENSMUSG00000064032 | Gm10143 | predicted gene | lncRNA |
| ENSMUSG00000022515 | Anks3 | ankyrin repeat | protein_coding |

| | | | | | | | | |
|---|---|---|---|---|---|---|---|---|
| ENSMUSG00000050545 | Fam228b | family with seq | protein_coding | 126 | 145 | 175 | 188 | [illegible] |
| ENSMUSG00000097331 | F420014N23Rik | RIKEN cDNA F4 | lncRNA | 24 | 36 | 53 | 38 | [illegible] |
| ENSMUSG00000074449 | Potefam3a | POTE ankyrin d | protein_coding | 29 | 37 | 39 | 41 | [illegible] |
| ENSMUSG00000043833 | 2900005J15Rik | RIKEN cDNA 29 | lncRNA | 50 | 60 | 63 | 69 | [illegible] |
| ENSMUSG00000111761 | 9230112J17Rik | RIKEN cDNA 92 | lncRNA | 9 | 15 | 14 | 14 | [illegible] |
| ENSMUSG00000021098 | 4930447C04Rik | RIKEN cDNA 49 | protein_coding | 593 | 719 | 727 | 914 | [illegible] |
| ENSMUSG00000026409 | Pfkfb2 | 6-phosphofruct | protein_coding | 771 | 903 | 943 | 1,034 | [illegible] |
| ENSMUSG00000102994 | Gm37767 | predicted gene, | TEC | 0 | 1 | 3 | 7 | [illegible] |
| ENSMUSG00000057110 | Cntrl | centriolin | protein_coding | 405 | 487 | 529 | 664 | [illegible] |
| ENSMUSG00000104275 | Gm37867 | predicted gene, | lncRNA | 5 | 8 | 11 | 15 | [illegible] |
| ENSMUSG00000111990 | Gm47033 | predicted gene, | lncRNA | 37 | 45 | 62 | 72 | [illegible] |
| ENSMUSG00000004626 | Stxbp2 | syntaxin bindin | protein_coding | 277 | 385 | 409 | 501 | [illegible] |
| ENSMUSG00000096822 | Or1j15 | olfactory recep | protein_coding | 61 | 86 | 84 | 112 | [illegible] |
| ENSMUSG00000102813 | Gm37795 | predicted gene, | TEC | 15 | 22 | 32 | 36 | [illegible] |
| ENSMUSG00000105700 | Gm42772 | predicted gene | TEC | 0 | 6 | 6 | 12 | [illegible] |
| ENSMUSG00000103831 | Gm37608 | predicted gene, | TEC | 15 | 30 | 33 | 33 | [illegible] |
| ENSMUSG00000103047 | Gm37310 | predicted gene, | TEC | 19 | 38 | 44 | 51 | [illegible] |
| ENSMUSG00000106832 | Gm42632 | predicted gene | TEC | 11 | 17 | 18 | 19 | [illegible] |
| ENSMUSG00000078190 | Dnm3os | dynamin 3, opp | lncRNA | 8 | 12 | 25 | 18 | [illegible] |
| ENSMUSG00000097619 | 4833422M21Ri | RIKEN cDNA 48 | lncRNA | 16 | 31 | 33 | 39 | [illegible] |
| ENSMUSG00000078813 | Leng1 | leukocyte recep | protein_coding | 226 | 233 | 273 | 259 | [illegible] |
| ENSMUSG00000070000 | Fcho1 | FCH domain on | protein_coding | 1,123 | 1,186 | 1,280 | 1,386 | [illegible] |
| ENSMUSG00000110529 | Gm45694 | predicted gene | TEC | 31 | 42 | 48 | 61 | [illegible] |
| ENSMUSG00000115837 | 9130002K18Rik | RIKEN cDNA 91 | TEC | 9 | 24 | 12 | 19 | [illegible] |
| ENSMUSG00000015501 | Hivep2 | human immuno | protein_coding | [illegible] | [illegible] | [illegible] | [illegible] | [illegible] |
| ENSMUSG00000112074 | Gm48278 | predicted gene, | lncRNA | [illegible] | [illegible] | [illegible] | [illegible] | [illegible] |
| ENSMUSG00000004567 | Mcoln1 | mucolipin 1 | protein_coding | [illegible] | [illegible] | [illegible] | [illegible] | [illegible] |
| ENSMUSG00000085037 | 4933421O10Rik | RIKEN cDNA 49 | lncRNA | [illegible] | [illegible] | [illegible] | [illegible] | [illegible] |
| ENSMUSG00000106819 | Gm43498 | predicted gene | processed_pseudoge | [illegible] | [illegible] | [illegible] | [illegible] | [illegible] |
| ENSMUSG00000107320 | Gm42549 | predicted gene | TEC | [illegible] | [illegible] | [illegible] | [illegible] | [illegible] |
| ENSMUSG00000105416 | Gm42553 | predicted gene | TEC | [illegible] | [illegible] | [illegible] | [illegible] | [illegible] |
| ENSMUSG00000106986 | Gm42887 | predicted gene | TEC | [illegible] | [illegible] | [illegible] | [illegible] | [illegible] |
| ENSMUSG00000024170 | Telo2 | telomere maint | protein_coding | [illegible] | [illegible] | [illegible] | [illegible] | [illegible] |
| ENSMUSG00000026970 | Rbms1 | RNA binding m | protein_coding | [illegible] | [illegible] | [illegible] | [illegible] | [illegible] |
| ENSMUSG00000047789 | Slc38a9 | solute carrier f | protein_coding | [illegible] | [illegible] | [illegible] | [illegible] | [illegible] |
| ENSMUSG00000022070 | Bora | bora, aurora kir | protein_coding | [illegible] | [illegible] | [illegible] | [illegible] | [illegible] |
| ENSMUSG00000037355 | Uvssa | UV stimulated s | protein_coding | [illegible] | [illegible] | [illegible] | [illegible] | [illegible] |
| ENSMUSG00000091945 | Vmn2r114 | vomeronasal 2, | protein_coding | [illegible] | [illegible] | [illegible] | [illegible] | [illegible] |
| ENSMUSG00000108264 | Gm20371 | predicted gene, | TEC | [illegible] | [illegible] | [illegible] | [illegible] | [illegible] |
| ENSMUSG00000086607 | 4930511M06Ri | RIKEN cDNA 49 | lncRNA | [illegible] | [illegible] | [illegible] | [illegible] | [illegible] |
| ENSMUSG00000117853 | Vmn1r88 | vomeronasal 1 | protein_coding | [illegible] | [illegible] | [illegible] | [illegible] | [illegible] |
| ENSMUSG00000105530 | Gm43829 | predicted gene | TEC | [illegible] | [illegible] | [illegible] | [illegible] | [illegible] |
| ENSMUSG00000109551 | 4930435N07Rik | RIKEN cDNA 49 | TEC | [illegible] | [illegible] | [illegible] | [illegible] | [illegible] |
| ENSMUSG00000068950 | Or1j10 | olfactory recep | protein_coding | [illegible] | [illegible] | [illegible] | [illegible] | [illegible] |
| ENSMUSG00000102136 | Gm37107 | predicted gene, | lncRNA | [illegible] | [illegible] | [illegible] | [illegible] | [illegible] |
| ENSMUSG00000103630 | Gm37242 | predicted gene, | TEC | [illegible] | [illegible] | [illegible] | [illegible] | [illegible] |
| ENSMUSG00000112169 | Gm48536 | predicted gene, | TEC | [illegible] | [illegible] | [illegible] | [illegible] | [illegible] |
| ENSMUSG00000034610 | Tut4 | terminal uridyly | protein_coding | [illegible] | [illegible] | [illegible] | [illegible] | [illegible] |
| ENSMUSG00000106386 | Gm42991 | predicted gene | TEC | [illegible] | [illegible] | [illegible] | [illegible] | [illegible] |
| ENSMUSG00000021182 | Ccdc88c | coiled-coil dom | protein_coding | [illegible] | [illegible] | [illegible] | [illegible] | [illegible] |
| ENSMUSG00000107826 | Gm44242 | predicted gene, | TEC | [illegible] | [illegible] | [illegible] | [illegible] | [illegible] |
| ENSMUSG00000028653 | Trit1 | tRNA isopenten | protein_coding | [illegible] | [illegible] | [illegible] | [illegible] | [illegible] |
| ENSMUSG00000113640 | Adat3 | adenosine dear | protein_coding | [illegible] | [illegible] | [illegible] | [illegible] | [illegible] |
| ENSMUSG00000027208 | Fgf7 | fibroblast grow | protein_coding | [illegible] | [illegible] | [illegible] | [illegible] | [illegible] |
| ENSMUSG00000070458 | Vmn2r73 | vomeronasal 2, | protein_coding | [illegible] | [illegible] | [illegible] | [illegible] | [illegible] |
| ENSMUSG00000105549 | Gm43540 | predicted gene | processed_pseudoge | [illegible] | [illegible] | [illegible] | [illegible] | [illegible] |
| ENSMUSG00000105974 | Gm43318 | predicted gene | TEC | [illegible] | [illegible] | [illegible] | [illegible] | [illegible] |
| ENSMUSG00000117819 | Gm50253 | predicted gene, | protein_coding | [illegible] | [illegible] | [illegible] | [illegible] | [illegible] |
| ENSMUSG00000032816 | Igdcc4 | immunoglobuli | protein_coding | [illegible] | [illegible] | [illegible] | [illegible] | [illegible] |
| ENSMUSG00000052296 | Ppp6r1 | protein phosph | protein_coding | [illegible] | [illegible] | [illegible] | [illegible] | [illegible] |
| ENSMUSG00000109916 | 2410039M03Ri | RIKEN cDNA 24 | unprocessed_pseudo | [illegible] | [illegible] | [illegible] | [illegible] | [illegible] |
| ENSMUSG00000028681 | Ptch2 | patched 2 | protein_coding | [illegible] | [illegible] | [illegible] | [illegible] | [illegible] |
| ENSMUSG00000109368 | Gm45015 | predicted gene | protein_coding | [illegible] | [illegible] | [illegible] | [illegible] | [illegible] |
| ENSMUSG00000105063 | Gm43488 | predicted gene | TEC | [illegible] | [illegible] | [illegible] | [illegible] | [illegible] |
| ENSMUSG00000117815 | Gm50166 | predicted gene, | lncRNA | [illegible] | [illegible] | [illegible] | [illegible] | [illegible] |
| ENSMUSG00000109378 | Gm49396 | predicted gene, | protein_coding | [illegible] | [illegible] | [illegible] | [illegible] | [illegible] |
| ENSMUSG00000111236 | Gm31614 | predicted gene, | lncRNA | [illegible] | [illegible] | [illegible] | [illegible] | [illegible] |
| ENSMUSG00000050954 | Zfp169 | zinc finger prot | protein_coding | [illegible] | [illegible] | [illegible] | [illegible] | [illegible] |
| ENSMUSG00000071665 | Foxr2 | forkhead box R | protein_coding | [illegible] | [illegible] | [illegible] | [illegible] | [illegible] |
| ENSMUSG00000107546 | Gm20560 | predicted gene, | lncRNA | [illegible] | [illegible] | [illegible] | [illegible] | [illegible] |
| ENSMUSG00000029202 | Pds5a | PDS5 cohesin a | protein_coding | [illegible] | [illegible] | [illegible] | [illegible] | [illegible] |
| ENSMUSG00000032175 | Tyk2 | tyrosine kinase | protein_coding | [illegible] | [illegible] | [illegible] | [illegible] | [illegible] |
| ENSMUSG00000051037 | Zfp455 | zinc finger prot | protein_coding | [illegible] | [illegible] | [illegible] | [illegible] | [illegible] |
| ENSMUSG00000045106 | Ccdc73 | coiled-coil dom | protein_coding | [illegible] | [illegible] | [illegible] | [illegible] | [illegible] |
| ENSMUSG00000029607 | Ankrd61 | ankyrin repeat | protein_coding | [illegible] | [illegible] | [illegible] | [illegible] | [illegible] |
| ENSMUSG00000030322 | Mbd4 | methyl-CpG bin | protein_coding | [illegible] | [illegible] | [illegible] | [illegible] | [illegible] |
| ENSMUSG00000097923 | Gm26577 | predicted gene, | lncRNA | [illegible] | [illegible] | [illegible] | [illegible] | [illegible] |
| ENSMUSG00000104391 | Gm37960 | predicted gene, | TEC | [illegible] | [illegible] | [illegible] | [illegible] | [illegible] |
| ENSMUSG00000112844 | Gm47036 | predicted gene, | processed_pseudoge | [illegible] | [illegible] | [illegible] | [illegible] | [illegible] |
| ENSMUSG00000097231 | Gm26852 | predicted gene, | lncRNA | [illegible] | [illegible] | [illegible] | [illegible] | [illegible] |
| ENSMUSG00000103685 | Gm37074 | predicted gene, | TEC | [illegible] | [illegible] | [illegible] | [illegible] | [illegible] |
| ENSMUSG00000103693 | Gm37529 | predicted gene, | lncRNA | [illegible] | [illegible] | [illegible] | [illegible] | [illegible] |
| ENSMUSG00000105469 | A930036I15Rik | RIKEN cDNA A9 | TEC | [illegible] | [illegible] | [illegible] | [illegible] | [illegible] |
| ENSMUSG00000038708 | Golga4 | golgi autoantig | protein_coding | [illegible] | [illegible] | [illegible] | [illegible] | [illegible] |
| ENSMUSG00000073116 | Gm10471 | predicted gene | protein_coding | [illegible] | [illegible] | [illegible] | [illegible] | [illegible] |
| ENSMUSG00000040850 | Psme4 | proteasome (pr | protein_coding | [illegible] | [illegible] | [illegible] | [illegible] | [illegible] |
| ENSMUSG00000027365 | Trpm7 | transient recep | protein_coding | [illegible] | [illegible] | [illegible] | [illegible] | [illegible] |
| ENSMUSG00000034973 | Dop1a | DOP1 leucine zi | protein_coding | [illegible] | [illegible] | [illegible] | [illegible] | [illegible] |
| ENSMUSG00000103983 | Gm20045 | predicted gene, | lncRNA | [illegible] | [illegible] | [illegible] | [illegible] | [illegible] |
| ENSMUSG00000107577 | Gm44103 | predicted gene, | TEC | [illegible] | [illegible] | [illegible] | [illegible] | [illegible] |
| ENSMUSG00000115027 | Vmn1r81 | vomeronasal 1 | protein_coding | [illegible] | [illegible] | [illegible] | [illegible] | [illegible] |
| ENSMUSG00000115105 | Gm49238 | predicted gene, | TEC | [illegible] | [illegible] | [illegible] | [illegible] | [illegible] |
| ENSMUSG00000037593 | Rskr | ribosomal prot | protein_coding | [illegible] | [illegible] | [illegible] | [illegible] | [illegible] |
| ENSMUSG00000040860 | Crocc | ciliary rootlet c | protein_coding | [illegible] | [illegible] | [illegible] | [illegible] | [illegible] |
| ENSMUSG00000028093 | Acp6 | acid phosphata | protein_coding | [illegible] | [illegible] | [illegible] | [illegible] | [illegible] |
| ENSMUSG00000084897 | Gm14226 | predicted gene | lncRNA | [illegible] | [illegible] | [illegible] | [illegible] | [illegible] |
| ENSMUSG00000036882 | Arhgap33 | Rho GTPase act | protein_coding | [illegible] | [illegible] | [illegible] | [illegible] | [illegible] |
| ENSMUSG00000104027 | Gm37211 | predicted gene, | TEC | [illegible] | [illegible] | [illegible] | [illegible] | [illegible] |
| ENSMUSG00000109094 | Gm44587 | predicted gene | TEC | [illegible] | [illegible] | [illegible] | [illegible] | [illegible] |
| ENSMUSG00000028456 | Unc13b | unc-13 homolo | protein_coding | [illegible] | [illegible] | [illegible] | [illegible] | [illegible] |
| ENSMUSG00000107306 | Gm42577 | predicted gene | TEC | [illegible] | [illegible] | [illegible] | [illegible] | [illegible] |
| ENSMUSG00000033257 | Ttll4 | tubulin tyrosine | protein_coding | [illegible] | [illegible] | [illegible] | [illegible] | [illegible] |
| ENSMUSG00000062470 | Fbxl12os | F-box and leuci | lncRNA | [illegible] | [illegible] | [illegible] | [illegible] | [illegible] |
| ENSMUSG00000034771 | Tle2 | transducin-like | protein_coding | [illegible] | [illegible] | [illegible] | [illegible] | [illegible] |
| ENSMUSG00000070427 | Il18bp | interleukin 18 b | protein_coding | [illegible] | [illegible] | [illegible] | [illegible] | [illegible] |
| ENSMUSG00000032582 | Rbm6 | RNA binding m | protein_coding | [illegible] | [illegible] | [illegible] | [illegible] | [illegible] |
| ENSMUSG00000113850 | Gm47191 | predicted gene, | transcribed_processe | [illegible] | [illegible] | [illegible] | [illegible] | [illegible] |
| ENSMUSG00000053483 | Usp21 | ubiquitin specif | protein_coding | [illegible] | [illegible] | [illegible] | [illegible] | [illegible] |
| ENSMUSG00000078984 | Gm11027 | predicted gene | protein_coding | [illegible] | [illegible] | [illegible] | [illegible] | [illegible] |
| ENSMUSG00000102816 | Gm37814 | predicted gene, | lncRNA | [illegible] | [illegible] | [illegible] | [illegible] | [illegible] |
| ENSMUSG00000083579 | Gm15538 | predicted gene | processed_pseudoge | [illegible] | [illegible] | [illegible] | [illegible] | [illegible] |
| ENSMUSG00000020703 | 5530401A14Rik | RIKEN cDNA 55 | lncRNA | [illegible] | [illegible] | [illegible] | [illegible] | [illegible] |
| ENSMUSG00000114345 | Gm10873 | predicted gene | lncRNA | [illegible] | [illegible] | [illegible] | [illegible] | [illegible] |
| ENSMUSG00000038342 | Mlxip | MLX interacting | protein_coding | [illegible] | [illegible] | [illegible] | [illegible] | [illegible] |
| ENSMUSG00000053552 | Ebf4 | early B cell fact | protein_coding | [illegible] | [illegible] | [illegible] | [illegible] | [illegible] |
| ENSMUSG00000067629 | Syngap1 | synaptic Ras GT | protein_coding | [illegible] | [illegible] | [illegible] | [illegible] | [illegible] |
| ENSMUSG00000060206 | Zfp462 | zinc finger prot | protein_coding | [illegible] | [illegible] | [illegible] | [illegible] | [illegible] |
| ENSMUSG00000004113 | Cacna1b | calcium channe | protein_coding | [illegible] | [illegible] | [illegible] | [illegible] | [illegible] |
| ENSMUSG00000027309 | Dnaaf9 | dynein axonem | protein_coding | [illegible] | [illegible] | [illegible] | [illegible] | [illegible] |
| ENSMUSG00000104069 | Gm37198 | predicted gene, | TEC | [illegible] | [illegible] | [illegible] | [illegible] | [illegible] |
| ENSMUSG00000029036 | Atad3a | ATPase family, | protein_coding | [illegible] | [illegible] | [illegible] | [illegible] | [illegible] |
| ENSMUSG00000097464 | Gm26736 | predicted gene, | TEC | [illegible] | [illegible] | [illegible] | [illegible] | [illegible] |
| ENSMUSG00000021819 | Zswim8 | zinc finger SWI | protein_coding | [illegible] | [illegible] | [illegible] | [illegible] | [illegible] |
| ENSMUSG00000050266 | Or52b1 | olfactory recep | protein_coding | [illegible] | [illegible] | [illegible] | [illegible] | [illegible] |
| ENSMUSG00000010097 | Nxf1 | nuclear RNA ex | protein_coding | [illegible] | [illegible] | [illegible] | [illegible] | [illegible] |
| ENSMUSG00000048703 | Speer4b | spermatogenes | protein_coding | [illegible] | [illegible] | [illegible] | [illegible] | [illegible] |
| ENSMUSG00000104724 | Gm43162 | predicted gene | TEC | [illegible] | [illegible] | [illegible] | [illegible] | [illegible] |
| ENSMUSG00000019763 | Rmnd1 | required for me | protein_coding | [illegible] | [illegible] | [illegible] | [illegible] | [illegible] |
| ENSMUSG00000034675 | Dbn1 | drebrin 1 | protein_coding | [illegible] | [illegible] | [illegible] | [illegible] | [illegible] |
| ENSMUSG00000034818 | Celf5 | CUGBP, Elav-lik | protein_coding | [illegible] | [illegible] | [illegible] | [illegible] | [illegible] |
| ENSMUSG00000091071 | 1700030C10Rik | RIKEN cDNA 17 | transcribed_unproce | [illegible] | [illegible] | [illegible] | [illegible] | [illegible] |
| ENSMUSG00000097099 | Gm9917 | predicted gene | lncRNA | [illegible] | [illegible] | [illegible] | [illegible] | [illegible] |
| ENSMUSG00000083084 | Gm14667 | predicted gene | processed_pseudoge | [illegible] | [illegible] | [illegible] | [illegible] | [illegible] |
| ENSMUSG00000104916 | Gm42830 | predicted gene | TEC | [illegible] | [illegible] | [illegible] | [illegible] | [illegible] |
| ENSMUSG00000062861 | Zfp28 | zinc finger prot | protein_coding | [illegible] | [illegible] | [illegible] | [illegible] | [illegible] |
| ENSMUSG00000035270 | Impg2 | interphotorece | protein_coding | [illegible] | [illegible] | [illegible] | [illegible] | [illegible] |
| ENSMUSG00000111374 | Gm50490 | predicted gene, | processed_pseudoge | [illegible] | [illegible] | [illegible] | [illegible] | [illegible] |
| ENSMUSG00000023942 | Slc29a1 | solute carrier f | protein_coding | [illegible] | [illegible] | [illegible] | [illegible] | [illegible] |
| ENSMUSG00000111063 | Zfp660 | zinc finger prot | lncRNA | [illegible] | [illegible] | [illegible] | [illegible] | [illegible] |
| ENSMUSG00000003233 | Dvl3 | dishevelled seg | protein_coding | [illegible] | [illegible] | [illegible] | [illegible] | [illegible] |
| ENSMUSG00000045103 | Dmd | dystrophin, mu | protein_coding | [illegible] | [illegible] | [illegible] | [illegible] | [illegible] |
| ENSMUSG00000081752 | Sms-ps | spermine synth | processed_pseudoge | [illegible] | [illegible] | [illegible] | [illegible] | [illegible] |
| ENSMUSG00000022313 | Utp23 | UTP23 small su | protein_coding | [illegible] | [illegible] | [illegible] | [illegible] | [illegible] |
| ENSMUSG00000020018 | Snrpf | small nuclear ri | protein_coding | [illegible] | [illegible] | [illegible] | [illegible] | [illegible] |
| ENSMUSG00000023845 | Lnpep | leucyl/cystinyl a | protein_coding | [illegible] | [illegible] | [illegible] | [illegible] | [illegible] |
| ENSMUSG00000025226 | Fbxl15 | F-box and leuci | protein_coding | [illegible] | [illegible] | [illegible] | [illegible] | [illegible] |
| ENSMUSG00000078377 | Gm4294 | predicted gene | processed_pseudoge | [illegible] | [illegible] | [illegible] | [illegible] | [illegible] |
| ENSMUSG00000038763 | Alpk3 | alpha-kinase 3 | protein_coding | [illegible] | [illegible] | [illegible] | [illegible] | [illegible] |
| ENSMUSG00000021250 | Fos | FBJ osteosarcor | protein_coding | [illegible] | [illegible] | [illegible] | [illegible] | [illegible] |
| ENSMUSG00000022553 | Maf1 | MAF1 homolog | protein_coding | [illegible] | [illegible] | [illegible] | [illegible] | [illegible] |
| ENSMUSG00000000184 | Ccnd2 | cyclin D2 | protein_coding | [illegible] | [illegible] | [illegible] | [illegible] | [illegible] |
| ENSMUSG00000020458 | Rtn4 | reticulon 4 | protein_coding | [illegible] | [illegible] | [illegible] | [illegible] | [illegible] |
| ENSMUSG00000015478 | Rnf5 | ring finger prot | protein_coding | [illegible] | [illegible] | [illegible] | [illegible] | [illegible] |
| ENSMUSG00000052214 | Opa3 | optic atrophy 3 | protein_coding | [illegible] | [illegible] | [illegible] | [illegible] | [illegible] |
| ENSMUSG00000030600 | Lrfn1 | leucine rich rep | protein_coding | [illegible] | [illegible] | [illegible] | [illegible] | [illegible] |
| ENSMUSG00000044628 | Rnf208 | ring finger prot | protein_coding | [illegible] | [illegible] | [illegible] | [illegible] | [illegible] |
| ENSMUSG00000003072 | Atp5d | ATP synthase, H | protein_coding | [illegible] | [illegible] | [illegible] | [illegible] | [illegible] |
| ENSMUSG00000025228 | Actr1a | ARP1 actin-rela | protein_coding | [illegible] | [illegible] | [illegible] | [illegible] | [illegible] |
| ENSMUSG00000032997 | Chpf | chondroitin pol | protein_coding | [illegible] | [illegible] | [illegible] | [illegible] | [illegible] |
| ENSMUSG00000003469 | Phyhip | phytanoyl-CoA | protein_coding | [illegible] | [illegible] | [illegible] | [illegible] | [illegible] |
| ENSMUSG00000039615 | Stub1 | STIP1 homology | protein_coding | [illegible] | [illegible] | [illegible] | [illegible] | [illegible] |
| ENSMUSG00000024121 | Atp6v0c | ATPase, H+ trar | protein_coding | [illegible] | [illegible] | [illegible] | [illegible] | [illegible] |
| ENSMUSG00000058756 | Thra | thyroid hormor | protein_coding | [illegible] | [illegible] | [illegible] | [illegible] | [illegible] |
| ENSMUSG00000079144 | A130010J15Rik | RIKEN cDNA A1 | protein_coding | [illegible] | [illegible] | [illegible] | [illegible] | [illegible] |
| ENSMUSG00000062825 | Actg1 | actin, gamma, c | protein_coding | [illegible] | [illegible] | [illegible] | [illegible] | [illegible] |
| ENSMUSG00000018846 | Pank3 | pantothenate k | protein_coding | [illegible] | [illegible] | [illegible] | [illegible] | [illegible] |
| ENSMUSG00000019124 | Scrn1 | secernin 1 | protein_coding | [illegible] | [illegible] | [illegible] | [illegible] | [illegible] |
| ENSMUSG00000067787 | Blcap | bladder cancer | protein_coding | [illegible] | [illegible] | [illegible] | [illegible] | [illegible] |
| ENSMUSG00000073062 | Zxdb | zinc finger, X-lir | protein_coding | [illegible] | [illegible] | [illegible] | [illegible] | [illegible] |
| ENSMUSG00000030410 | Dmwd | dystrophia myo | protein_coding | [illegible] | [illegible] | [illegible] | [illegible] | [illegible] |
| ENSMUSG00000020496 | Rnf187 | ring finger prot | protein_coding | [illegible] | [illegible] | [illegible] | [illegible] | [illegible] |
| ENSMUSG00000021395 | Spin1 | spindlin 1 | protein_coding | [illegible] | [illegible] | [illegible] | [illegible] | [illegible] |
| ENSMUSG00000033953 | Ppp3r1 | protein phosph | protein_coding | [illegible] | [illegible] | [illegible] | [illegible] | [illegible] |
| ENSMUSG00000034723 | Tmx4 | thioredoxin-rel | protein_coding | [illegible] | [illegible] | [illegible] | [illegible] | [illegible] |
| ENSMUSG00000030327 | Necap1 | NECAP endocyt | protein_coding | [illegible] | [illegible] | [illegible] | [illegible] | [illegible] |
| ENSMUSG00000047388 | Atmin | ATM interactor | protein_coding | [illegible] | [illegible] | [illegible] | [illegible] | [illegible] |
| ENSMUSG00000030842 | Lamtor1 | late endosomal | protein_coding | [illegible] | [illegible] | [illegible] | [illegible] | [illegible] |
| ENSMUSG00000004637 | Wwox | WW domain-co | protein_coding | [illegible] | [illegible] | [illegible] | [illegible] | [illegible] |
| ENSMUSG00000044147 | Arf6 | ADP-ribosylatio | protein_coding | [illegible] | [illegible] | [illegible] | [illegible] | [illegible] |
| ENSMUSG00000026159 | Agfg1 | ArfGAP with FG | protein_coding | [illegible] | [illegible] | [illegible] | [illegible] | [illegible] |
| ENSMUSG00000028833 | Ncdn | neurochondrin | protein_coding | [illegible] | [illegible] | [illegible] | [illegible] | [illegible] |
| ENSMUSG00000002379 | Ndufa11 | NADH:ubiquino | protein_coding | [illegible] | [illegible] | [illegible] | [illegible] | [illegible] |
| ENSMUSG00000014956 | Ppp1cb | protein phosph | protein_coding | [illegible] | [illegible] | [illegible] | [illegible] | [illegible] |
| ENSMUSG00000028373 | Astn2 | astrotactin 2 | protein_coding | [illegible] | [illegible] | [illegible] | [illegible] | [illegible] |
| ENSMUSG00000037049 | Smpd1 | sphingomyelin | protein_coding | [illegible] | [illegible] | [illegible] | [illegible] | [illegible] |
| ENSMUSG00000063873 | Slc24a3 | solute carrier f | protein_coding | [illegible] | [illegible] | [illegible] | [illegible] | [illegible] |
| ENSMUSG00000067279 | Ppp1r3c | protein phosph | protein_coding | [illegible] | [illegible] | [illegible] | [illegible] | [illegible] |
| ENSMUSG00000022619 | Mapk8ip2 | mitogen-activa | protein_coding | [illegible] | [illegible] | [illegible] | [illegible] | [illegible] |
| ENSMUSG00000024261 | Syt4 | synaptotagmin | protein_coding | [illegible] | [illegible] | [illegible] | [illegible] | [illegible] |
| ENSMUSG00000027327 | 1700037H04Rik | RIKEN cDNA 17 | protein_coding | [illegible] | [illegible] | [illegible] | [illegible] | [illegible] |
| ENSMUSG00000032875 | Arhgef17 | Rho guanine nu | protein_coding | [illegible] | [illegible] | [illegible] | [illegible] | [illegible] |
| ENSMUSG00000028953 | Abcf2 | ATP-binding cas | protein_coding | [illegible] | [illegible] | [illegible] | [illegible] | [illegible] |
| ENSMUSG00000046541 | Zfp526 | zinc finger prot | protein_coding | [illegible] | [illegible] | [illegible] | [illegible] | [illegible] |
| ENSMUSG00000026080 | Chst10 | carbohydrate s | protein_coding | [illegible] | [illegible] | [illegible] | [illegible] | [illegible] |
| ENSMUSG00000042229 | Rabif | RAB interacting | protein_coding | [illegible] | [illegible] | [illegible] | [illegible] | [illegible] |
| ENSMUSG00000029557 | Mrm2 | mitochondrial r | protein_coding | [illegible] | [illegible] | [illegible] | [illegible] | [illegible] |
| ENSMUSG00000031513 | Leprotl1 | leptin receptor | protein_coding | [illegible] | [illegible] | [illegible] | [illegible] | [illegible] |
| ENSMUSG00000031988 | Vps26b | VPS26 retrome | protein_coding | [illegible] | [illegible] | [illegible] | [illegible] | [illegible] |
| ENSMUSG00000001729 | Akt1 | thymoma viral | protein_coding | [illegible] | [illegible] | [illegible] | [illegible] | [illegible] |
| ENSMUSG00000038602 | Slc35f1 | solute carrier f | protein_coding | [illegible] | [illegible] | [illegible] | [illegible] | [illegible] |
| ENSMUSG00000070509 | Rgma | repulsive guida | protein_coding | [illegible] | [illegible] | [illegible] | [illegible] | [illegible] |
| ENSMUSG00000028525 | Pde4b | phosphodiester | protein_coding | [illegible] | [illegible] | [illegible] | [illegible] | [illegible] |
| ENSMUSG00000024892 | Pcx | pyruvate carbo | protein_coding | [illegible] | [illegible] | [illegible] | [illegible] | [illegible] |
| ENSMUSG00000029924 | Slc37a3 | solute carrier f | protein_coding | [illegible] | [illegible] | [illegible] | [illegible] | [illegible] |
| ENSMUSG00000019785 | Clvs2 | clavesin 2 | protein_coding | [illegible] | [illegible] | [illegible] | [illegible] | [illegible] |
| ENSMUSG00000034118 | Tpst1 | protein-tyrosin | protein_coding | [illegible] | [illegible] | [illegible] | [illegible] | [illegible] |
| ENSMUSG00000026869 | Psmd5 | proteasome (pr | protein_coding | [illegible] | [illegible] | [illegible] | [illegible] | [illegible] |
| ENSMUSG00000034958 | Atcay | ataxia, cerebell | protein_coding | [illegible] | [illegible] | [illegible] | [illegible] | [illegible] |
| ENSMUSG00000019731 | Slc35e1 | solute carrier f | protein_coding | [illegible] | [illegible] | [illegible] | [illegible] | [illegible] |
| ENSMUSG00000024767 | Otub1 | OTU domain, ul | protein_coding | [illegible] | [illegible] | [illegible] | [illegible] | [illegible] |
| ENSMUSG00000020261 | Slc36a1 | solute carrier f | protein_coding | [illegible] | [illegible] | [illegible] | [illegible] | [illegible] |
| ENSMUSG00000038582 | Pptc7 | PTC7 protein pl | protein_coding | [illegible] | [illegible] | [illegible] | [illegible] | [illegible] |
| ENSMUSG00000042444 | Mindy2 | MINDY lysine 4 | protein_coding | [illegible] | [illegible] | [illegible] | [illegible] | [illegible] |
| ENSMUSG00000039485 | Tspyl4 | TSPY-like 4 | protein_coding | [illegible] | [illegible] | [illegible] | [illegible] | [illegible] |
| ENSMUSG00000048027 | Rgmb | repulsive guida | protein_coding | [illegible] | [illegible] | [illegible] | [illegible] | [illegible] |
| ENSMUSG00000025786 | Zdhhc3 | zinc finger, DHH | protein_coding | [illegible] | [illegible] | [illegible] | [illegible] | [illegible] |
| ENSMUSG00000007812 | Zfp655 | zinc finger prot | protein_coding | [illegible] | [illegible] | [illegible] | [illegible] | [illegible] |
| ENSMUSG00000048732 | Klhl11 | kelch-like 11 | protein_coding | [illegible] | [illegible] | [illegible] | [illegible] | [illegible] |
| ENSMUSG00000001687 | Ubl3 | ubiquitin-like 3 | protein_coding | [illegible] | [illegible] | [illegible] | [illegible] | [illegible] |
| ENSMUSG00000041028 | Ghitm | growth hormor | protein_coding | [illegible] | [illegible] | [illegible] | [illegible] | [illegible] |
| ENSMUSG00000052738 | Suclg1 | succinate-CoA l | protein_coding | [illegible] | [illegible] | [illegible] | [illegible] | [illegible] |
| ENSMUSG00000017466 | Timp2 | tissue inhibitor | protein_coding | [illegible] | [illegible] | [illegible] | [illegible] | [illegible] |
| ENSMUSG00000041697 | Cox6a1 | cytochrome c o | protein_coding | [illegible] | [illegible] | [illegible] | [illegible] | [illegible] |
| ENSMUSG00000071379 | Hpcal1 | hippocalcin-like | protein_coding | [illegible] | [illegible] | [illegible] | [illegible] | [illegible] |
| ENSMUSG00000027673 | Ndufb5 | NADH:ubiquino | protein_coding | [illegible] | [illegible] | [illegible] | [illegible] | [illegible] |
| ENSMUSG00000033174 | Mgll | monoglyceride | protein_coding | [illegible] | [illegible] | [illegible] | [illegible] | [illegible] |
| ENSMUSG00000043445 | Pgp | phosphoglycola | protein_coding | [illegible] | [illegible] | [illegible] | [illegible] | [illegible] |
| ENSMUSG00000056234 | Ncoa4 | nuclear recepto | protein_coding | [illegible] | [illegible] | [illegible] | [illegible] | [illegible] |
| ENSMUSG00000035150 | Eif2s3x | eukaryotic tran | protein_coding | [illegible] | [illegible] | [illegible] | [illegible] | [illegible] |
| ENSMUSG00000036158 | Prickle1 | prickle planar c | protein_coding | [illegible] | [illegible] | [illegible] | [illegible] | [illegible] |
| ENSMUSG00000028756 | Pink1 | PTEN induced p | protein_coding | [illegible] | [illegible] | [illegible] | [illegible] | [illegible] |
| ENSMUSG00000021816 | Ppp3cb | protein phosph | protein_coding | [illegible] | [illegible] | [illegible] | [illegible] | [illegible] |
| ENSMUSG00000030307 | Slc6a11 | solute carrier f | protein_coding | [illegible] | [illegible] | [illegible] | [illegible] | [illegible] |
| ENSMUSG00000007038 | Neu1 | neuraminidase | protein_coding | [illegible] | [illegible] | [illegible] | [illegible] | [illegible] |
| ENSMUSG00000027599 | Armc1 | armadillo repea | protein_coding | [illegible] | [illegible] | [illegible] | [illegible] | [illegible] |
| ENSMUSG00000023004 | Tuba1b | tubulin, alpha 1 | protein_coding | [illegible] | [illegible] | [illegible] | [illegible] | [illegible] |
| ENSMUSG00000024758 | Rtn3 | reticulon 3 | protein_coding | [illegible] | [illegible] | [illegible] | [illegible] | [illegible] |
| ENSMUSG00000005683 | Cs | citrate synthase | protein_coding | [illegible] | [illegible] | [illegible] | [illegible] | [illegible] |
| ENSMUSG00000051675 | Trim32 | tripartite motif | protein_coding | [illegible] | [illegible] | [illegible] | [illegible] | [illegible] |
| ENSMUSG00000071649 | B3gat3 | beta-1,3-glucur | protein_coding | [illegible] | [illegible] | [illegible] | [illegible] | [illegible] |

| | | | | | | | | | | | | | | | | |
|---|---|---|---|---|---|---|---|---|---|---|---|---|---|---|---|---|
| ENSMUSG00000024065 | Ehd3 | EH-domain con | protein_coding | 2,207 | 2,055 | 2,198 | 1,992 | 1,900 | 1,920 | 1,847 | 1,975 | 1,907 | 1,972 | 1,910 | 1,931 | -1.1 | 0.000834 |
| ENSMUSG00000048022 | Tmem229a | transmembran | protein_coding | 3,479 | 3,260 | 3,518 | 3,054 | 2,878 | 2,936 | 2,858 | 3,006 | 2,970 | 2,911 | 2,870 | 2,988 | -1.14 | 0.00393 |
| ENSMUSG00000018865 | Sult4a1 | sulfotransferas | protein_coding | 7,145 | 6,749 | 6,975 | 6,221 | 6,137 | 5,852 | 5,759 | 6,210 | 6,152 | 6,119 | 5,786 | 5,981 | -1.13 | 0.00132 |
| ENSMUSG00000022415 | Syngr1 | synaptogyrin 1 | protein_coding | 11,423 | 11,175 | 11,270 | 10,313 | 10,000 | 9,713 | 9,434 | 9,934 | 10,146 | 10,148 | 9,456 | 9,846 | -1.13 | 0.00112 |
| ENSMUSG00000031913 | Vps4a | vacuolar protei | protein_coding | 2,326 | 2,289 | 2,264 | 2,158 | 2,080 | 1,954 | 2,013 | 2,143 | 2,100 | 2,105 | 1,980 | 2,091 | -1.1 | 0.00169 |
| ENSMUSG00000039953 | Clstn1 | calsyntenin 1 | protein_coding | 21,024 | 20,439 | 20,894 | 19,075 | 18,874 | 17,438 | 18,115 | 18,828 | 18,180 | 18,754 | 17,958 | 18,729 | -1.11 | 0.000269 |
| ENSMUSG00000037341 | Slc9a7 | solute carrier fa | protein_coding | 1,252 | 1,155 | 1,295 | 1,199 | 1,005 | 1,072 | 1,010 | 1,115 | 1,127 | 1,104 | 1,031 | 1,019 | -1.16 | 0.000199 |
| ENSMUSG00000051412 | | | | 918 | 855 | 1,029 | 906 | 783 | 825 | 806 | 869 | 875 | 813 | 789 | 787 | -1.13 | 0.00828 |
| ENSMUSG00000019297 | Nop9 | NOP9 nucleolar | protein_coding | 241 | 274 | 350 | 195 | 181 | 168 | 192 | 220 | 253 | 176 | 148 | 156 | -1.38 | 0.0065 |
| ENSMUSG00000032279 | Idh3a | isocitrate dehy | protein_coding | 5,143 | 5,144 | 5,235 | 4,783 | 4,316 | 4,573 | 4,427 | 4,788 | 4,638 | 4,530 | 4,469 | 4,474 | -1.12 | 0.0000807 |
| ENSMUSG00000047126 | Cltc | clathrin, heavy | protein_coding | 20,349 | 20,803 | 21,307 | 19,790 | 18,555 | 18,987 | 18,569 | 19,931 | 19,514 | 19,244 | 18,508 | 19,050 | -1.08 | 0.000664 |
| ENSMUSG00000047842 | Diras2 | DIRAS family, G | protein_coding | 2,300 | 2,213 | 2,312 | 2,008 | 1,882 | 1,928 | 1,904 | 1,951 | 2,098 | 1,970 | 1,903 | 1,815 | -1.15 | 0.000405 |
| ENSMUSG00000020321 | Mdh1 | malate dehydrc | protein_coding | 10,887 | 10,969 | 11,262 | 10,162 | 9,720 | 9,366 | 9,292 | 10,040 | 10,152 | 9,819 | 9,262 | 9,234 | -1.13 | 0.00116 |
| ENSMUSG00000024083 | Pja2 | praja ring finge | protein_coding | 18,744 | 17,965 | 18,772 | 17,517 | 16,545 | 16,702 | 16,432 | 17,205 | 17,285 | 16,602 | 16,276 | 16,385 | -1.09 | 0.0032 |
| ENSMUSG00000027419 | Pcsk2 | proprotein con | protein_coding | 7,909 | 7,743 | 8,298 | 7,477 | 7,123 | 7,230 | 6,994 | 7,341 | 7,389 | 7,325 | 6,942 | 7,090 | -1.1 | 0.00492 |
| ENSMUSG00000046598 | Bdh1 | 3-hydroxybutyr | protein_coding | 934 | 911 | 918 | 810 | 803 | 710 | 627 | 787 | 685 | 896 | 660 | 768 | -1.22 | 0.00156 |
| ENSMUSG00000040016 | Ptger3 | prostaglandin E | protein_coding | 285 | 327 | 335 | 276 | 265 | 284 | 209 | 261 | 261 | 268 | 231 | 233 | -1.19 | 0.0123 |
| ENSMUSG00000048978 | Nrsn1 | neurensin 1 | protein_coding | 7,479 | 7,253 | 7,735 | 6,789 | 6,568 | 6,599 | 5,942 | 7,029 | 6,544 | 6,746 | 6,433 | 6,279 | -1.12 | 0.00413 |
| ENSMUSG00000020402 | Vdac1 | voltage-depenc | protein_coding | 8,072 | 8,203 | 8,367 | 7,794 | 7,538 | 7,205 | 6,981 | 7,831 | 7,534 | 7,532 | 6,823 | 7,231 | -1.1 | 0.00648 |
| ENSMUSG00000078566 | Bnip3 | BCL2/adenoviru | protein_coding | 2,171 | 2,137 | 2,218 | 2,047 | 1,993 | 1,943 | 1,755 | 1,972 | 1,948 | 1,906 | 1,808 | 1,924 | -1.12 | 0.00513 |
| ENSMUSG00000027787 | Nmd3 | NMD3 ribosom | protein_coding | 1,188 | 929 | 1,073 | 1,047 | 878 | 1,002 | 858 | 757 | 912 | 902 | 906 | 750 | -1.21 | 0.0025 |
| ENSMUSG00000050786 | Ccdc126 | coiled-coil dom | protein_coding | 344 | 316 | 295 | 289 | 260 | 266 | 277 | 261 | 267 | 267 | 271 | 249 | -1.17 | 0.00992 |
| ENSMUSG00000055421 | Pcdh9 | protocadherin 9 | protein_coding | 5,485 | 4,824 | 4,802 | 4,822 | 4,156 | 4,461 | 4,199 | 4,127 | 4,361 | 4,242 | 4,326 | 4,300 | -1.18 | 0.000102 |
| ENSMUSG00000020905 | Usp43 | ubiquitin specif | protein_coding | 31 | 41 | 25 | 40 | 17 | 25 | 25 | 14 | 21 | 19 | 17 | 17 | -1.68 | 0.0132 |
| ENSMUSG00000036902 | Neto2 | neuropilin (NRF | protein_coding | 1,710 | 1,832 | 1,691 | 1,713 | 1,665 | 1,609 | 1,456 | 1,487 | 1,619 | 1,530 | 1,517 | 1,442 | -1.12 | 0.00196 |
| ENSMUSG00000035559 | Mpv17l2 | MPV17 mitocho | protein_coding | 199 | 209 | 200 | 200 | 176 | 160 | 170 | 146 | 168 | 179 | 149 | 140 | -1.24 | 0.011 |
| ENSMUSG00000055652 | Klhl25 | kelch-like 25 | protein_coding | 221 | 217 | 246 | 212 | 178 | 172 | 189 | 134 | 172 | 169 | 180 | 147 | -1.34 | 0.000881 |
| ENSMUSG00000035011 | Zbtb7a | zinc finger and | protein_coding | 4,784 | 4,571 | 4,434 | 4,461 | 4,124 | 3,964 | 4,006 | 4,226 | 3,801 | 3,863 | 4,169 | 4,323 | -1.12 | 0.00148 |
| ENSMUSG00000091264 | Smim13 | small integral m | protein_coding | 3,935 | 3,591 | 3,667 | 3,568 | 3,479 | 3,183 | 3,247 | 3,368 | 3,192 | 3,128 | 3,431 | 3,352 | -1.11 | 0.00644 |
| ENSMUSG00000037905 | Bri3bp | Bri3 binding prc | protein_coding | 3,895 | 3,846 | 3,853 | 3,767 | 3,460 | 3,391 | 3,221 | 3,523 | 3,343 | 3,408 | 3,478 | 3,524 | -1.13 | 0.000288 |
| ENSMUSG00000040907 | Atp1a3 | ATPase, Na+/K+ | protein_coding | 64,137 | 61,571 | 61,187 | 58,868 | 56,862 | 53,757 | 53,426 | 57,153 | 54,513 | 55,072 | 55,669 | 57,852 | -1.11 | 0.00123 |
| ENSMUSG00000042734 | Ttc9 | tetratricopeptic | protein_coding | 1,912 | 1,850 | 1,745 | 1,681 | 1,579 | 1,539 | 1,452 | 1,639 | 1,550 | 1,562 | 1,593 | 1,619 | -1.16 | 0.000483 |
| ENSMUSG00000038970 | Lmtk2 | lemur tyrosine | protein_coding | 3,283 | 3,352 | 3,234 | 3,175 | 2,951 | 2,938 | 2,853 | 3,043 | 2,863 | 2,973 | 3,079 | 2,911 | -1.11 | 0.000552 |
| ENSMUSG00000048481 | Mypop | Myb-related tra | protein_coding | 1,130 | 1,115 | 1,089 | 1,044 | 955 | 890 | 955 | 997 | 909 | 975 | 1,011 | 977 | -1.15 | 0.00111 |
| ENSMUSG00000033998 | Kcnk1 | potassium char | protein_coding | 1,279 | 1,173 | 1,219 | 1,118 | 1,061 | 1,056 | 935 | 1,088 | 936 | 1,058 | 1,051 | 1,049 | -1.16 | 0.00218 |
| ENSMUSG00000029875 | Ccdc184 | coiled-coil dom | protein_coding | 1,451 | 1,275 | 1,369 | 1,155 | 1,205 | 1,092 | 1,053 | 1,150 | 1,072 | 1,126 | 1,133 | 1,156 | -1.17 | 0.00101 |
| ENSMUSG00000032294 | Pkm | pyruvate kinase | protein_coding | 29,273 | 27,077 | 28,636 | 26,486 | 25,915 | 24,989 | 24,975 | 26,208 | 24,677 | 25,470 | 25,276 | 25,458 | -1.09 | 0.000494 |
| ENSMUSG00000005312 | Ubqln1 | ubiquilin 1 | protein_coding | 6,973 | 6,537 | 6,727 | 6,146 | 6,068 | 6,002 | 5,965 | 6,098 | 5,866 | 6,115 | 6,171 | 5,911 | -1.09 | 0.0066 |
| ENSMUSG00000033389 | Arhgap44 | Rho GTPase act | protein_coding | 3,469 | 3,206 | 3,292 | 3,140 | 2,930 | 2,841 | 2,843 | 2,966 | 2,866 | 2,924 | 2,998 | 2,892 | -1.13 | 0.000167 |
| ENSMUSG00000034254 | Agpat1 | 1-acylglycerol-3 | protein_coding | 2,985 | 2,817 | 2,781 | 2,488 | 2,535 | 2,394 | 2,472 | 2,591 | 2,372 | 2,528 | 2,569 | 2,601 | -1.11 | 0.00307 |
| ENSMUSG00000029120 | Ppp2r2c | protein phosph | protein_coding | 7,545 | 6,995 | 7,156 | 6,540 | 6,334 | 6,374 | 6,536 | 6,654 | 6,420 | 6,185 | 6,436 | 6,516 | -1.09 | 0.0141 |
| ENSMUSG00000051515 | Fam181b | family with seq | protein_coding | 428 | 444 | 434 | 344 | 326 | 327 | 331 | 370 | 334 | 326 | 333 | 348 | -1.22 | 0.00834 |
| ENSMUSG00000048696 | Mex3d | mex3 RNA bind | protein_coding | 567 | 493 | 504 | 534 | 428 | 394 | 452 | 401 | 361 | 394 | 492 | 403 | -1.25 | 0.000844 |
| ENSMUSG00000017314 | Mpp2 | membrane prot | protein_coding | 3,198 | 3,056 | 2,977 | 2,908 | 2,711 | 2,629 | 2,896 | 2,718 | 2,687 | 2,862 | 2,873 | 2,669 | -1.11 | 0.0002 |
| ENSMUSG00000033610 | Pank1 | pantothenate k | protein_coding | 933 | 908 | 878 | 808 | 760 | 753 | 813 | 742 | 705 | 815 | 844 | 730 | -1.15 | 0.00335 |
| ENSMUSG00000101878 | Gm8203 | predicted pseu | processed_pseudoge | 393 | 377 | 467 | 343 | 344 | 305 | 346 | 311 | 279 | 345 | 309 | 336 | -1.21 | 0.0124 |
| ENSMUSG00000020601 | Trib2 | tribbles pseudo | protein_coding | [illegible] | [illegible] | [illegible] | [illegible] | [illegible] | [illegible] | [illegible] | [illegible] | [illegible] | [illegible] | [illegible] | [illegible] | [illegible] | [illegible] |
| [illegible] | | | | | | | | | | | | | | | | | |

| | | | | |
|---|---|---|---|---|
| ENSMUSG00000031840 | Rab3a | RAB3A, membe | protein_coding | [illegible] |
| ENSMUSG00000042532 | Golga7b | golgi autoantig | protein_coding | [illegible] |
| ENSMUSG00000078994 | Zfp429 | zinc finger prot | protein_coding | [illegible] |
| ENSMUSG00000001802 | Lrp3 | low density lipo | protein_coding | [illegible] |
| ENSMUSG00000033287 | Kctd17 | potassium chan | protein_coding | [illegible] |
| ENSMUSG00000030246 | Ldhb | lactate dehydro | protein_coding | [illegible] |
| ENSMUSG00000022956 | Atp5o | ATP synthase, H | protein_coding | [illegible] |
| ENSMUSG00000063524 | Eno1 | enolase 1, alpha | protein_coding | [illegible] |
| ENSMUSG00000050043 | Tmx2 | thioredoxin-rela | protein_coding | [illegible] |
| ENSMUSG00000029076 | Sdf4 | stromal cell der | protein_coding | [illegible] |
| ENSMUSG00000037085 | Trmt12 | tRNA methyltra | protein_coding | [illegible] |
| ENSMUSG00000026249 | Serpine2 | serine (or cyste | protein_coding | [illegible] |
| ENSMUSG00000029777 | Gars | glycyl-tRNA syn | protein_coding | [illegible] |
| ENSMUSG00000030695 | Aldoa | aldolase A, fruc | protein_coding | [illegible] |
| ENSMUSG00000022295 | Atp6v1c1 | ATPase, H+ tran | protein_coding | [illegible] |
| ENSMUSG00000038486 | Sv2a | synaptic vesicle | protein_coding | [illegible] |
| ENSMUSG00000040010 | Slc7a5 | solute carrier fa | protein_coding | [illegible] |
| ENSMUSG00000019494 | Cops6 | COP9 signaloso | protein_coding | [illegible] |
| ENSMUSG00000021024 | Psma6 | proteasome sul | protein_coding | [illegible] |
| ENSMUSG00000023921 | Mmut | methylmalonyl- | protein_coding | [illegible] |
| ENSMUSG00000042670 | Immp1l | IMP1 inner mito | protein_coding | [illegible] |
| ENSMUSG00000008734 | Gprc5b | G protein-coupl | protein_coding | [illegible] |
| ENSMUSG00000020519 | Sap30l | SAP30-like | protein_coding | [illegible] |
| ENSMUSG00000027351 | Spred1 | sprouty protein | protein_coding | [illegible] |
| ENSMUSG00000050148 | Ubqln2 | ubiquilin 2 | protein_coding | [illegible] |
| ENSMUSG00000005483 | Dnajb1 | DnaJ heat shocl | protein_coding | [illegible] |
| ENSMUSG00000089762 | Ier5l | immediate earl | protein_coding | [illegible] |
| ENSMUSG00000040321 | Zfp770 | zinc finger prot | protein_coding | [illegible] |
| ENSMUSG00000045598 | Zfp553 | zinc finger prot | protein_coding | [illegible] |
| ENSMUSG00000030087 | Klf15 | Kruppel-like fac | protein_coding | [illegible] |
| ENSMUSG00000034880 | Mrpl34 | mitochondrial r | protein_coding | [illegible] |
| ENSMUSG00000042903 | Foxo4 | forkhead box O | protein_coding | [illegible] |
| ENSMUSG00000028893 | Sesn2 | sestrin 2 | protein_coding | [illegible] |
| ENSMUSG00000051495 | Irf2bp2 | interferon regu | protein_coding | [illegible] |
| ENSMUSG00000047260 | Emc6 | ER membrane p | protein_coding | [illegible] |
| ENSMUSG00000047777 | Phf13 | PHD finger prot | protein_coding | [illegible] |
| ENSMUSG00000045252 | Zfp574 | zinc finger prot | protein_coding | [illegible] |
| ENSMUSG00000060152 | Pop5 | processing of p | protein_coding | [illegible] |
| ENSMUSG00000007617 | Homer1 | homer scaffoldi | protein_coding | [illegible] |
| ENSMUSG00000070462 | Tlnrd1 | talin rod domai | protein_coding | [illegible] |
| ENSMUSG00000050428 | Fbxo46 | F-box protein 4 | protein_coding | [illegible] |
| ENSMUSG00000059486 | Kbtbd2 | kelch repeat an | protein_coding | [illegible] |
| ENSMUSG00000026696 | Vamp4 | vesicle-associat | protein_coding | [illegible] |
| ENSMUSG00000037278 | Tmem97 | transmembrane | protein_coding | [illegible] |
| ENSMUSG00000024427 | Spry4 | sprouty RTK sig | protein_coding | [illegible] |
| ENSMUSG00000027534 | Snx16 | sorting nexin 1 | protein_coding | [illegible] |
| ENSMUSG00000043298 | Smco3 | single-pass mer | protein_coding | [illegible] |
| ENSMUSG00000050144 | Slc25a44 | solute carrier fa | protein_coding | [illegible] |
| ENSMUSG00000021690 | Jmy | junction-media | protein_coding | [illegible] |
| ENSMUSG00000033717 | Adra2a | adrenergic rece | protein_coding | [illegible] |
| ENSMUSG00000040331 | Nsmce4a | NSE4 homolog | protein_coding | [illegible] |
| ENSMUSG00000032652 | Crebl2 | cAMP responsiv | protein_coding | [illegible] |
| ENSMUSG00000042763 | Maneal | mannosidase, e | protein_coding | [illegible] |
| ENSMUSG00000017631 | Abr | active BCR-rela | protein_coding | [illegible] |
| ENSMUSG00000047181 | Samd14 | sterile alpha mo | protein_coding | [illegible] |
| ENSMUSG00000027088 | Phospho2 | phosphatase, o | protein_coding | [illegible] |
| ENSMUSG00000028341 | Nr4a3 | nuclear recepto | protein_coding | [illegible] |
| ENSMUSG00000040653 | Ppp1r14c | protein phosph | protein_coding | [illegible] |
| ENSMUSG00000093789 | Methig1 | methyltransfera | protein_coding | [illegible] |
| ENSMUSG00000033319 | Fem1c | fem 1 homolog | protein_coding | [illegible] |
| ENSMUSG00000038612 | Mcl1 | myeloid cell leu | protein_coding | [illegible] |
| ENSMUSG00000022103 | Gfra2 | glial cell line de | protein_coding | [illegible] |
| ENSMUSG00000036422 | Pcdh8 | protocadherin 8 | protein_coding | [illegible] |
| ENSMUSG00000018500 | Adora2b | adenosine A2b | protein_coding | [illegible] |
| ENSMUSG00000027531 | Impa1 | inositol (myo)-1 | protein_coding | [illegible] |
| ENSMUSG00000029518 | Rab35 | RAB35, membe | protein_coding | [illegible] |
| ENSMUSG00000029064 | Gnb1 | guanine nucleo | protein_coding | [illegible] |
| ENSMUSG00000042406 | Atf4 | activating trans | protein_coding | [illegible] |
| ENSMUSG00000029708 | Gcc1 | golgi coiled coil | protein_coding | [illegible] |
| ENSMUSG00000024491 | Rbm27 | RNA binding mo | protein_coding | [illegible] |
| ENSMUSG00000031545 | Gpat4 | glycerol-3-phos | protein_coding | [illegible] |
| ENSMUSG00000071074 | Yipf3 | Yip1 domain fa | protein_coding | [illegible] |
| ENSMUSG00000002948 | Map2k7 | mitogen-activat | protein_coding | [illegible] |
| ENSMUSG00000033938 | Ndufb7 | NADH:ubiquino | protein_coding | [illegible] |
| ENSMUSG00000003813 | Rad23a | RAD23 homolog | protein_coding | [illegible] |
| ENSMUSG00000045071 | E130308A19Rik | RIKEN cDNA E1 | protein_coding | [illegible] |
| ENSMUSG00000021774 | Ube2e1 | ubiquitin-conju | protein_coding | [illegible] |
| ENSMUSG00000022951 | Rcan1 | regulator of cal | protein_coding | [illegible] |
| ENSMUSG00000023175 | Bsg | basigin | protein_coding | [illegible] |
| ENSMUSG00000046058 | Eid2 | EP300 interacti | protein_coding | [illegible] |
| ENSMUSG00000022247 | Brix1 | BRX1, biogenes | protein_coding | [illegible] |
| ENSMUSG00000039058 | Ak5 | adenylate kinas | protein_coding | [illegible] |
| ENSMUSG00000004069 | Dnaja3 | DnaJ heat shocl | protein_coding | [illegible] |
| ENSMUSG00000017721 | Pigt | phosphatidylino | protein_coding | [illegible] |
| ENSMUSG00000036199 | Ndufa13 | NADH:ubiquino | protein_coding | [illegible] |
| ENSMUSG00000016349 | Eef1a2 | eukaryotic tran | protein_coding | [illegible] |
| ENSMUSG00000020483 | Dynll2 | dynein light cha | protein_coding | [illegible] |
| ENSMUSG00000019173 | Rab5c | RAB5C, membe | protein_coding | [illegible] |
| ENSMUSG00000017390 | Aldoc | aldolase C, fruc | protein_coding | [illegible] |
| ENSMUSG00000025208 | Mrpl43 | mitochondrial r | protein_coding | [illegible] |
| ENSMUSG00000053838 | Nudcd3 | NudC domain c | protein_coding | [illegible] |
| ENSMUSG00000047379 | B4gat1 | beta-1,4-glucur | protein_coding | [illegible] |
| ENSMUSG00000053253 | Ndfip2 | Nedd4 family in | protein_coding | [illegible] |
| ENSMUSG00000034190 | Chmp7 | charged multiv | protein_coding | [illegible] |
| ENSMUSG00000029146 | Snx17 | sorting nexin 17 | protein_coding | [illegible] |
| ENSMUSG00000035765 | Dym | dymeclin | protein_coding | [illegible] |
| ENSMUSG00000028517 | Plpp3 | phospholipid pl | protein_coding | [illegible] |
| ENSMUSG00000008035 | Mid1ip1 | Mid1 interactin | protein_coding | [illegible] |
| ENSMUSG00000033917 | Gde1 | glycerophospho | protein_coding | [illegible] |
| ENSMUSG00000047187 | Rab2a | RAB2A, membe | protein_coding | [illegible] |
| ENSMUSG00000025016 | Tm9sf3 | transmembrane | protein_coding | [illegible] |
| ENSMUSG00000013593 | Ndufs2 | NADH:ubiquino | protein_coding | [illegible] |
| ENSMUSG00000075702 | Selenom | selenoprotein M | protein_coding | [illegible] |
| ENSMUSG00000027170 | Eif3m | eukaryotic tran | protein_coding | [illegible] |
| ENSMUSG00000029198 | Grpel1 | GrpE-like 1, mit | protein_coding | [illegible] |
| ENSMUSG00000037242 | Clic4 | chloride intrace | protein_coding | [illegible] |
| ENSMUSG00000020986 | Sec23a | SEC23 homolog | protein_coding | [illegible] |
| ENSMUSG00000032097 | Ddx6 | DEAD (Asp-Glu- | protein_coding | [illegible] |
| ENSMUSG00000016382 | Pls3 | plastin 3 (T-isof | protein_coding | [illegible] |
| ENSMUSG00000022018 | Rgcc | regulator of cel | protein_coding | [illegible] |
| ENSMUSG00000003948 | Mmd | monocyte to m | protein_coding | [illegible] |
| ENSMUSG00000037843 | Vstm2l | V-set and trans | protein_coding | [illegible] |
| ENSMUSG00000045136 | Tubb2b | tubulin, beta 2B | protein_coding | [illegible] |
| ENSMUSG00000014313 | Cox6c | cytochrome c o | protein_coding | [illegible] |
| ENSMUSG00000116165 | Pdxp | pyridoxal (pyrid | protein_coding | [illegible] |
| ENSMUSG00000028445 | Enho | energy homeos | protein_coding | [illegible] |
| ENSMUSG00000019210 | Atp6v1e1 | ATPase, H+ tran | protein_coding | [illegible] |
| ENSMUSG00000025204 | Ndufb8 | NADH:ubiquino | protein_coding | [illegible] |
| ENSMUSG00000031633 | Slc25a4 | solute carrier fa | protein_coding | [illegible] |
| ENSMUSG00000006315 | Tmem147 | transmembrane | protein_coding | [illegible] |
| ENSMUSG00000054793 | Cadm4 | cell adhesion m | protein_coding | [illegible] |
| ENSMUSG00000031818 | Cox4i1 | cytochrome c o | protein_coding | [illegible] |
| ENSMUSG00000034994 | Eef2 | eukaryotic tran | protein_coding | [illegible] |
| ENSMUSG00000058254 | Tspan7 | tetraspanin 7 | protein_coding | [illegible] |
| ENSMUSG00000067925 | Rtl8a | retrotransposo | protein_coding | [illegible] |
| ENSMUSG00000020708 | Psmc5 | protease (prosc | protein_coding | [illegible] |
| ENSMUSG00000051391 | Ywhag | tyrosine 3-mon | protein_coding | [illegible] |
| ENSMUSG00000026932 | Nacc2 | nucleus accumb | protein_coding | [illegible] |
| ENSMUSG00000068923 | Syt11 | synaptotagmin | protein_coding | [illegible] |
| ENSMUSG00000025428 | Atp5a1 | ATP synthase, H | protein_coding | [illegible] |
| ENSMUSG00000059995 | Atxn7l3 | ataxin 7-like 3 | protein_coding | [illegible] |
| ENSMUSG00000025781 | Atp5c1 | ATP synthase, H | protein_coding | [illegible] |
| ENSMUSG00000047710 | Champ1 | chromosome al | protein_coding | [illegible] |
| ENSMUSG00000017707 | Serinc3 | serine incorpor | protein_coding | [illegible] |
| ENSMUSG00000027195 | Hsd17b12 | hydroxysteroid | protein_coding | [illegible] |
| ENSMUSG00000009863 | Sdhb | succinate dehy | protein_coding | [illegible] |
| ENSMUSG00000031703 | Itfg1 | integrin alpha F | protein_coding | [illegible] |
| ENSMUSG00000115987 | Vps28 | vacuolar protei | protein_coding | [illegible] |
| ENSMUSG00000000420 | Galnt1 | polypeptide N-a | protein_coding | [illegible] |
| ENSMUSG00000001017 | Chtop | chromatin targe | protein_coding | [illegible] |
| ENSMUSG00000028779 | Pef1 | penta-EF hand | protein_coding | [illegible] |
| ENSMUSG00000027546 | Atp9a | ATPase, class II, | protein_coding | [illegible] |
| ENSMUSG00000034189 | Hsdl1 | hydroxysteroid | protein_coding | [illegible] |
| ENSMUSG00000036275 | 9530068E07Rik | RIKEN cDNA 95 | protein_coding | [illegible] |
| ENSMUSG00000040385 | Ppp1ca | protein phosph | protein_coding | [illegible] |
| ENSMUSG00000003346 | Abhd17a | abhydrolase do | protein_coding | [illegible] |
| ENSMUSG00000048758 | Rpl29 | ribosomal prote | protein_coding | [illegible] |
| ENSMUSG00000028757 | Ddost | dolichyl-di-pho | protein_coding | [illegible] |
| ENSMUSG00000029632 | Ndufa4 | Ndufa4, mitoch | protein_coding | [illegible] |
| ENSMUSG00000032046 | Abhd12 | abhydrolase do | protein_coding | [illegible] |
| ENSMUSG00000037419 | Endod1 | endonuclease d | protein_coding | [illegible] |
| ENSMUSG00000004267 | Eno2 | enolase 2, gam | protein_coding | [illegible] |
| ENSMUSG00000024099 | Ndufv2 | NADH:ubiquino | protein_coding | [illegible] |
| ENSMUSG00000031451 | Gas6 | growth arrest s | protein_coding | [illegible] |
| ENSMUSG00000048307 | Ankrd46 | ankyrin repeat | protein_coding | [illegible] |
| ENSMUSG00000017776 | Crk | v-crk avian sarc | protein_coding | [illegible] |
| ENSMUSG00000039740 | Alg2 | asparagine-link | protein_coding | [illegible] |
| ENSMUSG00000038888 | Ctu1 | cytosolic thiour | protein_coding | [illegible] |
| ENSMUSG00000011752 | Pgam1 | phosphoglycera | protein_coding | [illegible] |
| ENSMUSG00000051065 | Mb21d2 | Mab-21 domain | protein_coding | [illegible] |
| ENSMUSG00000058672 | Tubb2a | tubulin, beta 2A | protein_coding | [illegible] |
| ENSMUSG00000024091 | Vapa | vesicle-associat | protein_coding | [illegible] |
| ENSMUSG00000032320 | Rcn2 | reticulocalbin 2 | protein_coding | [illegible] |
| ENSMUSG00000035964 | Tmem59l | transmembrane | protein_coding | [illegible] |
| ENSMUSG00000057614 | Gnai1 | guanine nucleo | protein_coding | [illegible] |
| ENSMUSG00000018326 | Ywhab | tyrosine 3-mon | protein_coding | [illegible] |
| ENSMUSG00000021087 | Rtn1 | reticulon 1 | protein_coding | [illegible] |
| ENSMUSG00000021877 | Arf4 | ADP-ribosylatio | protein_coding | [illegible] |
| ENSMUSG00000032399 | Rpl4 | ribosomal prote | protein_coding | [illegible] |
| ENSMUSG00000029030 | Tprgl | transformation | protein_coding | [illegible] |
| ENSMUSG00000042364 | Snx18 | sorting nexin 18 | protein_coding | [illegible] |
| ENSMUSG00000027637 | Rab5if | RAB5 interactin | protein_coding | [illegible] |
| ENSMUSG00000021496 | Pcbd2 | pterin 4 alpha c | protein_coding | [illegible] |
| ENSMUSG00000035274 | Tpbg | trophoblast gly | protein_coding | [illegible] |
| ENSMUSG00000033735 | Spr | sepiapterin red | protein_coding | [illegible] |
| ENSMUSG00000040945 | Rcc2 | regulator of chr | protein_coding | [illegible] |
| ENSMUSG00000015981 | Stk32c | serine/threonin | protein_coding | [illegible] |
| ENSMUSG00000030588 | Yif1b | Yip1 interacting | protein_coding | [illegible] |
| ENSMUSG00000049086 | Bmyc | brain expressed | protein_coding | [illegible] |
| ENSMUSG00000027223 | Mapk8ip1 | mitogen-activat | protein_coding | [illegible] |
| ENSMUSG00000028436 | Dcaf12 | DDB1 and CUL4 | protein_coding | [illegible] |
| ENSMUSG00000031778 | Cx3cl1 | chemokine (C-X | protein_coding | [illegible] |
| ENSMUSG00000069227 | Gprin1 | G protein-regul | protein_coding | [illegible] |
| ENSMUSG00000059734 | Ndufs8 | NADH:ubiquino | protein_coding | [illegible] |
| ENSMUSG00000020149 | Rab1a | RAB1A, membe | protein_coding | [illegible] |
| ENSMUSG00000047514 | Tspyl1 | testis-specific p | protein_coding | [illegible] |
| ENSMUSG00000026112 | Coa5 | cytochrome C c | protein_coding | [illegible] |
| ENSMUSG00000028081 | Rps3a1 | ribosomal prote | protein_coding | [illegible] |
| ENSMUSG00000031059 | Ndufb11 | NADH:ubiquino | protein_coding | [illegible] |
| ENSMUSG00000005873 | Reep5 | receptor access | protein_coding | [illegible] |
| ENSMUSG00000020612 | Prkar1a | protein kinase, | protein_coding | [illegible] |
| ENSMUSG00000031534 | Smim19 | small integral m | protein_coding | [illegible] |
| ENSMUSG00000007892 | Rplp1 | ribosomal prote | protein_coding | [illegible] |
| ENSMUSG00000023010 | Tmbim6 | transmembrane | protein_coding | [illegible] |
| ENSMUSG00000056429 | Tgoln1 | trans-golgi netv | protein_coding | [illegible] |
| ENSMUSG00000019054 | Fis1 | fission, mitocho | protein_coding | [illegible] |
| ENSMUSG00000074649 | BC029722 | cDNA sequence | protein_coding | [illegible] |
| ENSMUSG00000032437 | Stt3b | STT3, subunit o | protein_coding | [illegible] |
| ENSMUSG00000048076 | Arf1 | ADP-ribosylatio | protein_coding | [illegible] |
| ENSMUSG00000001783 | Rtcb | RNA 2',3'-cyclic | protein_coding | [illegible] |
| ENSMUSG00000060636 | Rpl35a | ribosomal prote | protein_coding | [illegible] |
| ENSMUSG00000031827 | Cotl1 | coactosin-like 1 | protein_coding | [illegible] |
| ENSMUSG00000041355 | Ssr2 | signal sequence | protein_coding | [illegible] |
| ENSMUSG00000057880 | Abat | 4-aminobutyrat | protein_coding | [illegible] |
| ENSMUSG00000018677 | Slc25a39 | solute carrier fa | protein_coding | [illegible] |
| ENSMUSG00000025132 | Arhgdia | Rho GDP dissoc | protein_coding | [illegible] |
| ENSMUSG00000020349 | Ppp2ca | protein phosph | protein_coding | [illegible] |
| ENSMUSG00000022982 | Sod1 | superoxide dism | protein_coding | [illegible] |
| ENSMUSG00000006932 | Ctnnb1 | catenin (cadher | protein_coding | [illegible] |
| ENSMUSG00000005354 | Txn2 | thioredoxin 2 | protein_coding | [illegible] |
| ENSMUSG00000020132 | Rab21 | RAB21, membe | protein_coding | [illegible] |
| ENSMUSG00000031823 | Zdhhc7 | zinc finger, DHH | protein_coding | [illegible] |
| ENSMUSG00000020267 | Hint1 | histidine triad n | protein_coding | [illegible] |
| ENSMUSG00000020978 | Klhdc2 | kelch domain co | protein_coding | [illegible] |
| ENSMUSG00000021314 | Amph | amphiphysin | protein_coding | [illegible] |
| ENSMUSG00000009630 | Ppp2cb | protein phosph | protein_coding | [illegible] |
| ENSMUSG00000056579 | Tug1 | taurine upregul | protein_coding | [illegible] |
| ENSMUSG00000028152 | Tspan5 | tetraspanin 5 | protein_coding | [illegible] |
| ENSMUSG00000075232 | Amd1 | S-adenosylmeth | protein_coding | [illegible] |
| ENSMUSG00000041740 | Rnf10 | ring finger prot | protein_coding | [illegible] |
| ENSMUSG00000044792 | Isca1 | iron-sulfur clust | protein_coding | [illegible] |
| ENSMUSG00000022351 | Sqle | squalene epoxi | protein_coding | [illegible] |
| ENSMUSG00000026229 | Psmd1 | proteasome (pr | protein_coding | [illegible] |
| ENSMUSG00000002844 | Adprh | ADP-ribosylargi | protein_coding | [illegible] |
| ENSMUSG00000054364 | Rhob | ras homolog fa | protein_coding | [illegible] |
| ENSMUSG00000014606 | Slc25a11 | solute carrier fa | protein_coding | [illegible] |
| ENSMUSG00000034353 | Ramp1 | receptor (calcit | protein_coding | [illegible] |
| ENSMUSG00000026750 | Psmb7 | proteasome (pr | protein_coding | [illegible] |
| ENSMUSG00000068749 | Psma5 | proteasome sul | protein_coding | [illegible] |
| ENSMUSG00000029059 | Prxl2b | peroxiredoxin li | protein_coding | [illegible] |
| ENSMUSG00000035530 | Eif1 | eukaryotic tran | protein_coding | [illegible] |
| ENSMUSG00000030662 | Ipo5 | importin 5 | protein_coding | [illegible] |
| ENSMUSG00000044533 | Rps2 | ribosomal prote | protein_coding | [illegible] |
| ENSMUSG00000060036 | Rpl3 | ribosomal prote | protein_coding | [illegible] |
| ENSMUSG00000028452 | Vcp | valosin containi | protein_coding | [illegible] |
| ENSMUSG00000058569 | Tmed9 | transmembrane | protein_coding | [illegible] |
| ENSMUSG00000003970 | Rpl8 | ribosomal prote | protein_coding | [illegible] |
| ENSMUSG00000024425 | Ndfip1 | Nedd4 family in | protein_coding | [illegible] |
| ENSMUSG00000015776 | Med22 | mediator comp | protein_coding | [illegible] |
| ENSMUSG00000045912 | C2cd4c | C2 calcium-dep | protein_coding | [illegible] |

| | | | | | | | | | | | | | | | | |
|---|---|---|---|---|---|---|---|---|---|---|---|---|---|---|---|---|
| ENSMUSG00000051232 | Tmem199 | transmembran | protein_coding | 802 | 883 | 859 | 771 | 730 | 723 | 859 | 813 | 740 | 764 | 645 | 708 | -1.06 |
| ENSMUSG00000106037 | Gm4332 | predicted gene | processed_pseudoge | 691 | 845 | 774 | 605 | 552 | 495 | 705 | 702 | 575 | 634 | 430 | 459 | -1.19 |
| ENSMUSG00000001525 | Tubb5 | tubulin, beta 5 | protein_coding | 12,506 | 12,174 | 12,640 | 10,792 | 10,525 | 10,448 | 11,960 | 11,024 | 10,967 | 10,829 | 10,098 | 9,919 | -1.09 |
| ENSMUSG00000008036 | Ap2s1 | adaptor-relatec | protein_coding | 1,898 | 1,788 | 1,973 | 1,527 | 1,549 | 1,462 | 1,748 | 1,609 | 1,597 | 1,598 | 1,411 | 1,366 | -1.13 |
| ENSMUSG00000028692 | Akr1a1 | aldo-keto reduc | protein_coding | 5,971 | 5,782 | 6,277 | 5,243 | 5,027 | 5,067 | 5,894 | 5,220 | 5,298 | 5,363 | 4,788 | 4,771 | -1.1 |
| ENSMUSG00000093674 | Rpl41 | ribosomal prot | protein_coding | 5,806 | 5,677 | 6,131 | 4,721 | 4,750 | 4,538 | 5,642 | 4,821 | 5,013 | 5,077 | 4,249 | 4,043 | -1.13 |
| ENSMUSG00000067274 | Rplp0 | ribosomal prot | protein_coding | 3,866 | 3,737 | 3,904 | 3,294 | 3,282 | 3,331 | 3,827 | 3,588 | 3,339 | 3,454 | 3,117 | 3,113 | -1.05 |
| ENSMUSG00000039105 | Atp6v1g1 | ATPase, H+ trar | protein_coding | 1,985 | 1,963 | 2,107 | 1,693 | 1,750 | 1,620 | 1,976 | 1,783 | 1,718 | 1,805 | 1,584 | 1,578 | -1.09 |
| ENSMUSG00000039530 | | | | 3,347 | 3,355 | 3,705 | 2,853 | 2,929 | 2,695 | 3,464 | 3,102 | 2,924 | 3,019 | 2,467 | 2,478 | -1.09 |
| ENSMUSG00000032959 | Pebp1 | phosphatidyletl | protein_coding | 6,252 | 6,435 | 6,469 | 5,113 | 5,205 | 5,052 | 5,707 | 5,463 | 5,344 | 5,351 | 4,517 | 4,471 | -1.13 |
| ENSMUSG00000006273 | Atp6v1b2 | ATPase, H+ trar | protein_coding | 13,167 | 13,620 | 13,456 | 11,781 | 11,336 | 11,126 | 12,345 | 11,986 | 11,953 | 11,813 | 10,636 | 10,490 | -1.11 |
| ENSMUSG00000029621 | Arpc1a | actin related pr | protein_coding | 2,750 | 2,867 | 2,878 | 2,464 | 2,355 | 2,331 | 2,634 | 2,499 | 2,542 | 2,519 | 2,151 | 2,141 | -1.12 |
| ENSMUSG00000062933 | Gm10123 | predicted pseu | processed_pseudoge | 6,464 | 6,886 | 7,149 | 5,693 | 5,655 | 5,704 | 6,240 | 6,445 | 5,932 | 5,916 | 5,007 | 5,023 | -1.09 |
| ENSMUSG00000076437 | Selenoh | selenoprotein H | protein_coding | 320 | 359 | 367 | 284 | 272 | 265 | 305 | 310 | 295 | 290 | 222 | 242 | -1.15 |
| ENSMUSG00000035078 | Mtmr9 | myotubularin r | protein_coding | 2,144 | 2,331 | 2,334 | 1,980 | 2,018 | 1,840 | 2,134 | 2,065 | 2,052 | 1,990 | 1,813 | 1,749 | -1.09 |
| ENSMUSG00000040952 | Rps19 | ribosomal prot | protein_coding | 2,791 | 3,195 | 3,087 | 2,371 | 2,459 | 2,186 | 2,944 | 2,489 | 2,496 | 2,426 | 2,168 | 2,055 | -1.14 |
| ENSMUSG00000025870 | Arl10 | ADP-ribosylatio | protein_coding | 925 | 959 | 1,002 | 749 | 799 | 723 | 897 | 766 | 788 | 810 | 673 | 683 | -1.14 |
| ENSMUSG00000028691 | Prdx1 | peroxiredoxin 1 | protein_coding | 3,124 | 3,297 | 3,335 | 2,761 | 2,733 | 2,556 | 3,071 | 2,699 | 2,811 | 2,842 | 2,380 | 2,237 | -1.13 |
| ENSMUSG00000025351 | Cd63 | CD63 antigen | protein_coding | 1,532 | 1,589 | 1,576 | 1,249 | 1,367 | 1,322 | 1,543 | 1,430 | 1,385 | 1,364 | 1,173 | 1,141 | -1.05 |
| ENSMUSG00000069972 | Rps13-ps2 | ribosomal prot | processed_pseudoge | 1,255 | 1,316 | 1,435 | 1,013 | 996 | 1,023 | 1,360 | 1,063 | 1,132 | 1,067 | 922 | 777 | -1.13 |
| ENSMUSG00000027963 | Extl2 | exostosin-like g | protein_coding | 2,195 | 2,087 | 2,286 | 1,958 | 1,927 | 2,130 | 2,129 | 1,888 | 1,889 | 1,914 | 1,683 | 1,855 | -1.06 |
| ENSMUSG00000058655 | Eif4b | eukaryotic tran | protein_coding | 8,743 | 8,296 | 8,909 | 8,167 | 8,268 | 8,422 | 8,430 | 8,086 | 8,235 | 8,089 | 7,633 | 7,886 | -1.03 |
| ENSMUSG00000015932 | Dstn | destrin | protein_coding | 3,879 | 3,529 | 3,882 | 3,425 | 3,405 | 3,499 | 3,776 | 3,533 | 3,469 | 3,336 | 3,188 | 3,251 | -1.04 |
| ENSMUSG00000025967 | Eef1b2 | eukaryotic tran | protein_coding | 3,094 | 2,816 | 3,089 | 2,745 | 2,794 | 2,632 | 2,992 | 2,857 | 2,681 | 2,746 | 2,527 | 2,551 | -1.04 |
| ENSMUSG00000022108 | Itm2b | integral membr | protein_coding | 17,043 | 15,713 | 17,144 | 15,429 | 15,204 | 15,199 | 16,597 | 15,652 | 14,658 | 14,864 | 14,042 | 14,109 | -1.04 |
| ENSMUSG00000030224 | Strap | serine/threonir | protein_coding | 3,199 | 3,023 | 3,241 | 2,909 | 2,697 | 2,733 | 3,112 | 2,891 | 2,694 | 2,722 | 2,523 | 2,569 | -1.08 |
| ENSMUSG00000018446 | C1qbp | complement co | protein_coding | 1,225 | 1,181 | 1,199 | 1,081 | 1,089 | 1,114 | 1,127 | 1,081 | 1,071 | 1,083 | 987 | 983 | -1.06 |
| ENSMUSG00000039501 | Znfx1 | zinc finger, NFX | protein_coding | 1,773 | 1,677 | 1,764 | 1,522 | 1,440 | 1,578 | 1,609 | 1,536 | 1,532 | 1,436 | 1,404 | 1,312 | -1.09 |
| ENSMUSG00000007836 | Hnrnpa0 | heterogeneous | protein_coding | 3,688 | 3,591 | 3,962 | 3,282 | 3,262 | 3,260 | 3,458 | 3,225 | 3,221 | 3,310 | 2,852 | 2,908 | -1.1 |
| ENSMUSG00000021610 | Clptm1l | CLPTM1-like | protein_coding | 2,086 | 2,045 | 2,294 | 1,816 | 1,829 | 1,928 | 1,972 | 1,846 | 1,864 | 1,963 | 1,596 | 1,628 | -1.09 |
| ENSMUSG00000022661 | Cd200 | CD200 molecul | protein_coding | 5,344 | 5,328 | 5,699 | 4,687 | 4,746 | 4,802 | 5,333 | 5,144 | 4,933 | 4,860 | 4,300 | 4,358 | -1.05 |
| ENSMUSG00000006333 | Rps9 | ribosomal prot | protein_coding | 5,420 | 5,362 | 5,770 | 4,840 | 4,806 | 4,866 | 5,293 | 4,951 | 4,846 | 4,798 | 4,393 | 4,268 | -1.07 |
| ENSMUSG00000009927 | Rps25 | ribosomal prot | protein_coding | 4,379 | 4,245 | 4,639 | 3,735 | 3,810 | 3,772 | 4,210 | 3,948 | 3,848 | 3,888 | 3,487 | 3,423 | -1.08 |
| ENSMUSG00000022336 | Eif3e | eukaryotic tran | protein_coding | 2,126 | 2,025 | 2,126 | 1,832 | 1,856 | 1,823 | 2,025 | 1,882 | 1,899 | 1,835 | 1,688 | 1,729 | -1.07 |
| ENSMUSG00000057841 | Rpl32 | ribosomal prot | protein_coding | 3,544 | 3,375 | 3,548 | 3,003 | 3,115 | 3,044 | 3,386 | 3,145 | 3,121 | 3,160 | 2,760 | 2,787 | -1.06 |
| ENSMUSG00000051234 | Rnf7 | ring finger prot | protein_coding | 1,633 | 1,586 | 1,662 | 1,449 | 1,417 | 1,458 | 1,563 | 1,446 | 1,440 | 1,452 | 1,335 | 1,355 | -1.08 |
| ENSMUSG00000061111 | Mcrip1 | MAPK regulate | protein_coding | 1,099 | 1,001 | 1,124 | 927 | 921 | 938 | 1,009 | 924 | 943 | 938 | 822 | 836 | -1.09 |
| ENSMUSG00000021114 | Atp6v1d | ATPase, H+ trar | protein_coding | 4,485 | 4,452 | 4,796 | 4,243 | 4,218 | 4,034 | 4,531 | 4,294 | 4,019 | 4,277 | 3,794 | 3,943 | -1.05 |
| ENSMUSG00000047675 | Rps8 | ribosomal prot | protein_coding | 6,408 | 6,390 | 7,073 | 6,059 | 6,073 | 5,605 | 6,376 | 6,042 | 5,739 | 5,916 | 4,969 | 5,241 | -1.08 |
| ENSMUSG00000056665 | Them6 | thioesterase su | protein_coding | 552 | 555 | 601 | 501 | 487 | 479 | 528 | 499 | 476 | 497 | 419 | 480 | -1.11 |
| ENSMUSG00000024150 | Mcfd2 | multiple coagul | protein_coding | 2,387 | 2,384 | 2,520 | 2,204 | 2,177 | 2,116 | 2,307 | 2,184 | 2,084 | 2,182 | 1,969 | 2,048 | -1.08 |
| ENSMUSG00000078812 | Eif5a | eukaryotic tran | protein_coding | 4,813 | 4,881 | 5,199 | 4,425 | 4,364 | 4,085 | 4,585 | 4,312 | 4,180 | 4,337 | 3,845 | 4,093 | -1.11 |
| ENSMUSG00000021660 | Btf3 | basic transcript | protein_coding | 2,562 | 2,590 | 2,735 | 2,347 | 2,417 | 2,416 | 2,557 | 2,353 | 2,317 | 2,397 | 2,160 | 2,286 | -1.05 |
| ENSMUSG00000030754 | Copb1 | coatomer prote | protein_coding | 2,054 | 1,992 | 2,118 | 1,901 | 1,942 | 1,923 | 2,015 | 1,912 | 1,874 | 1,856 | 1,773 | 1,790 | -1.04 |
| ENSMUSG00000031447 | Lamp1 | lysosomal-asso | protein_coding | 10,726 | 10,516 | 11,104 | 9,568 | 10,011 | 9,667 | 10,788 | 9,647 | 9,662 | 9,483 | 8,861 | 9,234 | -1.04 |
| ENSMUSG00000036155 | Mgat5 | mannoside ace | protein_coding | 4,135 | 4,144 | 4,419 | 4,058 | 4,020 | 3,859 | 4,139 | 3,803 | 3,964 | 3,920 | 3,747 | 3,798 | -1.06 |
| ENSMUSG00000027828 | Ssr3 | signal sequence | protein_coding | 4,828 | 4,580 | 5,229 | 4,357 | 4,504 | 4,361 | 4,682 | 4,325 | 4,435 | 4,315 | 4,070 | 4,137 | -1.06 |
| ENSMUSG00000100755 | Rps23-ps1 | ribosomal prot | processed_pseudoge | 3,686 | 3,478 | 4,198 | 3,403 | 3,477 | 3,234 | 3,682 | 3,263 | 3,331 | 3,235 | 2,888 | 3,120 | -1.08 |
| ENSMUSG00000030772 | Dkk3 | dickkopf WNT s | protein_coding | 2,186 | 2,195 | 2,342 | 2,138 | 2,105 | 2,110 | 2,250 | 2,027 | 2,089 | 1,957 | 1,871 | 1,899 | -1.04 |
| ENSMUSG00000025544 | Tm9sf2 | transmembran | protein_coding | 4,161 | 4,255 | 4,445 | 4,072 | 4,087 | 3,886 | 4,314 | 3,928 | 3,930 | 3,884 | 3,586 | 3,464 | -1.04 |
| ENSMUSG00000028138 | Adh5 | alcohol dehydr | protein_coding | 1,054 | 1,032 | 1,102 | 975 | 975 | 920 | 1,049 | 974 | 961 | 905 | 785 | 827 | -1.06 |
| ENSMUSG00000060743 | H3f3a | H3.3 histone A | protein_coding | 2,851 | 2,816 | 3,089 | 2,530 | 2,618 | 2,562 | 2,949 | 2,564 | 2,561 | 2,506 | 2,054 | 2,185 | -1.06 |
| ENSMUSG00000067870 | Rpl31-ps8 | ribosomal prot | processed_pseudoge | 2,693 | 2,725 | 3,026 | 2,587 | 2,556 | 2,503 | 2,769 | 2,627 | 2,398 | 2,460 | 2,387 | 2,277 | -1.05 |
| ENSMUSG00000078974 | Sec61g | SEC61, gamma | protein_coding | 194 | 217 | 258 | 183 | 190 | 165 | 234 | 189 | 171 | 174 | 155 | 152 | -1.09 |
| ENSMUSG00000025579 | Gaa | glucosidase, alp | protein_coding | 31,611 | 31,772 | 33,170 | 30,840 | 29,231 | 30,138 | 32,744 | 30,311 | 29,473 | 29,796 | 28,479 | 29,031 | -1.04 |
| ENSMUSG00000006373 | Pgrmc1 | progesterone r | protein_coding | 12,519 | 12,204 | 13,188 | 11,472 | 10,971 | 11,205 | 13,012 | 11,422 | 11,278 | 11,351 | 10,222 | 10,438 | -1.06 |
| ENSMUSG00000037742 | Eef1a1 | eukaryotic tran | protein_coding | 67,929 | 67,442 | 72,242 | 62,743 | 62,467 | 62,803 | 70,801 | 65,820 | 62,658 | 62,623 | 56,731 | 57,925 | -1.03 |
| ENSMUSG00000058927 | Gm10053 | predicted gene | protein_coding | 1,704 | 1,765 | 1,880 | 1,606 | 1,553 | 1,606 | 1,653 | 1,821 | 1,553 | 1,488 | 1,471 | 1,536 | -1.05 |
| ENSMUSG00000024683 | Mrpl16 | mitochondrial r | protein_coding | 425 | 444 | 456 | 381 | 370 | 359 | 411 | 408 | 379 | 363 | 329 | 391 | -1.1 |
| ENSMUSG00000069682 | Gm10275 | predicted pseu | transcribed_processe | 3,765 | 3,649 | 3,971 | 3,271 | 3,232 | 3,261 | 3,514 | 3,558 | 3,215 | 3,261 | 2,972 | 3,161 | -1.08 |
| ENSMUSG00000032324 | Tspan3 | tetraspanin 3 | protein_coding | 6,650 | 6,302 | 7,045 | 6,000 | 5,918 | 6,032 | 6,196 | 6,101 | 5,936 | 5,743 | 5,477 | 5,871 | -1.07 |
| ENSMUSG00000049036 | Tmem121 | transmembran | protein_coding | 284 | 257 | 290 | 215 | 205 | 219 | 242 | 222 | 214 | 205 | 187 | 202 | -1.18 |
| ENSMUSG00000022421 | Nptxr | neuronal pentr | protein_coding | 10,012 | 9,934 | 10,015 | 9,181 | 9,016 | 8,980 | 9,807 | 9,385 | 8,820 | 9,159 | 8,810 | 9,153 | -1.05 |
| ENSMUSG00000003316 | Glg1 | golgi apparatus | protein_coding | 9,870 | 9,979 | 10,201 | 9,332 | 9,088 | 9,102 | 9,694 | 9,609 | 8,798 | 8,968 | 8,993 | 9,045 | -1.05 |
| ENSMUSG00000031016 | Wee1 | WEE 1 homolog | protein_coding | 540 | 572 | 584 | 486 | 489 | 476 | 564 | 516 | 458 | 473 | 477 | 482 | -1.07 |
| ENSMUSG00000060126 | Tpt1 | tumor protein, | protein_coding | 7,385 | 7,064 | 7,917 | 6,059 | 6,378 | 6,265 | 7,432 | 7,247 | 5,940 | 6,501 | 5,716 | 5,931 | -1.04 |
| ENSMUSG00000068391 | Chrac1 | chromatin acce | protein_coding | 258 | 253 | 257 | 226 | 217 | 215 | 261 | 237 | 191 | 215 | 183 | 203 | -1.07 |
| ENSMUSG00000020038 | Cry1 | cryptochrome 1 | protein_coding | 592 | 629 | 669 | 547 | 465 | 543 | 571 | 566 | 514 | 552 | 490 | 502 | -1.13 |
| ENSMUSG00000020520 | Galnt10 | polypeptide N- | protein_coding | 864 | 848 | 892 | 740 | 687 | 794 | 800 | 816 | 745 | 761 | 703 | 736 | -1.08 |
| ENSMUSG00000035666 | Gtf3c4 | general transcri | protein_coding | 2,048 | 2,087 | 2,099 | 1,860 | 1,769 | 1,877 | 2,036 | 1,949 | 1,830 | 1,873 | 1,771 | 1,809 | -1.06 |
| ENSMUSG00000042628 | Zfyve1 | zinc finger, FYV | protein_coding | 1,360 | 1,376 | 1,460 | 1,084 | 948 | 1,136 | 1,380 | 1,108 | 1,024 | 1,048 | 906 | 1,020 | -1.16 |
| ENSMUSG00000044030 | Irf2bp1 | interferon regu | protein_coding | 1,386 | 1,385 | 1,414 | 1,163 | 1,173 | 1,196 | 1,343 | 1,214 | 1,147 | 1,175 | 1,108 | 1,145 | -1.09 |
| ENSMUSG00000071657 | Bscl2 | Berardinelli-Sei | protein_coding | 2,181 | 2,226 | 2,162 | 1,803 | 1,841 | 1,915 | 2,062 | 1,945 | 1,838 | 1,900 | 1,805 | 1,775 | -1.08 |
| ENSMUSG00000028964 | Park7 | Parkinson disea | protein_coding | 1,924 | 2,002 | 2,129 | 1,890 | 1,743 | 1,752 | 1,914 | 1,829 | 1,785 | 1,794 | 1,728 | 1,724 | -1.1 |
| ENSMUSG00000031007 | Atp6ap2 | ATPase, H+ trar | protein_coding | 5,068 | 5,055 | 5,365 | 4,861 | 4,543 | 4,457 | 4,853 | 4,771 | 4,620 | 4,663 | 4,457 | 4,434 | -1.09 |
| ENSMUSG00000022454 | Nell2 | NEL-like 2 | protein_coding | 5,400 | 5,358 | 5,606 | 5,096 | 4,668 | 4,704 | 4,937 | 4,927 | 4,944 | 4,732 | 4,633 | 4,463 | -1.12 |
| ENSMUSG00000031227 | Magee1 | MAGE family m | protein_coding | 7,066 | 7,194 | 7,076 | 6,768 | 6,351 | 6,290 | 6,801 | 6,641 | 6,482 | 6,377 | 6,256 | 6,100 | -1.08 |
| ENSMUSG00000002957 | Ap2a2 | adaptor-relatec | protein_coding | 5,858 | 5,937 | 5,750 | 5,276 | 5,075 | 4,874 | 5,458 | 5,342 | 5,214 | 5,185 | 4,930 | 4,992 | -1.1 |
| ENSMUSG00000021957 | Tkt | transketolase | protein_coding | 5,961 | 5,861 | 5,820 | 5,311 | 5,209 | 4,963 | 5,606 | 5,484 | 5,275 | 5,216 | 4,837 | 5,078 | -1.08 |
| ENSMUSG00000040652 | Oaz2 | ornithine decar | protein_coding | 4,323 | 4,266 | 4,526 | 3,887 | 3,683 | 3,577 | 4,095 | 4,028 | 3,797 | 3,891 | 3,575 | 3,570 | -1.1 |
| ENSMUSG00000047921 | Trappc9 | trafficking prot | protein_coding | 2,238 | 2,174 | 2,294 | 2,019 | 1,828 | 1,819 | 2,103 | 2,038 | 1,945 | 1,900 | 1,824 | 1,740 | -1.12 |
| ENSMUSG00000006299 | Aamp | angio-associate | protein_coding | 3,268 | 3,231 | 3,277 | 2,971 | 2,828 | 2,814 | 3,250 | 3,044 | 2,861 | 2,893 | 2,803 | 2,765 | -1.07 |
| ENSMUSG00000007777 | 0610009B22Rik | RIKEN cDNA 06 | protein_coding | 392 | 396 | 387 | 329 | 310 | 292 | 374 | 346 | 314 | 332 | 302 | 278 | -1.13 |
| ENSMUSG00000052833 | Sae1 | SUMO1 activati | protein_coding | 1,871 | 2,003 | 1,888 | 1,595 | 1,532 | 1,337 | 1,724 | 1,729 | 1,594 | 1,490 | 1,583 | 1,416 | -1.16 |
| ENSMUSG00000001416 | Cct3 | chaperonin con | protein_coding | 3,167 | 3,362 | 3,374 | 2,766 | 2,930 | 2,697 | 3,145 | 2,975 | 2,883 | 2,778 | 2,781 | 2,724 | -1.08 |
| ENSMUSG00000022426 | Josd1 | Josephin domai | protein_coding | 935 | 1,012 | 1,008 | 868 | 856 | 785 | 961 | 901 | 828 | 823 | 834 | 816 | -1.09 |
| ENSMUSG00000043881 | Kbtbd7 | kelch repeat an | protein_coding | 1,525 | 1,601 | 1,668 | 1,417 | 1,379 | 1,322 | 1,503 | 1,491 | 1,404 | 1,344 | 1,383 | 1,324 | -1.09 |
| ENSMUSG00000023147 | Get1 | guided entry of | protein_coding | 2,154 | 2,200 | 2,292 | 1,946 | 1,890 | 1,825 | 2,070 | 1,986 | 1,890 | 1,895 | 1,844 | 1,712 | -1.1 |
| ENSMUSG00000044894 | Uqcrq | ubiquinol-cytoc | protein_coding | 1,228 | 1,325 | 1,337 | 1,067 | 1,023 | 965 | 1,104 | 1,084 | 1,019 | 1,033 | 979 | 890 | -1.19 |
| ENSMUSG00000007950 | Abhd8 | abhydrolase do | protein_coding | 4,080 | 4,326 | 4,513 | 3,382 | 3,509 | 3,408 | 3,802 | 3,600 | 3,470 | 3,488 | 3,301 | 3,082 | -1.14 |
| ENSMUSG00000070493 | Chchd2 | coiled-coil-helix | protein_coding | 3,347 | 3,406 | 3,616 | 2,704 | 2,736 | 2,530 | 2,983 | 2,798 | 2,836 | 2,746 | 2,601 | 2,375 | -1.18 |
| ENSMUSG00000041881 | Ndufa7 | NADH:ubiquino | protein_coding | 987 | 1,007 | 1,004 | 778 | 789 | 724 | 878 | 843 | 817 | 799 | 725 | 659 | -1.17 |
| ENSMUSG00000026123 | Plekhb2 | pleckstrin hom | protein_coding | 9,847 | 9,590 | 9,760 | 8,415 | 8,385 | 8,082 | 8,839 | 8,677 | 8,492 | 8,436 | 7,890 | 7,659 | -1.11 |
| ENSMUSG00000033629 | Hacd3 | 3-hydroxyacyl-C | protein_coding | 6,227 | 6,103 | 6,273 | 5,610 | 5,622 | 5,398 | 5,891 | 5,744 | 5,613 | 5,616 | 5,397 | 5,225 | -1.07 |
| ENSMUSG00000030654 | Arl6ip1 | ADP-ribosylatio | protein_coding | 6,380 | 6,491 | 6,901 | 5,577 | 5,721 | 5,249 | 6,050 | 5,971 | 5,642 | 5,656 | 5,183 | 5,044 | -1.1 |
| ENSMUSG00000074748 | Atxn7l3b | ataxin 7-like 3B | protein_coding | 4,581 | 4,651 | 4,850 | 4,149 | 4,112 | 3,792 | 4,341 | 4,267 | 4,083 | 3,990 | 3,830 | 3,814 | -1.1 |
| ENSMUSG00000007739 | Cct4 | chaperonin con | protein_coding | 3,358 | 3,318 | 3,358 | 2,813 | 2,964 | 2,651 | 3,188 | 2,957 | 2,818 | 2,844 | 2,867 | 2,658 | -1.09 |
| ENSMUSG00000051695 | Pcbp1 | poly(rC) binding | protein_coding | 4,529 | 4,307 | 4,563 | 3,863 | 3,919 | 3,664 | 4,391 | 3,934 | 3,796 | 3,904 | 3,876 | 3,626 | -1.09 |
| ENSMUSG00000025791 | Pgm1 | phosphoglucon | protein_coding | 3,157 | 3,222 | 3,258 | 2,866 | 2,737 | 2,746 | 3,063 | 2,875 | 2,842 | 2,866 | 2,856 | 2,534 | -1.09 |
| ENSMUSG00000030007 | Cct7 | chaperonin con | protein_coding | 4,695 | 4,640 | 4,714 | 4,225 | 3,977 | 3,847 | 4,455 | 4,058 | 4,002 | 3,966 | 4,031 | 3,722 | -1.12 |
| ENSMUSG00000017978 | Cadps2 | Ca2+-depender | protein_coding | 2,799 | 2,977 | 2,994 | 2,676 | 2,666 | 2,416 | 3,006 | 2,588 | 2,507 | 2,621 | 2,516 | 2,382 | -1.07 |
| ENSMUSG00000061983 | Rps12 | ribosomal prot | protein_coding | 2,123 | 2,159 | 2,280 | 1,811 | 1,865 | 1,738 | 2,232 | 1,802 | 1,758 | 1,851 | 1,763 | 1,725 | -1.1 |
| ENSMUSG00000035585 | Tsen34 | tRNA splicing e | protein_coding | 1,438 | 1,438 | 1,528 | 1,286 | 1,326 | 1,222 | 1,418 | 1,247 | 1,229 | 1,285 | 1,218 | 1,182 | -1.09 |
| ENSMUSG00000051615 | Rap2a | RAS related pr | protein_coding | 2,833 | 2,767 | 3,030 | 2,503 | 2,515 | 2,391 | 2,794 | 2,516 | 2,408 | 2,463 | 2,302 | 2,349 | -1.09 |
| ENSMUSG00000001663 | Gstt1 | glutathione S-tr | protein_coding | 393 | 376 | 380 | 289 | 348 | 305 | 412 | 299 | 287 | 326 | 269 | 254 | -1.06 |
| ENSMUSG00000060419 | Rps16-ps2 | ribosomal prot | processed_pseudoge | 1,836 | 1,670 | 1,887 | 1,497 | 1,452 | 1,577 | 1,908 | 1,485 | 1,411 | 1,617 | 1,386 | 1,174 | -1.07 |
| ENSMUSG00000067578 | Cbln4 | cerebellin 4 pre | protein_coding | 1,030 | 1,005 | 1,046 | 958 | 902 | 903 | 1,096 | 898 | 962 | 842 | 826 | 835 | -1.06 |
| ENSMUSG00000006651 | Aplp1 | amyloid beta (A | protein_coding | 24,061 | 22,033 | 24,186 | 20,928 | 20,371 | 20,294 | 23,154 | 20,361 | 20,814 | 20,753 | 19,677 | 19,565 | -1.08 |
| ENSMUSG00000007458 | M6pr | mannose-6-pho | protein_coding | 3,463 | 3,330 | 3,501 | 3,154 | 3,107 | 3,147 | 3,398 | 3,120 | 3,119 | 3,102 | 3,013 | 2,968 | -1.05 |
| ENSMUSG00000034300 | Fam53c | family with seq | protein_coding | 1,585 | 1,580 | 1,645 | 1,435 | 1,378 | 1,379 | 1,642 | 1,355 | 1,419 | 1,365 | 1,350 | 1,331 | -1.09 |
| ENSMUSG00000117310 | Ptp4a1 | protein tyrosin | protein_coding | 3,406 | 3,221 | 3,495 | 3,057 | 3,039 | 2,899 | 3,364 | 2,960 | 3,023 | 2,913 | 2,955 | 2,908 | -1.07 |
| ENSMUSG00000041773 | Enc1 | ectodermal-neu | protein_coding | 5,094 | 5,147 | 5,166 | 4,670 | 4,187 | 4,487 | 5,223 | 4,629 | 4,428 | 4,293 | 4,712 | 4,248 | -1.08 |
| ENSMUSG00000005803 | Sqor | sulfide quinone | protein_coding | 267 | 227 | 269 | 208 | 163 | 179 | 244 | 178 | 179 | 167 | 188 | 134 | -1.28 |
| ENSMUSG00000021770 | Samd8 | sterile alpha m | protein_coding | 2,431 | 2,303 | 2,410 | 2,294 | 2,048 | 2,079 | 2,314 | 2,111 | 2,120 | 1,966 | 2,017 | 1,985 | -1.1 |
| ENSMUSG00000024231 | Cul2 | cullin 2 | protein_coding | 2,082 | 2,128 | 2,280 | 2,269 | 2,044 | 2,084 | 2,166 | 2,247 | 2,010 | 2,042 | 1,884 | 1,922 | -1.02 |
| ENSMUSG00000022526 | Zfp251 | zinc finger prot | protein_coding | 1,081 | 1,044 | 1,214 | 1,124 | 1,019 | 1,053 | 1,043 | 1,062 | 951 | 1,032 | 968 | 966 | -1.07 |
| ENSMUSG00000050628 | Ubald2 | UBA-like domai | protein_coding | 289 | 287 | 361 | 319 | 285 | 250 | 267 | 290 | 236 | 267 | 239 | 250 | -1.14 |
| ENSMUSG00000028646 | Rragc | Ras-related GTI | protein_coding | 2,080 | 2,213 | 2,270 | 2,189 | 1,862 | 1,753 | 2,197 | 2,015 | 1,908 | 1,931 | 1,809 | 1,892 | -1.12 |
| ENSMUSG00000024742 | Fen1 | flap structure s | protein_coding | 247 | 247 | 283 | 247 | 210 | 181 | 241 | 235 | 218 | 197 | 171 | 202 | -1.17 |
| ENSMUSG00000031897 | Psmb10 | proteasome (pr | protein_coding | 498 | 485 | 521 | 519 | 428 | 380 | 503 | 460 | 434 | 432 | 360 | 406 | -1.14 |
| ENSMUSG00000037253 | Mex3c | mex3 RNA bind | protein_coding | 819 | 846 | 968 | 807 | 860 | 731 | 853 | 822 | 796 | 702 | 749 | 720 | -1.05 |
| ENSMUSG00000038622 | Med30 | mediator comp | protein_coding | 365 | 378 | 464 | 400 | 379 | 341 | 406 | 395 | 355 | 344 | 322 | 336 | -1.05 |
| ENSMUSG00000040133 | Gpr176 | G protein-coup | protein_coding | 576 | 561 | 682 | 598 | 536 | 482 | 557 | 562 | 556 | 524 | 484 | 462 | -1.13 |
| ENSMUSG00000036885 | Arhgef26 | Rho guanine nu | protein_coding | 1,296 | 1,478 | 1,679 | 1,092 | 1,143 | 1,222 | 1,250 | 1,204 | 1,238 | 1,148 | 1,029 | 1,114 | -1.15 |
| ENSMUSG00000073755 | 5730409E04Rik | RIKEN cDNA 57 | protein_coding | 1,708 | 1,753 | 1,894 | 1,552 | 1,520 | 1,561 | 1,552 | 1,580 | 1,548 | 1,518 | 1,456 | 1,545 | -1.11 |
| ENSMUSG00000024151 | Msh2 | mutS homolog | protein_coding | 1,180 | 1,244 | 1,386 | 1,175 | 1,130 | 1,125 | 1,155 | 1,180 | 1,107 | 1,135 | 1,106 | 1,101 | -1.08 |
| ENSMUSG00000045328 | Cenpe | centromere pro | protein_coding | 22 | 56 | 80 | 19 | 23 | 20 | 17 | 28 | 16 | 16 | 13 | 22 | -1.79 |
| ENSMUSG00000050029 | Rap2c | RAP2C, membe | protein_coding | 969 | 994 | 1,023 | 911 | 900 | 880 | 906 | 846 | 900 | 859 | 892 | 796 | -1.1 |
| ENSMUSG00000028675 | Pnrc2 | proline-rich nuc | protein_coding | 1,401 | 1,576 | 1,821 | 1,288 | 1,266 | 1,258 | 1,141 | 1,211 | 1,155 | 1,054 | 1,205 | 1,104 | -1.25 |
| ENSMUSG00000036949 | Slc39a12 | solute carrier f | protein_coding | 2,042 | 1,953 | 2,245 | 1,929 | 1,815 | 1,828 | 1,806 | 1,750 | 1,845 | 1,696 | 1,813 | 1,686 | -1.13 |
| ENSMUSG00000020646 | Mboat2 | membrane bou | protein_coding | 2,443 | 2,612 | 2,630 | 2,389 | 2,405 | 2,358 | 2,300 | 2,206 | 2,375 | 2,304 | 2,114 | 2,205 | -1.09 |
| ENSMUSG00000002741 | Ykt6 | YKT6 v-SNARE I | protein_coding | 1,548 | 1,620 | 1,723 | 1,529 | 1,467 | 1,432 | 1,472 | 1,488 | 1,483 | 1,370 | 1,382 | 1,463 | -1.09 |
| ENSMUSG00000024601 | Isoc1 | isochorismatas | protein_coding | 1,326 | 1,416 | 1,408 | 1,303 | 1,243 | 1,179 | 1,178 | 1,231 | 1,200 | 1,189 | 1,162 | 1,165 | -1.13 |
| ENSMUSG00000032281 | Acsbg1 | acyl-CoA synth | protein_coding | 4,282 | 4,507 | 4,809 | 3,982 | 3,908 | 3,841 | 3,775 | 3,879 | 3,896 | 3,807 | 3,707 | 3,581 | -1.14 |
| ENSMUSG00000000339 | Rtca | RNA 3'-termina | protein_coding | 824 | 824 | 867 | 798 | 758 | 776 | 759 | 820 | 802 | 738 | 623 | 705 | -1.06 |
| ENSMUSG00000034501 | Pcnx4 | pecanex homol | protein_coding | 1,526 | 1,577 | 1,626 | 1,499 | 1,391 | 1,484 | 1,422 | 1,513 | 1,438 | 1,426 | 1,303 | 1,344 | -1.07 |
| ENSMUSG00000032564 | Cpne4 | copine IV | protein_coding | 829 | 952 | 1,042 | 915 | 701 | 898 | 791 | 765 | 786 | 786 | 655 | 697 | -1.19 |
| ENSMUSG00000028945 | Rheb | Ras homolog er | protein_coding | 2,115 | 2,187 | 2,277 | 2,113 | 2,013 | 2,069 | 1,959 | 1,982 | 2,096 | 1,930 | 1,895 | 1,862 | -1.08 |
| ENSMUSG00000001891 | Ugp2 | UDP-glucose py | protein_coding | 1,966 | 1,969 | 2,162 | 1,941 | 1,758 | 1,868 | 1,795 | 1,855 | 1,874 | 1,751 | 1,641 | 1,572 | -1.1 |
| ENSMUSG00000051359 | Ncald | neurocalcin del | protein_coding | 10,656 | 10,183 | 11,332 | 10,250 | 9,490 | 9,766 | 9,625 | 9,869 | 10,018 | 9,649 | 9,136 | 9,148 | -1.09 |
| ENSMUSG00000005802 | Slc30a4 | solute carrier f | protein_coding | 1,297 | 1,410 | 1,491 | 1,369 | 1,242 | 1,266 | 1,240 | 1,260 | 1,295 | 1,340 | 1,126 | 1,142 | -1.11 |
| ENSMUSG00000032932 | Hspa13 | heat shock prot | protein_coding | 2,472 | 2,689 | 2,681 | 2,450 | 2,302 | 2,265 | 2,398 | 2,384 | 2,315 | 2,480 | 2,110 | 2,122 | -1.1 |
| ENSMUSG00000058600 | Rpl30 | ribosomal prot | protein_coding | 2,449 | 2,604 | 2,864 | 2,408 | 2,399 | 2,289 | 2,418 | 2,424 | 2,293 | 2,384 | 2,131 | 2,036 | -1.08 |
| ENSMUSG00000035478 | Mbd3 | methyl-CpG bin | protein_coding | 1,701 | 1,848 | 2,043 | 1,671 | 1,612 | 1,520 | 1,707 | 1,633 | 1,595 | 1,600 | 1,512 | 1,482 | -1.12 |
| ENSMUSG00000016940 | Kctd2 | potassium char | protein_coding | 1,235 | 1,388 | 1,402 | 1,169 | 1,164 | 1,055 | 1,195 | 1,109 | 1,121 | 1,113 | 1,029 | 1,011 | -1.15 |
| ENSMUSG00000048807 | Slc35e4 | solute carrier f | protein_coding | 538 | 585 | 633 | 535 | 537 | 468 | 538 | 496 | 497 | 503 | 452 | 429 | -1.12 |
| ENSMUSG00000000751 | Rpa1 | replication prot | protein_coding | 1,597 | 1,574 | 1,683 | 1,404 | 1,470 | 1,531 | 1,407 | 1,479 | 1,459 | 1,392 | 1,325 | 1,242 | -1.06 |
| ENSMUSG00000024208 | Uqcc2 | ubiquinol-cytoc | protein_coding | 954 | 900 | 940 | 736 | 746 | 788 | 745 | 787 | 802 | 737 | 712 | 648 | -1.15 |
| ENSMUSG00000054843 | Atrnl1 | attractin like 1 | protein_coding | 2,011 | 2,030 | 2,177 | 1,812 | 1,739 | 1,918 | 1,735 | 1,788 | 1,907 | 1,754 | 1,700 | 1,583 | -1.12 |
| ENSMUSG00000025139 | Tollip | toll interacting | protein_coding | 4,065 | 4,075 | 4,366 | 3,575 | 3,528 | 3,518 | 3,858 | 3,788 | 3,684 | 3,469 | 3,390 | 3,415 | -1.09 |
| ENSMUSG00000031021 | Tmem9b | TMEM9 domain | protein_coding | 2,003 | 1,943 | 2,057 | 1,719 | 1,709 | 1,714 | 1,833 | 1,806 | 1,847 | 1,666 | 1,650 | 1,568 | -1.09 |
| ENSMUSG00000027698 | Nceh1 | neutral cholest | protein_coding | 3,685 | 3,626 | 3,827 | 3,431 | 3,274 | 3,374 | 3,448 | 3,404 | 3,443 | 3,319 | 3,093 | 3,152 | -1.08 |
| ENSMUSG00000036766 | Dner | delta/notch-lik | protein_coding | 8,114 | 8,185 | 8,745 | 7,313 | 6,866 | 7,247 | 7,453 | 7,497 | 7,343 | 7,219 | 6,513 | 6,328 | -1.11 |
| ENSMUSG00000020152 | Actr2 | ARP2 actin-rela | protein_coding | 5,678 | 5,657 | 6,112 | 5,140 | 4,984 | 5,126 | 5,195 | 5,242 | 5,414 | 5,096 | 4,861 | 4,855 | -1.1 |
| ENSMUSG00000029126 | Nsg1 | neuron specific | protein_coding | 9,100 | 9,222 | 9,750 | 8,008 | 7,658 | 7,762 | 8,188 | 8,194 | 8,031 | 7,922 | 7,437 | 7,256 | -1.13 |
| ENSMUSG00000063358 | Mapk1 | mitogen-activa | protein_coding | 8,891 | 8,943 | 9,301 | 8,365 | 7,578 | 7,770 | 8,761 | 8,011 | 7,830 | 8,196 | 7,355 | 7,035 | -1.11 |
| ENSMUSG00000058603 | Rpl28-ps1 | ribosomal prot | processed_pseudoge | 1,779 | 1,761 | 2,061 | 1,610 | 1,506 | 1,487 | 1,716 | 1,628 | 1,581 | 1,603 | 1,418 | 1,238 | -1.14 |
| ENSMUSG00000031696 | Vps35 | VPS35 retrome | protein_coding | 5,075 | 5,039 | 5,344 | 4,653 | 4,558 | 4,560 | 4,889 | 4,747 | 4,606 | 4,629 | 4,446 | 4,198 | -1.07 |
| ENSMUSG00000035152 | Ap2b1 | adaptor-relatec | protein_coding | 8,579 | 8,656 | 8,745 | 8,115 | 7,952 | 7,929 | 8,342 | 8,202 | 8,117 | 7,986 | 7,878 | 7,482 | -1.05 |
| ENSMUSG00000056296 | Synpr | synaptoporin | protein_coding | 1,539 | 1,544 | 1,741 | 1,363 | 1,096 | 1,302 | 1,366 | 1,309 | 1,431 | 1,340 | 1,150 | 994 | -1.22 |
| ENSMUSG00000031029 | Eif3f | eukaryotic tran | protein_coding | 2,738 | 3,038 | 3,314 | 2,456 | 2,306 | 2,565 | 2,629 | 2,561 | 2,612 | 2,541 | 2,218 | 2,164 | -1.15 |
| ENSMUSG00000022010 | Tsc22d1 | TSC22 domain f | protein_coding | 7,088 | 7,513 | 7,782 | 6,631 | 6,518 | 6,698 | 6,973 | 6,761 | 6,860 | 6,555 | 6,133 | 5,887 | -1.08 |
| ENSMUSG00000047215 | Rpl9 | ribosomal prot | protein_coding | 6,099 | 6,620 | 7,126 | 5,658 | 5,247 | 5,416 | 6,228 | 5,800 | 5,788 | 5,656 | 4,764 | 4,570 | -1.12 |
| ENSMUSG00000000693 | Loxl3 | lysyl oxidase-lik | protein_coding | 122 | 224 | 135 | 155 | 128 | 131 | 137 | 127 | 118 | 106 | 119 | 123 | -1.19 |
| ENSMUSG00000114147 | Gm18939 | predicted gene | transcribed_processe | 0 | 112 | 5 | 42 | 1 | 3 | 1 | 8 | 1 | 0 | 0 | 0 | -12.2 |
| ENSMUSG00000028889 | Yrdc | yrdC domain co | protein_coding | 345 | 391 | 363 | 425 | 294 | 371 | 347 | 343 | 278 | 308 | 280 | 340 | -1.12 |
| ENSMUSG00000054836 | Elp6 | elongator acety | protein_coding | 257 | 307 | 263 | 269 | 168 | 236 | 250 | 254 | 212 | 203 | 190 | 212 | -1.2 |
| ENSMUSG00000053025 | Sv2b | synaptic vesicle | protein_coding | 2,298 | 2,412 | 2,463 | 2,212 | 1,977 | 2,247 | 2,292 | 2,084 | 2,168 | 2,044 | 2,149 | 1,931 | -1.09 |
| ENSMUSG00000056153 | Socs6 | suppressor of c | protein_coding | 788 | 867 | 886 | 751 | 701 | 790 | 796 | 704 | 748 | 701 | 667 | 642 | -1.1 |
| ENSMUSG00000060780 | Lrrtm1 | leucine rich rep | protein_coding | 808 | 916 | 884 | 854 | 742 | 722 | 850 | 802 | 744 | 730 | 755 | 680 | -1.11 |
| ENSMUSG00000023169 | Slc38a1 | solute carrier f | protein_coding | 11,886 | 12,463 | 12,325 | 11,720 | 11,089 | 11,485 | 12,038 | 11,663 | 11,683 | 11,324 | 11,584 | 10,955 | -1.05 |
| ENSMUSG00000039262 | Prrc2b | proline-rich coil | protein_coding | 19,481 | 20,385 | 20,470 | 19,745 | 18,856 | 19,182 | 20,092 | 19,546 | 19,505 | 19,041 | 18,960 | 18,309 | -1.03 |
| ENSMUSG00000026999 | Nup35 | nucleoporin 35 | protein_coding | 363 | 361 | 353 | 316 | 274 | 357 | 335 | 346 | 299 | 273 | 281 | 264 | -1.06 |
| ENSMUSG00000012405 | Rpl15 | ribosomal prot | protein_coding | 2,903 | 3,221 | 3,085 | 2,923 | 2,652 | 3,017 | 2,972 | 3,054 | 2,664 | 2,580 | 2,270 | 2,500 | -1.04 |
| ENSMUSG00000026374 | Tsn | translin | protein_coding | 2,875 | 2,900 | 2,916 | 2,745 | 2,474 | 2,783 | 2,913 | 2,740 | 2,641 | 2,582 | 2,265 | 2,404 | -1.05 |
| ENSMUSG00000024074 | Crim1 | cysteine rich tr | protein_coding | 3,662 | 3,865 | 3,681 | 3,392 | 3,272 | 3,163 | 3,590 | 3,348 | 3,318 | 2,996 | 3,144 | 3,118 | -1.09 |
| ENSMUSG00000031974 | Abcb10 | ATP-binding ca | protein_coding | 1,070 | 1,119 | 1,061 | 971 | 957 | 986 | 1,070 | 1,035 | 948 | 914 | 881 | 896 | -1.04 |
| ENSMUSG00000068732 | Tmem167b | transmembran | protein_coding | 1,576 | 1,636 | 1,697 | 1,482 | 1,398 | 1,384 | 1,675 | 1,577 | 1,481 | 1,361 | 1,339 | 1,289 | -1.06 |
| ENSMUSG00000002416 | Ndufb2 | NADH:ubiquino | protein_coding | 1,303 | 1,351 | 1,334 | 1,250 | 1,094 | 1,121 | 1,196 | 1,269 | 1,132 | 1,154 | 1,054 | 1,007 | -1.12 |
| ENSMUSG00000086171 | Pcsk2os1 | proprotein con | lncRNA | 168 | 207 | 160 | 155 | 118 | 135 | 162 | 158 | 122 | 126 | 95 | 84 | -1.21 |
| ENSMUSG00000029247 | Paics | phosphoribosyl | protein_coding | 2,511 | 2,820 | 2,601 | 2,491 | 2,464 | 2,411 | 2,594 | 2,461 | 2,343 | 2,332 | 2,289 | 2,159 | -1.05 |
| ENSMUSG00000042682 | Selenok | selenoprotein K | protein_coding | 1,938 | 2,017 | 2,033 | 1,929 | 1,875 | 1,941 | 1,962 | 1,948 | 1,852 | 1,834 | 1,797 | 1,754 | -1.02 |
| ENSMUSG00000044502 | Bod1 | biorientation o | protein_coding | 1,342 | 1,386 | 1,387 | 1,284 | 1,247 | 1,233 | 1,310 | 1,273 | 1,176 | 1,175 | 1,111 | 1,057 | -1.07 |
| ENSMUSG00000026988 | Wdsub1 | WD repeat, SAI | protein_coding | 432 | 504 | 465 | 365 | 433 | 389 | 402 | 351 | 344 | 334 | 316 | 379 | -1.12 |
| ENSMUSG00000032519 | Slc25a38 | solute carrier f | protein_coding | 439 | 519 | 478 | 395 | 441 | 406 | 422 | 405 | 375 | 351 | 406 | 406 | -1.09 |
| ENSMUSG00000021670 | Hmgcr | 3-hydroxy-3-m | protein_coding | 2,970 | 3,502 | 2,961 | 2,575 | 2,487 | 2,253 | 3,010 | 2,745 | 2,444 | 2,549 | 2,328 | 2,839 | -1.14 |
| ENSMUSG00000022947 | Cbr3 | carbonyl reduc | protein_coding | 91 | 108 | 90 | 72 | 55 | 71 | 79 | 77 | 65 | 63 | 54 | 67 | -1.28 |
| ENSMUSG00000024030 | Abcg1 | ATP binding cas | protein_coding | 2,649 | 2,794 | 2,523 | 2,276 | 2,050 | 2,156 | 2,364 | 2,177 | 2,172 | 2,149 | 1,979 | 2,116 | -1.17 |
| ENSMUSG00000038807 | Rap1gap2 | RAP1 GTPase a | protein_coding | 4,304 | 4,509 | 4,333 | 4,086 | 3,836 | 3,775 | 4,108 | 3,925 | 3,883 | 3,737 | 3,662 | 3,830 | -1.1 |
| ENSMUSG00000094708 | Gm10359 | predicted gene | processed_pseudoge | 0 | 4,588 | 878 | 0 | 0 | 0 | 1,077 | 0 | 7 | 0 | 0 | 0 | -5.07 |
| ENSMUSG00000095937 | Gm12671 | predicted gene | processed_pseudoge | 0 | 4,588 | 878 | 0 | 0 | 0 | 1,077 | 0 | 7 | 0 | 0 | 0 | -5.07 |
| ENSMUSG00000079562 | Maea | macrophage er | protein_coding | 1,975 | 2,182 | 2,235 | 2,022 | 1,908 | 1,948 | 2,077 | 2,046 | 1,820 | 1,965 | 1,810 | 1,908 | -1.05 |
| ENSMUSG00000042787 | Exog | endo/exonucle | protein_coding | 493 | 623 | 604 | 503 | 420 | 458 | 538 | 475 | 446 | 471 | 460 | 432 | -1.18 |
| ENSMUSG00000061517 | Sox21 | SRY (sex detern | protein_coding | 275 | 332 | 325 | 265 | 233 | 221 | 284 | 267 | 222 | 246 | 222 | 224 | -1.19 |
| ENSMUSG00000027601 | Mtfr1 | mitochondrial f | protein_coding | 317 | 368 | 323 | 288 | 297 | 265 | 297 | 293 | 244 | 287 | 272 | 249 | -1.12 |
| ENSMUSG00000036957 | Lrfn3 | leucine rich rep | protein_coding | 560 | 596 | 563 | 491 | 477 | 464 | 536 | 491 | 462 | 480 | 444 | 442 | -1.12 |
| ENSMUSG00000048965 | Mrgpre | MAS-related GI | protein_coding | 985 | 1,061 | 1,019 | 915 | 902 | 870 | 979 | 860 | 839 | 881 | 814 | 838 | -1.1 |
| ENSMUSG00000031980 | Agt | angiotensinoge | protein_coding | 5,730 | 6,122 | 5,935 | 5,418 | 5,375 | 5,234 | 5,808 | 5,294 | 5,191 | 5,523 | 5,086 | 4,799 | -1.07 |
| ENSMUSG00000032594 | Ip6k1 | inositol hexaph | protein_coding | 7,182 | 8,264 | 7,649 | 6,954 | 7,298 | 6,682 | 7,589 | 6,418 | 6,371 | 6,969 | 6,118 | 6,058 | -1.07 |
| ENSMUSG00000034432 | Cops8 | COP9 signaloso | protein_coding | 1,864 | 2,110 | 1,761 | 1,752 | 1,626 | 1,632 | 1,460 | 1,699 | 1,726 | 1,769 | 1,375 | 1,286 | -1.17 |
| ENSMUSG00000027316 | Gfra4 | glial cell line de | protein_coding | 470 | 505 | 427 | 448 | 406 | 431 | 385 | 410 | 412 | 409 | 379 | 383 | -1.14 |
| ENSMUSG00000051343 | Rab11fip5 | RAB11 family ir | protein_coding | 3,094 | 3,140 | 2,649 | 2,674 | 2,528 | 2,567 | 2,357 | 2,689 | 2,514 | 2,586 | 2,392 | 2,399 | -1.14 |
| ENSMUSG00000020346 | Mgat1 | mannoside ace | protein_coding | 1,308 | 1,429 | 1,379 | 1,234 | 1,204 | 1,170 | 1,231 | 1,169 | 1,245 | 1,208 | 1,090 | 1,037 | -1.12 |
| ENSMUSG00000026667 | Uhmk1 | U2AF homolog | protein_coding | 12,857 | 13,454 | 13,662 | 12,135 | 11,884 | 11,554 | 12,225 | 12,057 | 12,348 | 12,001 | 11,551 | 10,916 | -1.09 |
| ENSMUSG00000049680 | Urgcp | upregulator of | protein_coding | 2,150 | 2,313 | 2,232 | 1,878 | 1,887 | 1,830 | 1,930 | 1,920 | 1,820 | 1,984 | 1,791 | 1,624 | -1.13 |
| ENSMUSG00000001270 | Ckb | creatine kinase | protein_coding | 17,445 | 17,871 | 18,166 | 14,826 | 15,529 | 15,316 | 15,352 | 15,409 | 15,844 | 15,642 | 14,134 | 13,343 | -1.11 |
| ENSMUSG00000020271 | Fbxw11 | F-box and WD- | protein_coding | 4,114 | 4,127 | 4,236 | 3,734 | 3,700 | 3,616 | 3,789 | 3,726 | 3,668 | 3,785 | 3,423 | 3,316 | -1.09 |
| ENSMUSG00000025580 | Eif4a3 | eukaryotic tran | protein_coding | 1,066 | 1,139 | 1,064 | 948 | 909 | 967 | 961 | 943 | 939 | 927 | 911 | 834 | -1.11 |
| ENSMUSG00000030105 | Arl8b | ADP-ribosylatio | protein_coding | 6,079 | 6,185 | 5,660 | 5,177 | 5,354 | 5,202 | 5,415 | 5,238 | 5,402 | 5,152 | 4,798 | 4,636 | -1.09 |
| ENSMUSG00000038462 | Uqcrfs1 | ubiquinol-cytoc | protein_coding | 1,885 | 2,025 | 1,863 | 1,564 | 1,589 | 1,561 | 1,650 | 1,734 | 1,671 | 1,621 | 1,475 | 1,358 | -1.12 |
| ENSMUSG00000024896 | Minpp1 | multiple inosit | protein_coding | 1,568 | 1,588 | 1,488 | 1,402 | 1,393 | 1,324 | 1,489 | 1,409 | 1,321 | 1,391 | 1,329 | 1,181 | -1.08 |
| ENSMUSG00000017188 | Coa3 | cytochrome C o | protein_coding | 1,024 | 987 | 955 | 793 | 830 | 829 | 888 | 830 | 815 | 858 | 780 | 703 | -1.11 |
| ENSMUSG00000049686 | Orai1 | ORAI calcium re | protein_coding | 184 | 189 | 156 | 131 | 147 | 129 | 152 | 140 | 124 | 140 | 124 | 115 | -1.16 |
| ENSMUSG00000046722 | Cdc42se1 | CDC42 small eff | protein_coding | 1,067 | 1,111 | 1,039 | 947 | 953 | 896 | 1,047 | 993 | 944 | 986 | 864 | 846 | -1.07 |
| ENSMUSG00000000171 | Sdhd | succinate dehy | protein_coding | 1,383 | 1,599 | 1,358 | 1,131 | 1,119 | 1,037 | 1,339 | 1,216 | 1,198 | 1,191 | 1,051 | 993 | -1.16 |
| ENSMUSG00000029581 | Fscn1 | fascin actin-bur | protein_coding | 2,611 | 2,666 | 2,510 | 2,187 | 2,173 | 2,106 | 2,404 | 2,199 | 2,224 | 2,252 | 2,047 | 1,972 | -1.12 |
| ENSMUSG00000005510 | Ndufs3 | NADH:ubiquino | protein_coding | 1,353 | 1,556 | 1,425 | 1,245 | 1,213 | 1,178 | 1,338 | 1,319 | 1,207 | 1,279 | 1,154 | 1,105 | -1.1 |
| ENSMUSG00000055839 | Elob | elongin B | protein_coding | 1,623 | 1,722 | 1,613 | 1,343 | 1,364 | 1,397 | 1,489 | 1,446 | 1,360 | 1,457 | 1,225 | 1,248 | -1.11 |
| ENSMUSG00000002803 | Btbd6 | BTB (POZ) dom | protein_coding | 908 | 911 | 851 | 784 | 873 | 846 | 821 | 838 | 799 | 774 | 756 | 716 | -1.02 |
| ENSMUSG00000037190 | Cyb561d2 | cytochrome b-5 | protein_coding | 196 | 206 | 177 | 153 | 170 | 155 | 160 | 194 | 162 | 148 | 132 | 123 | -1.07 |
| ENSMUSG00000014550 | Rbsn | rabenosyn, RAE | protein_coding | 2,040 | 2,085 | 2,025 | 1,902 | 1,981 | 1,796 | 1,971 | 1,807 | 1,922 | 1,789 | 1,755 | 1,717 | -1.07 |
| ENSMUSG00000028156 | Eif4e | eukaryotic tran | protein_coding | 3,711 | 3,851 | 3,627 | 3,594 | 3,487 | 3,350 | 3,477 | 3,492 | 3,416 | 3,194 | 3,213 | 3,034 | -1.07 |
| ENSMUSG00000022114 | Spry2 | sprouty RTK sig | protein_coding | 584 | 606 | 646 | 536 | 606 | 558 | 539 | 579 | 482 | 506 | 505 | 521 | -1.04 |
| ENSMUSG00000050721 | Plekho2 | pleckstrin hom | protein_coding | 630 | 719 | 782 | 634 | 623 | 638 | 591 | 630 | 571 | 605 | 591 | 513 | -1.11 |
| ENSMUSG00000002580 | Mien1 | migration and i | protein_coding | 601 | 644 | 677 | 513 | 574 | 534 | 575 | 577 | 519 | 509 | 524 | 462 | -1.08 |
| ENSMUSG00000036278 | Macrod1 | mono-ADP ribo | protein_coding | 291 | 288 | 313 | 263 | 275 | 254 | 266 | 276 | 229 | 247 | 238 | 210 | -1.07 |
| ENSMUSG00000038489 | Polr2l | polymerase (RN | protein_coding | 414 | 404 | 415 | 373 | 363 | 329 | 397 | 399 | 339 | 330 | 360 | 301 | -1.08 |
| ENSMUSG00000020091 | Eif4ebp2 | eukaryotic tran | protein_coding | 2,214 | 2,289 | 2,396 | 2,054 | 2,055 | 1,836 | 2,136 | 2,062 | 1,852 | 1,953 | 2,102 | 1,756 | -1.11 |
| ENSMUSG00000066232 | Ipo7 | importin 7 | protein_coding | 4,331 | 4,408 | 4,466 | 4,331 | 4,252 | 4,057 | 4,208 | 4,221 | 4,116 | 4,069 | 4,195 | 3,942 | -1.05 |
| ENSMUSG00000019528 | Gyg | glycogenin | protein_coding | 735 | 762 | 760 | 686 | 687 | 650 | 731 | 706 | 641 | 647 | 681 | 638 | -1.06 |
| ENSMUSG00000020482 | Ccdc117 | coiled-coil dom | protein_coding | 568 | 664 | 654 | 531 | 492 | 500 | 574 | 555 | 451 | 449 | 491 | 435 | -1.14 |
| ENSMUSG00000071652 | Ints5 | integrator com | protein_coding | 386 | 423 | 497 | 360 | 348 | 347 | 400 | 397 | 345 | 338 | 368 | 323 | -1.11 |
| ENSMUSG00000073139 | Tmem185a | transmembran | protein_coding | 1,091 | 1,190 | 1,296 | 1,015 | 1,071 | 1,019 | 1,150 | 1,070 | 973 | 982 | 1,056 | 981 | -1.06 |
| ENSMUSG00000049521 | Cdc42ep1 | CDC42 effector | protein_coding | 850 | 858 | 771 | 718 | 718 | 672 | 684 | 675 | 676 | 694 | 627 | 674 | -1.16 |
| ENSMUSG00000000088 | Cox5a | cytochrome c o | protein_coding | 2,177 | 2,339 | 2,128 | 1,828 | 1,903 | 1,715 | 1,784 | 1,938 | 1,845 | 1,844 | 1,657 | 1,653 | -1.15 |
| ENSMUSG00000034216 | Vps18 | VPS18 CORVET | protein_coding | 1,549 | 1,614 | 1,545 | 1,348 | 1,369 | 1,251 | 1,340 | 1,381 | 1,267 | 1,307 | 1,255 | 1,277 | -1.13 |
| ENSMUSG00000050751 | Pgbd5 | piggyBac trans | protein_coding | 1,201 | 1,239 | 1,212 | 1,046 | 1,089 | 1,098 | 1,060 | 1,063 | 1,023 | 1,146 | 1,029 | 988 | -1.09 |
| ENSMUSG00000008855 | Hdac5 | histone deacet | protein_coding | 4,084 | 4,067 | 4,014 | 3,455 | 3,578 | 3,559 | 3,336 | 3,584 | 3,340 | 3,597 | 3,328 | 3,473 | -1.11 |
| ENSMUSG00000041199 | Rpusd1 | RNA pseudouri | protein_coding | 919 | 890 | 924 | 764 | 814 | 778 | 783 | 783 | 721 | 792 | 719 | 728 | -1.11 |
| ENSMUSG00000020456 | Ogdh | oxoglutarate (a | protein_coding | 8,824 | 8,763 | 8,696 | 8,004 | 8,104 | 7,894 | 8,008 | 8,137 | 7,789 | 8,119 | 7,929 | 7,874 | -1.07 |

| | | | | | | | | | | | | | | | | | | | | | | | | | | | | | | | | | | | | | | | | | |
|---|---|---|---|---|---|---|---|---|---|---|---|---|---|---|---|---|---|---|---|---|---|---|---|---|---|---|---|---|---|---|---|---|---|---|---|---|---|---|---|---|---|
| ENSMUSG00000027602 | Map1lc3a | microtubule-as | protein_coding | 2,227 | 2,194 | 2,322 | 1,867 | 1,865 | 1,757 | 1,804 | 1,892 | 1,699 | 1,956 | 1,731 | 1,708 | -1.18 | 0.00292 | 0.0782 | | -1.21 | 0.000347 | 0.00846 | down | -1.03 | 0.563 | 1 | | 7 | 2397 | 1.43056425 | 1.28895632 | 1.81937409 | -0.20341168 | -0.21357165 | -0.75867924 | -0.5209237 | -0.08146723 | -1.06861251 | 0.22657431 | -0.89939045 | -1.01941252 |
| ENSMUSG00000013646 | Sh3bp5l | SH3 binding do | protein_coding | 929 | 1,018 | 1,021 | 892 | 856 | 867 | 859 | 932 | 849 | 838 | 856 | 829 | -1.1 | 0.0374 | 0.237 | | -1.14 | 0.00176 | 0.0236 | down | -1.04 | 0.313 | 1 | | 7 | 2398 | 0.54602915 | 1.82986083 | 1.86352733 | -0.03243914 | -0.59721084 | -0.42708939 | -0.5453446 | 0.58266132 | -0.710085 | -0.88563964 | -0.59303546 | -1.03123455 |
| ENSMUSG00000014349 | Ube2z | ubiquitin-conju | protein_coding | 2,566 | 2,640 | 2,691 | 2,338 | 2,297 | 2,340 | 2,323 | 2,451 | 2,361 | 2,282 | 2,302 | 2,216 | -1.09 | 0.0225 | 0.192 | | -1.12 | 0.00203 | 0.0259 | down | -1.03 | 0.438 | 1 | | 7 | 2399 | 1.10062547 | 1.56533022 | 1.87223547 | -0.39547336 | -0.68020132 | -0.38748453 | -0.50006447 | 0.35984465 | -0.2412956 | -0.78547929 | -0.64675248 | -1.26128476 |
| ENSMUSG00000029405 | G3bp2 | GTPase activati | protein_coding | 12,783 | 12,876 | 13,178 | 11,924 | 11,891 | 11,840 | 11,841 | 12,184 | 11,719 | 11,659 | 11,849 | 11,313 | -1.06 | 0.0349 | 0.232 | | -1.09 | 0.0025 | 0.0297 | down | -1.03 | 0.365 | 1 | | 7 | 2400 | 1.24382675 | 1.40259961 | 1.91050349 | -0.27896353 | -0.33935266 | -0.43259386 | -0.43076343 | 0.1939595 | -0.65731644 | -0.76935985 | -0.41588157 | -1.42665801 |
| ENSMUSG00000004383 | Large1 | LARGE xylosyl- | protein_coding | 1,974 | 1,999 | 1,974 | 1,776 | 1,736 | 1,730 | 1,816 | 1,909 | 1,715 | 1,729 | 1,805 | 1,687 | -1.07 | 0.0588 | 0.292 | | -1.11 | 0.00299 | 0.0333 | down | -1.04 | 0.292 | 1 | | 7 | 2401 | 1.34462936 | 1.54614062 | 1.337321 | -0.3770734 | -0.74708753 | -0.80374411 | -0.01258119 | 0.79867741 | -0.94432961 | -0.81487026 | -0.11935581 | -1.20772647 |
| ENSMUSG00000035764 | Fbxo45 | F-box protein 4 | protein_coding | 1,493 | 1,462 | 1,457 | 1,318 | 1,332 | 1,324 | 1,325 | 1,382 | 1,250 | 1,267 | 1,327 | 1,271 | -1.07 | 0.0755 | 0.327 | | -1.12 | 0.00145 | 0.021 | down | -1.05 | 0.168 | 1 | | 7 | 2402 | 1.72686819 | 1.37349696 | 1.31628165 | -0.39046195 | -0.21618459 | -0.30855174 | -0.30035811 | 0.41674614 | -1.27733343 | -1.05700299 | -0.27567644 | -1.00782371 |
| ENSMUSG00000017639 | Rab11fip4 | RAB11 family in | protein_coding | 8,550 | 8,694 | 9,027 | 8,135 | 7,919 | 7,661 | 7,909 | 8,121 | 7,650 | 7,944 | 7,661 | 7,885 | -1.09 | 0.00952 | 0.128 | | -1.1 | 0.00214 | 0.027 | down | -1.02 | 0.642 | 1 | | 7 | 2403 | 1.0459458 | 1.35904431 | 2.0647728 | 0.11387606 | -0.38957909 | -1.01111424 | -0.41480043 | 0.08230071 | -1.03790692 | -0.33195302 | -1.01073132 | -0.46985467 |
| ENSMUSG00000042604 | Kcna4 | potassium volta | protein_coding | 707 | 730 | 769 | 658 | 582 | 592 | 581 | 627 | 550 | 602 | 542 | 590 | -1.2 | 0.000368 | 0.0363 | down | -1.25 | 0.00000642 | 0.000749 | down | -1.04 | 0.39 | 1 | | 7 | 2404 | 1.1081475 | 1.39797754 | 1.87029102 | 0.46723294 | -0.61434172 | -0.46394169 | -0.63660772 | 0.0382929 | -1.11072247 | -0.32878106 | -1.22981204 | -0.4977352 |
| ENSMUSG00000003721 | Insig2 | insulin induced | protein_coding | 1,325 | 1,513 | 1,456 | 1,242 | 1,066 | 1,079 | 1,134 | 1,152 | 1,074 | 1,152 | 1,033 | 991 | -1.25 | 0.0000429 | 0.0204 | down | -1.3 | 0.000000815 | 0.000231 | down | -1.04 | 0.43 | 1 | | 7 | 2405 | 0.8867831 | 1.88987569 | 1.59957232 | 0.40469393 | -0.71583002 | -0.63293042 | -0.26988845 | -0.15404841 | -0.66552201 | -0.15241085 | -0.94490064 | -1.24539423 |
| ENSMUSG00000046791 | Riox1 | ribosomal oxyg | protein_coding | 227 | 242 | 242 | 200 | 168 | 172 | 185 | 205 | 157 | 174 | 170 | 161 | -1.24 | 0.00282 | 0.0767 | | -1.37 | 0.00000472 | 0.000638 | down | -1.11 | 0.142 | 1 | | 7 | 2406 | 1.15324493 | 1.58365645 | 1.58666715 | 0.31438377 | -0.76652345 | -0.65408524 | -0.20255492 | 0.47071525 | -1.17500104 | -0.58027535 | -0.70151224 | -1.02871531 |
| ENSMUSG00000072437 | Nanos1 | nanos C2HC-typ | protein_coding | 1,167 | 1,226 | 1,229 | 1,039 | 991 | 994 | 1,032 | 1,049 | 973 | 995 | 978 | 988 | -1.15 | 0.00629 | 0.106 | | -1.18 | 0.000534 | 0.0113 | down | -1.03 | 0.493 | 1 | | 7 | 2407 | 1.19303435 | 1.76037442 | 1.78847278 | -0.1371723 | -0.67491139 | -0.63629116 | -0.22025449 | -0.03167136 | -0.8815811 | -0.63003235 | -0.81923673 | -0.71073066 |
| ENSMUSG00000041040 | Fam117b | family with seq | protein_coding | 2,868 | 2,894 | 3,052 | 2,841 | 2,775 | 2,601 | 2,701 | 2,742 | 2,650 | 2,746 | 2,685 | 2,556 | -1.08 | 0.0152 | 0.161 | | -1.1 | 0.00229 | 0.0281 | down | -1.02 | 0.559 | 1 | | 7 | 2408 | 0.80145615 | 0.98233843 | 2.06097083 | 0.61056009 | 0.13321111 | -1.16616038 | -0.40663077 | -0.10607213 | -0.79240383 | -0.07228023 | -0.52555351 | -1.51943575 |
| ENSMUSG00000049657 | Zbtb5 | zinc finger and | protein_coding | 440 | 444 | 446 | 418 | 393 | 364 | 391 | 412 | 378 | 379 | 381 | 367 | -1.12 | 0.0295 | 0.216 | | -1.16 | 0.00206 | 0.0263 | down | -1.04 | 0.414 | 1 | | 7 | 2409 | 1.29889884 | 1.42055778 | 1.46016518 | 0.57453455 | -0.25956449 | -1.26640249 | -0.30272445 | 0.39658057 | -0.7634435 | -0.72886447 | -0.66529035 | -1.16444717 |
| ENSMUSG00000055943 | Emc7 | ER membrane p | protein_coding | 3,564 | 3,537 | 3,671 | 3,379 | 3,179 | 3,195 | 3,254 | 3,418 | 3,211 | 3,288 | 3,213 | 3,084 | -1.08 | 0.0154 | 0.162 | | -1.11 | 0.00221 | 0.0276 | down | -1.02 | 0.541 | 1 | | 7 | 2410 | 1.27584056 | 1.13304122 | 1.82815665 | 0.28092755 | -0.85322143 | -0.75970432 | -0.42238239 | 0.49173238 | -0.6699125 | -0.22970675 | -0.65781919 | -1.41695177 |
| ENSMUSG00000007944 | Ttc9b | tetratricopeptic | protein_coding | 563 | 596 | 573 | 483 | 523 | 510 | 629 | 535 | 460 | 445 | 467 | 439 | -1.01 | 0.898 | 0.968 | | -1.22 | 0.00132 | 0.0199 | down | -1.21 | 0.00183 | 1 | | 8 | 2411 | 0.74072648 | 1.23271877 | 0.8849265 | -0.55139812 | 0.11555068 | -0.1030013 | 1.70182867 | 0.30133264 | -0.95674412 | -1.21975186 | -0.82071115 | -1.3254772 |
| ENSMUSG00000020376 | Rnf130 | ring finger prot | protein_coding | 4,638 | 4,864 | 4,905 | 4,806 | 4,548 | 4,546 | 5,107 | 4,835 | 4,460 | 4,626 | 4,217 | 4,258 | -1.01 | 0.781 | 0.923 | | -1.09 | 0.00571 | 0.0496 | down | -1.08 | 0.0126 | 1 | | 8 | 2412 | -0.02218497 | 0.80600751 | 0.95307098 | 0.59482099 | -0.36274504 | -0.37074875 | 1.65665656 | 0.70105909 | -0.70366091 | -0.0683685 | -1.67628995 | -1.50761701 |
| ENSMUSG00000023367 | Tmem176a | transmembrane | protein_coding | 850 | 870 | 707 | 701 | 805 | 801 | 919 | 666 | 694 | 697 | 594 | 606 | 1.02 | 0.767 | 0.917 | | -1.2 | 0.00277 | 0.0316 | down | -1.23 | 0.000865 | 1 | | 8 | 2413 | 1.02475263 | 1.19731081 | -0.29871402 | -0.35230919 | 0.63293154 | 0.59928794 | 1.60081326 | -0.71097341 | -0.42665103 | -0.39297211 | -1.50833027 | -1.36514615 |
| ENSMUSG00000002602 | Axl | AXL receptor ty | protein_coding | 1,506 | 1,631 | 1,430 | 1,594 | 1,567 | 1,571 | 1,581 | 1,589 | 1,471 | 1,400 | 1,135 | 1,131 | 1.02 | 0.7 | 0.887 | | -1.2 | 0.00329 | 0.0354 | down | -1.23 | 0.00085 | 1 | | 8 | 2414 | 0.26031562 | 0.90027077 | -0.15797038 | 0.71414299 | 0.57673258 | 0.60001233 | 0.64777624 | 0.68835828 | 0.06727685 | -0.32434605 | -1.97348922 | -1.99907999 |
| ENSMUSG00000043257 | Pigv | phosphatidylinc | protein_coding | 313 | 398 | 281 | 338 | 350 | 321 | 354 | 310 | 290 | 254 | 214 | 224 | 1 | 0.961 | 0.989 | | -1.35 | 0.00119 | 0.0186 | down | -1.36 | 0.000941 | 1 | | 8 | 2415 | 0.22008828 | 1.58166373 | -0.37102394 | 0.64927536 | 0.83587965 | 0.35919127 | 0.91392021 | 0.16750817 | -0.20186518 | -0.88203373 | -1.75023297 | -1.52237083 |
| ENSMUSG00000109523 | Gdf1 | growth differen | protein_coding | 52 | 0 | 95 | 0 | 0 | 83 | 50 | 0 | 0 | 0 | 0 | 0 | -1.11 | 0.947 | | | -240 | 0.001 | 0.0168 | down | -217 | 0.00124 | 1 | | 8 | 2416 | 1.08798336 | -0.66807575 | 1.67278438 | -0.66807575 | -0.66807575 | 1.52577034 | 1.05806796 | -0.66807575 | -0.66807575 | -0.66807575 | -0.66807575 | -0.66807575 |
| ENSMUSG00000058135 | Gstm1 | glutathione S-tr | protein_coding | 5,333 | 4,857 | 5,477 | 4,720 | 5,521 | 4,709 | 5,041 | 4,862 | 4,430 | 4,641 | 4,567 | 4,353 | -1.01 | 0.746 | 0.909 | | -1.13 | 0.00116 | 0.0184 | down | -1.12 | 0.00337 | 1 | | 8 | 2417 | 1.16725101 | -0.01360422 | 1.50427233 | -0.37360402 | 1.60729688 | -0.4051629 | 0.45533823 | -0.00069987 | -1.1715538 | -0.58812179 | -0.78877317 | -1.39263868 |
| ENSMUSG00000028029 | Aimp1 | aminoacyl tRNA | protein_coding | 919 | 849 | 897 | 818 | 829 | 879 | 895 | 830 | 795 | 812 | 747 | 770 | -1.02 | 0.701 | 0.888 | | -1.11 | 0.00573 | 0.0497 | down | -1.1 | 0.0158 | 1 | | 8 | 2418 | 1.51148955 | 0.24974575 | 1.11715702 | -0.33299211 | -0.11154356 | 0.7999878 | 1.09505192 | -0.09760808 | -0.77933806 | -0.44772218 | -1.73928304 | -1.26494502 |
| ENSMUSG00000090946 | Ccdc71l | coiled-coil dom | protein_coding | 562 | 454 | 551 | 468 | 500 | 453 | 554 | 478 | 437 | 424 | 402 | 415 | -1.03 | 0.654 | 0.865 | | -1.21 | 0.000602 | 0.0121 | down | -1.18 | 0.00256 | 1 | | 8 | 2419 | 1.52231487 | -0.3439562 | 1.35490474 | -0.08132081 | 0.49715743 | -0.36040812 | 1.39087503 | 0.09454031 | -0.67847838 | -0.92125044 | -1.36175103 | -1.1126274 |

**Supplemental table S4**

#### Mid aged hypothalami KEGG pathways

| | Overlapping genes | Percentage of changed genes | Adjusted P-value |
|---|---|---|---|
| Parkinson disease | 37/249 | 14.86 | 1.5991978964679698E-12 |
| Prion disease | 36/273 | 13.19 | 7.590195460486483E-11 |
| Oxidative phosphorylation | 24/133 | 18.05 | 4.850765470128784E-10 |
| Huntington disease | 36/306 | 11.76 | 1.1257268903987683E-9 |
| Pathways of neurodegeneration | 44/475 | 9.26 | 1.519335649516149E-8 |
| Amyotrophic lateral sclerosis | 37/364 | 10.16 | 2.6925896140360836E-8 |
| Non-alcoholic fatty liver disease | 23/155 | 14.84 | 3.3836591015647834E-8 |
| Ribosome | 23/158 | 14.56 | 4.3529759198135816E-8 |
| Diabetic cardiomyopathy | 25/203 | 12.32 | 2.5453131731948394E-7 |
| Alzheimer disease | 34/369 | 9.21 | 9.611151192276847E-7 |
| Thermogenesis | 24/232 | 10.34 | 1.1429371668208388E-5 |
| Retrograde endocannabinoid signaling | 18/148 | 12.16 | 2.8035964895516726E-5 |
| Coronavirus disease | 23/232 | 9.91 | 3.511793871377748E-5 |
| Vibrio cholerae infection | 10/50 | 20.00 | 6.572758668345648E-5 |
| Salmonella infection | 23/249 | 9.24 | 1.0090689089411308E-4 |
| Protein processing in endoplasmic reticulum | 17/171 | 9.94 | 6.13676917896875E-4 |
| Glycolysis / Gluconeogenesis | 10/67 | 14.93 | 8.022505842935701E-4 |
| Phagosome | 15/152 | 9.87 | 0.001683424 |
| Vasopressin-regulated water reabsorption | 07/44 | 15.91 | 0.006520571 |
| Endocytosis | 19/252 | 7.54 | 0.006899784 |
| Synaptic vesicle cycle | 09/78 | 11.54 | 0.010981511546577808 |
| RNA transport | 15/186 | 8.06 | 0.011901998670108155 |
| N-Glycan biosynthesis | 07/50 | 14.00 | 0.011901998670108155 |
| Collecting duct acid secretion | 05/27 | 18.52 | 0.016505806 |
| Autophagy | 12/137 | 8.76 | 0.016505806 |
| Epithelial cell signaling in Helicobacter pylori infection | 08/70 | 11.43 | 0.018144784424553764 |
| HIF-1 signaling pathway | 10/109 | 9.17 | 0.026280416365842274 |
| Bacterial invasion of epithelial cells | 08/77 | 10.39 | 0.030925364 |
| Proteasome | 06/46 | 13.04 | 0.03111624 |
| mTOR signaling pathway | 12/154 | 7.79 | 0.036392459 |
| Cortisol synthesis and secretion | 07/65 | 10.77 | 0.040816803 |
| Longevity regulating pathway | 9/102 | 8.82 | 0.046177619873792874 |
| Mitophagy | 07/68 | 10.29 | 0.049202567 |

**Young hypothalami KEGG pathways**

| | Overlapping genes | Percentage of changed genes | Adjusted P-value |
|---|---|---|---|
| Insulin signaling pathway | 13/137 | 9.49 | 2.3218886596153645E-4 |
| Neurotrophin signaling pathway | 12/119 | 10.08 | 2.3218886596153645E-4 |
| Dopaminergic synapse | 12/132 | 9.09 | 4.5998615046183625E-4 |
| Lipid and atherosclerosis | 15/215 | 6.98 | 6.441239928866919E-4 |
| Amphetamine addiction | 08/69 | 11.59 | 0.00175691 |
| Insulin resistance | 9/108 | 8.33 | 0.006607123 |
| Estrogen signaling pathway | 10/137 | 7.30 | 0.007172165 |
| Kaposi sarcoma-associated herpesvirus infection | 12/193 | 6.22 | 0.007172165 |
| Human cytomegalovirus infection | 13/225 | 5.78 | 0.007297474 |
| Signaling pathways regulating pluripotency of stem cell | 10/143 | 6.99 | 0.007378476 |
| Growth hormone synthesis, secretion and action | 9/119 | 7.56 | 0.007473632 |
| Axon guidance | 11/182 | 6.04 | 0.010499673 |
| Longevity regulating pathway | 8/102 | 7.84 | 0.010499673 |
| Apelin signaling pathway | 9/137 | 6.57 | 0.016007478326142174 |
| Colorectal cancer | 07/86 | 8.14 | 0.016007478326142174 |
| Tight junction | 10/169 | 5.92 | 0.016971707441899155 |
| Focal adhesion | 11/201 | 5.47 | 0.017403907 |
| Diabetic cardiomyopathy | 11/203 | 5.42 | 0.017558078 |
| Long-term potentiation | 06/67 | 8.96 | 0.017558078 |
| Prion disease | 13/273 | 4.76 | 0.019736277322445303 |
| Circadian entrainment | 07/97 | 7.22 | 0.021519642230805252 |
| Prostate cancer | 07/97 | 7.22 | 0.021519642230805252 |
| Oxytocin signaling pathway | 9/154 | 5.84 | 0.021519642230805252 |
| cAMP signaling pathway | 11/216 | 5.09 | 0.021519642230805252 |
| Non-alcoholic fatty liver disease | 9/155 | 5.81 | 0.021519642230805252 |
| Regulation of actin cytoskeleton | 11/218 | 5.05 | 0.021604731326235044 |
| Relaxin signaling pathway | 8/129 | 6.20 | 0.02274885 |
| Chronic myeloid leukemia | 06/76 | 7.89 | 0.02274885 |
| C-type lectin receptor signaling pathway | 7/104 | 6.73 | 0.023353882263573496 |
| T cell receptor signaling pathway | 7/104 | 6.73 | 0.023353882263573496 |
| Hepatitis B | 9/162 | 5.56 | 0.023353882263573496 |
| Hippo signaling pathway | 9/163 | 5.52 | 0.023569705 |
| Parathyroid hormone synthesis, secretion and action | 7/106 | 6.60 | 0.024188684211157985 |
| Alzheimer disease | 15/369 | 4.06504065 | 0.024233355988527307 |
| cGMP-PKG signaling pathway | 9/167 | 5.389221557 | 0.024233355988527307 |
| Autophagy | 8/137 | 5.839416058 | 0.024233355988527307 |
| Yersinia infection | 8/137 | 5.839416058 | 0.024233355988527307 |
| Fluid shear stress and atherosclerosis | 8/139 | 5.755395683 | 0.025757209662296634 |
| Aldosterone-regulated sodium reabsorption | 04/37 | 10.81081081 | 0.027408099 |
| ErbB signaling pathway | 06/85 | 7.058823529 | 0.027686810428600727 |
| Cholinergic synapse | 7/113 | 6.194690265 | 0.027686810428600727 |
| Leukocyte transendothelial migration | 7/114 | 6.140350877 | 0.028354276914465768 |
| Human immunodeficiency virus 1 infection | 10/212 | 4.716981132 | 0.032405794 |
| Sphingolipid signaling pathway | 7/119 | 5.882352941 | 0.034110154 |
| Pathways of neurodegeneration | 17/475 | 3.578947368 | 0.036686998 |
| GnRH signaling pathway | 06/93 | 6.451612903 | 0.036829921 |
| Alcoholism | 9/186 | 4.838709677 | 0.036829921 |
| IL-17 signaling pathway | 06/94 | 6.382978723 | 0.036829921 |
| TGF-beta signaling pathway | 06/94 | 6.382978723 | 0.036829921 |
| Renal cell carcinoma | 05/69 | 7.246376812 | 0.040668074429793644 |
| Phosphatidylinositol signaling system | 06/97 | 6.18556701 | 0.040668074429793644 |
| Chemokine signaling pathway | 9/192 | 4.6875 | 0.040668074429793644 |
| Prolactin signaling pathway | 05/70 | 7.142857143 | 0.040668074429793644 |
| Aldosterone synthesis and secretion | 06/98 | 6.12244898 | 0.040668074429793644 |
| Pathways in cancer | 18/531 | 3.389830508 | 0.040964442 |
| AGE-RAGE signaling pathway in diabetic complications | 6/100 | 6 | 0.043088308 |
| Melanogenesis | 6/101 | 5.940594059 | 0.043886879 |
| Ras signaling pathway | 10/232 | 4.310344828 | 0.043886879 |
| Vascular smooth muscle contraction | 7/133 | 5.263157895 | 0.045541846596865616 |
| Wnt signaling pathway | 8/166 | 4.819277108 | 0.045807181 |
| Glioma | 05/75 | 6.666666667 | 0.046248289831599294 |
| Thyroid hormone synthesis | 05/75 | 6.666666667 | 0.046248289831599294 |
